# Genomic islands of differentiation in a rapid avian radiation have been driven by recent selective sweeps

**DOI:** 10.1101/2020.03.07.977694

**Authors:** Hussein A. Hejase, Ayelet Salman-Minkov, Leonardo Campagna, Melissa J. Hubisz, Irby J. Lovette, Ilan Gronau, Adam Siepel

## Abstract

Numerous studies of emerging species have identified genomic “islands” of elevated differentiation against a background of relative homogeneity. The causes of these islands remain unclear, however, with some signs pointing toward “speciation genes” that locally restrict gene flow and others suggesting selective sweeps that have occurred within nascent species after speciation. Here, we examine this question through the lens of genome sequence data for five species of southern capuchino seedeaters, finch-like birds from South America that have undergone a species radiation during the last ∼50,000 generations. By applying newly developed statistical methods for ancestral recombination graph inference and machine-learning methods for the prediction of selective sweeps, we show that previously identified islands of differentiation in these birds appear to be generally associated with relatively recent, species-specific selective sweeps, most of which are predicted to be “soft” sweeps acting on standing genetic variation. Many of these sweeps coincide with genes associated with melanin-based variation in plumage, suggesting a prominent role for sexual selection. At the same time, a few loci also exhibit indications of possible selection against gene flow. These observations shed new light on the complex manner in which natural selection shapes genome sequences during speciation.

**Significance Statement:** Genome-wide scans can identify differentiated loci between species that may have promoted speciation. So-called “islands of differentiation” have generally been identified and characterized using standard population genetic summary statistics (e.g., *F*_*ST*_ and *d*_*xy*_), which are limited in distinguishing among possible causes of differentiation, such as selection against gene flow and selective sweeps. We introduce a powerful strategy for analyzing such islands, combining new methods for inferring the full Ancestral Recombination Graph and machine learning methods for identifying selective sweeps. We applied our methods to genomic sequences from closely related southern capuchino seedeaters (Neotropical tanagers) and found signatures of recent selective sweeps around pigmentation genes, including many “soft” sweeps that acted on standing variation.

## Introduction

The question of how new species arise is one of the oldest and thorniest puzzles in evolutionary biology, having occupied investigators at least since Darwin and Wallace^1–3^. A key component of the neodarwinian synthesis of the early 20th century was to formulate a genetic basis for the process of speciation^1,4,5^. By the mid-1900s, theories of speciation had come to rely heavily on the notion of allopatry, or geographical or ecological barriers to interbreeding^6^. However, it has since become clear that separate species may also emerge in the absence of such barriers, and there are now numerous well-supported cases of speciation with gene flow^7–14^.

In recent decades, inexpensive, large-scale DNA sequencing has allowed genetic studies of speciation to be extended from traditional laboratory model systems to natural populations. Particularly useful are natural systems that consist of populations in the early stages of genetic separation—that is, groups of organisms “caught in the act” of speciation^1^. These systems allow genomic regions with relevant differences to stand out clearly against a background of relatively low genetic differentiation. Examples include *Heliconius* butterflies^15,16^, *Anopheles* mosquitos^17,18^, pea aphids^19^, stick insects^20^, sunflowers^21^, monkeyflowers^22^, house mice^23,24^, threespine stickleback^25^ and cichlid^26^ fish, and various birds, including carrion and hooded crows^27–29^, flycatchers^30^, and blue- and golden-winged warblers^31^. In addition to enabling investigation of the genetic architecture of reproductive barriers, these systems allow many other questions to be addressed, such as what is the timeline of speciation, and what are the roles of chromosomal rearrangements and sex chromosomes^1^? In some cases, they also allow identification of particular mutations underlying important differences between incipient species.

A number of such studies have focused, in particular, on the intriguing observation that recently separated species often exhibit local genomic regions of pronounced interspecies differentiation against a background of relative homogeneity—a phenomenon dubbed “islands of differentiation” or, sometimes, “islands of speciation”^1,17^. Early on, these islands were widely believed to reflect “speciation genes” (sometimes called “barrier loci”) that contribute in some way to reduced gene flow, presumably through inviability, sterility, or reduced fitness of hybrids^17,32^. Later, however, it was shown that a similar local elevation of genetic differentiation could be generated instead by reductions of genetic diversity within one or both nascent species during and following speciation, stemming from selective sweeps and possibly background selection^33–35^. In other words, the observed “islands” may not reflect differential gene flow, but instead, differential influences of selection. This ambiguity follows from the widespread use of relative measures of genetic differentiation such as *F*_ST_, which can be elevated either by increases in interspecies divergence or decreases in intraspecies variation. Some recent studies have considered competing hypotheses for the origins of islands of differentiation and found mixed evidence regarding their primary causes^28,29,36^.

Recently, we collected whole-genome sequence data for another natural system: southern capuchino seedeaters of the genus *Sporophila*, a group of passerine birds native to South America. These birds are similar to Darwin’s finches (also members of the family Thraupidae) in that they appear to have recently separated into numerous distinct and largely genetically separate species. The southern capuchino species, however, are morphologically and ecologically indistinct and differ primarily in male plumage, suggesting that sexual selection may have been important in driving speciation^37–39^. These species have adjacent and, in many cases, overlapping ranges^40^, and can be syntopic at breeding sites. In addition, they include many pairs with similar levels of genetic differentiation, allowing for numerous quasi-independent pairwise comparisons.

Our data set includes nine species of southern capuchinos, eight of which are estimated to have emerged <50,000 generations ago from a large, highly polymorphic ancestral species^41^. We focus here on the five of these eight species that were most broadly sampled (12 diploid individuals per species), including *Sporophila pileata* (pil), *S. melanogaster* (mel), *S. nigrorufa* (nig), *S. palustris* (pal), and *S. hypoxantha* (hypox). The combination of a recent radiation and a large ancestral population for these species leads to modest levels of genetic differentiation and a great deal of shared polymorphism, reflecting widespread incomplete lineage sorting. Nevertheless, these genomes include numerous striking islands of differentiation, as defined by peaks in pairwise *F*_ST_, and these islands are strongly enriched for genes in the melanogenesis pathway, suggesting an association with melanin-based variation in plumage^40^. Campagna *et al*. focused in particular on 25 peaks with elevated *F*_ST_, which represent candidate targets of selection during or after speciation. Many of these islands appear recurrently across different pairwise comparisons and 10 of the 25 peaks are located on the sex-linked *Z* chromosome.

In this paper, we re-examine these islands of differentiation in the southern capuchinos. Rather than focusing on the use of summary statistics in windows across the genome, as in most previous studies, we base our analysis on the underlying genealogies that precisely describe the relationships among the sequenced genomes. In particular, we make use of methods we recently developed for sequence-based statistical inference of the ancestral recombination graph (ARG), which describes both genealogical relationships and changes to those relationships along the genome due to historical recombination events^42,43^. This approach enables us to delve more deeply into the evolutionary processes underlying these islands, and distinguish among their possible causes. Furthermore, we combine ARG inference with the latest machine-learning methods^44^ to identify selective sweeps of various ages and types, and systematically compare these predictions with the previously identified islands of differentiation. As we discuss in detail below, we find multiple lines of evidence for a strong relationship between selective sweeps—particularly recent, species-specific “soft” sweeps—and the observed islands. By contrast, we find much less evidence for an influence from selection against gene flow, with a few notable exceptions. Altogether, our study provides new insights into the processes leading to genetic differentiation between species in a powerful model system for rapid speciation.

## Results

### Genealogical Patterns in *F*_ST_ Peaks

We began by reanalyzing the 25 *F*_ST_ peaks (**Supplementary Table S1 and Supplementary Figure S1**) identified by Campagna et al.^40^, with the goal of gaining further insight into their evolutionary causes. As noted above, these peaks are presumed to reflect the speciation process in some manner, but the previous reliance on summary statistics made it difficult to distinguish among the specific evolutionary processes that may have generated them. We were particularly interested in using ARG-based methods to distinguish between a scenario where *F*_ST_ peaks were driven by selection against gene flow that was established early in the speciation process, and one where they were driven by more recent selective sweeps (**Figure 1**), allowing for the possibility that both scenarios could be at play (see **Discussion**).

**Figure 1:**
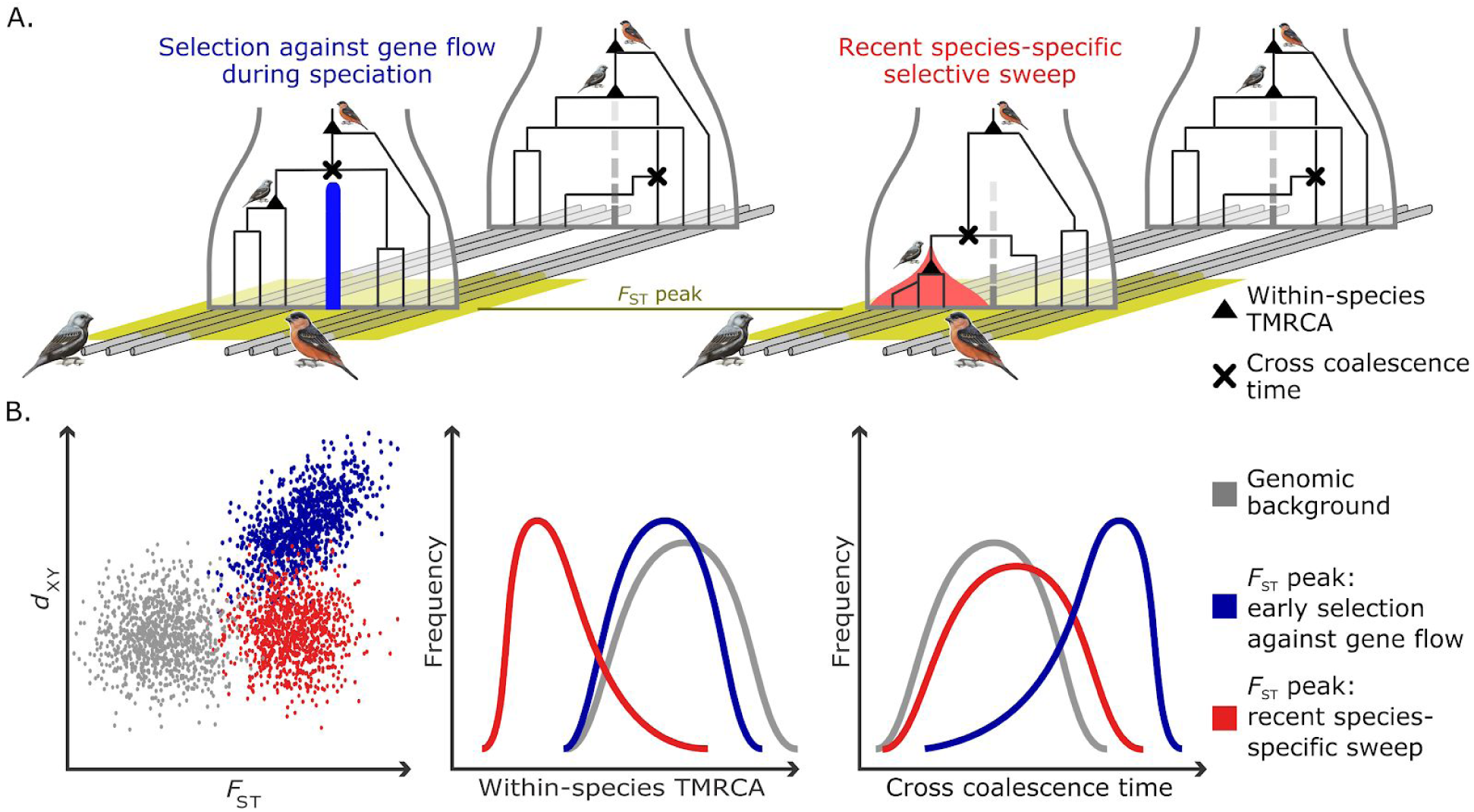
Two models for the formation of islands of differentiation (*F*_ST_ peaks): selection against gene flow during speciation vs. recent species-specific selective sweeps. (**A**) Representative genealogies for six individuals from two recently diverged species in an *F*_ST_ peak (*yellow*) and in a nearby neutrally evolving genomic region (*gray*). Under both models, neutral genealogies exhibit frequent incomplete lineage sorting (ILS) due to large ancestral population sizes and possible ongoing gene flow between species (*vertical dashed gray line*). ILS is reduced in the *F*_ST_ peaks either because the time back to the first cross-coalescence (marked by *X*) is elevated by selection against gene flow (*left*), or because the within-species time to most recent common ancestry (TMRCA; marked by *triangle*) is reduced by a selective sweep in one or both species (*right*). (**B**) Hypothetical distributions of various sequence-based and genealogy-based measures under the two models. Selection against gene flow (*blue*) will tend to induce a positive correlation between *F*_ST_ and *d*_XY_, whereas species-specific sweeps (*red*) will not (*left*). The within-species TMRCA is expected to be reduced by selective sweeps, but not substantially by selection against gene flow (*middle*). The cross-coalescence time is expected to be elevated by selection against gene flow but not substantially by species-specific sweeps (*right*).

We applied the *ARGweaver* program to the 60 individual genome sequences from five *Sporophila* species across 1.06 Gb of sequenced scaffolds (see **Supplementary Text**). This program jointly infers genealogies across all nucleotide sites, accounting for their correlation structure via an assumed Markovian recombination process. By efficiently pooling information across haplotypes, *ARGweaver* can obtain a fairly accurate reconstruction of the genealogy at each position^42^. We summarized the inferred genealogies both within the peaks and in flanking “background” regions using several features, including the expected time to most recent common ancestry (TMRCA) within each species—which should be reduced in peaks in the presence of local sweeps but unaffected by selection against gene flow between species during speciation—and the cross-coalescence time between species—which should be increased in peaks by selection against gene flow but unaffected by local sweeps (**Figure 1**). We also calculated the standard summary statistics *F*_ST_ and *d*_XY_ in a 10kb sliding window. Notice that the within-species TMRCA and the cross-coalescence time are related to the conventional summary statistics π_Within_ (a.k.a. π_W_) and *d*_XY_ (a.k.a. π_Between_, π_B_, or π_XY_), respectively, which have been previously used to distinguish among different contributing factors to *F*_ST_ peaks^33,36,45^. However, the ARG-based statistics are potentially more informative, by reflecting a comprehensive, model-based inference that considers correlated, fully resolved genealogies and the full distribution of coalescence times. Notably, these ARG-based statistics also provide a natural means for accommodating background selection and mutation rate variation (see **Supplementary Text** and **Discussion**).

We first compared the windowed *F*_ST_ and *d*_XY_ statistics within the *F*_ST_ peaks. If the *F*_ST_ peaks are driven by selection against gene flow, then these *F*_ST_ and *d*_XY_ estimates should be positively correlated, with larger values indicating deeper, and smaller values indicating shallower, cross-species coalescence (**Figure 1**)^33,36^. By contrast, if the *F*_ST_ peaks are driven by recent, species-specific sweeps, then no correlation is expected, because *F*_ST_ will be determined by the strength and ages of the sweeps, and *d*_XY_ will be largely consistent with the background. We found that the correlation between these statistics was quite low overall (Pearson’s *r* = 0.197; **Supplementary Figure S2**), more consistent with a “recent sweeps” model than an “early selection against gene flow” model.

Next we turned to the *ARGweaver*-inferred genealogies and compared the within-species TMRCAs in the *F*_ST_ peaks and the background regions. If the *F*_ST_ peaks are driven by selective sweeps, we would expect them to exhibit a reduced within-species TMRCA relative to the background, reflecting recent increases in the frequency of selected alleles and corresponding “bursts” of coalescence. Indeed, the inferred genealogies at several *F*_ST_ peaks, such as the one on scaffold 404 near the *SLC45A2* gene (**Figure 2A**), display this qualitative behavior, with dense clusters of recent coalescence events suggesting selective sweeps (see additional examples in **Supplementary Figure S3**). Accordingly, a measure of the within-species TMRCA that allows for partial sweeps, called TMRCAH (the TMRCA for the first 50% of all lineages from the species; see **Methods**), is depressed at these *F*_ST_ peaks (**Figure 2B**). At the same time, the cross-coalescence time does not tend to be elevated at these peaks, as would be expected if they were driven by selection against gene flow during speciation (**Figure 2C**).

**Figure 2:**
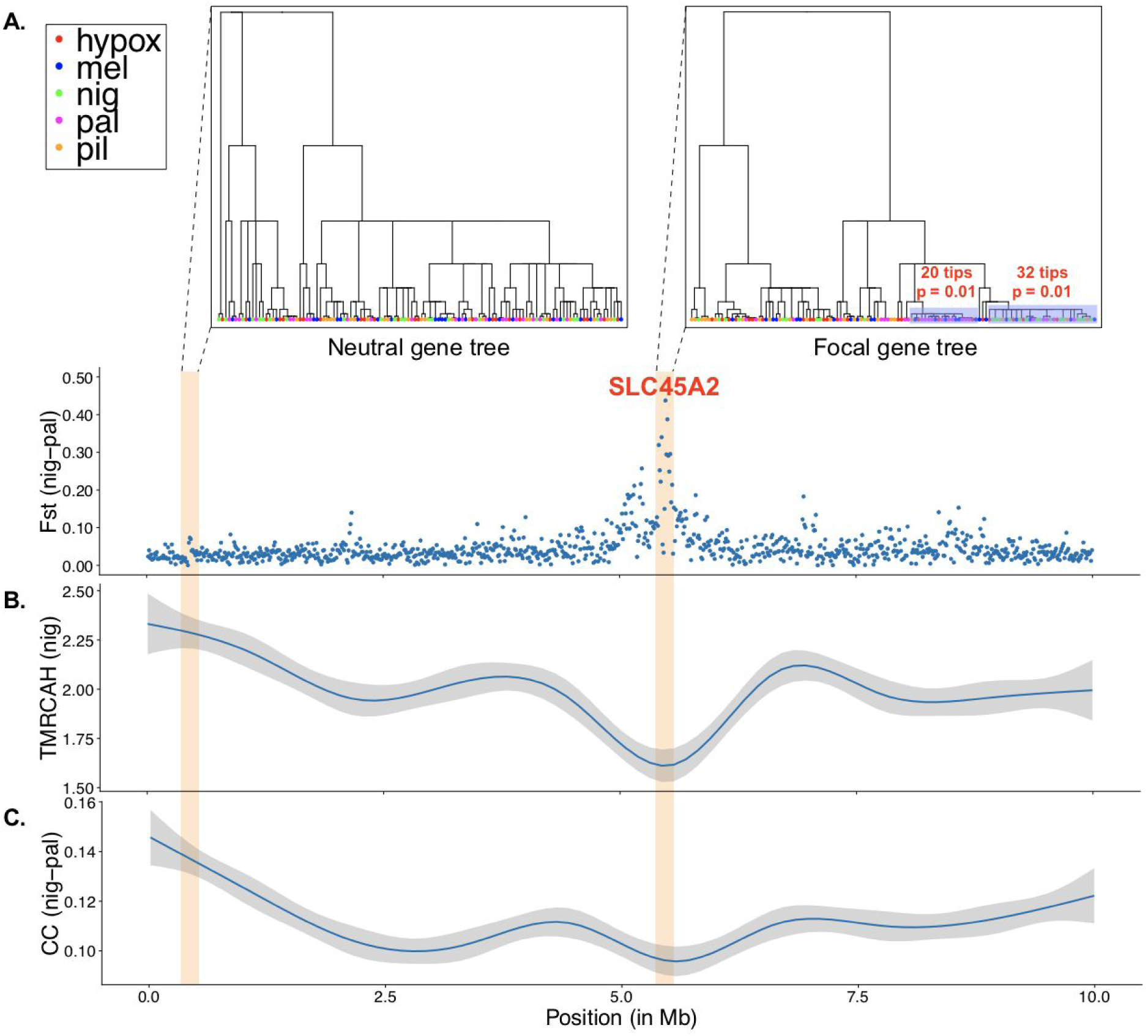
Comparison of inferred genealogies in an *F*_ST_ peak and in flanking neutral regions for the first 10 Mb of scaffold 404. (**A**) Average *F*_ST_ values in non-overlapping 10-kb windows between *S. nigrorufa* (nig) and *S. palustris* (pal), showing a pronounced peak near the pigmentation-related gene, *SLC45A2*. Local trees are shown within the peak (*top right*) and in a flanking neutral region (*top left*), representing the relationships among 120 haploid samples from five species (see *legend*). The tree within the *F*_ST_ peak has two very young clades (*gray boxes*) each of which is enriched for *S. palustris* or *S. nigrorufa* and has a more recent TMRCA than expected given its size (*p*=0.01, **Supplementary Text**). By contrast, the flanking tree is characterized by frequent deep coalescence events and incomplete lineage sorting. (**B**) Times to most recent common ancestry (in millions of generations) for half of the haploid samples (TMRCAH) from *S. nigrorufa*. Values are reduced in the *F*_ST_ peak compared to flanking regions, suggesting species-specific selective sweeps. (**C**) Cross-coalescence times (CC) in millions of generations between *S. nigrorufa* and *S. palustris*, as computed from the inferred ARG. No elevated cross coalescence times are observed in the *F*_ST_ peak, as would be expected if selection against gene flow were predominant. The smoothing method used in panels **B** and **C** was local polynomial regression; the gray bands represent 95% confidence intervals.

To see whether these trends held more generally, we devised statistical tests for (1) enrichment of particular species within clades of these trees, and (2) a corresponding reduction in the within-species TMRCA. The first test identifies instances of species differentiation, and the second identifies likely species-specific sweeps (see **Methods** and **Supplementary Text**). In both cases, we applied stringent significance thresholds based on the results of the tests when applied to genomic scaffolds that do not contain *F*_ST_ peaks. In 23 of the 25 (92%) *F*_ST_ peaks, we observed a significant enrichment for at least one species based on test 1, with 7 peaks (28%) showing significant enrichment for more than one species (**Table 1**). Overall, we detected 35 instances of species-enriched clades in these peaks, with *S. nigrorufa* showing the largest contribution (14 peaks) and *S. hypoxantha* showing the smallest (2 peaks). Of these 35 instances, 18 (64%) in 16 different peaks had significantly reduced within-species TMRCAs compared to the background, based on test 2. Together, these observations suggest a substantial contribution from recent selective sweeps to differentiation between southern capuchino species in these genomic islands.

**Table 1:**
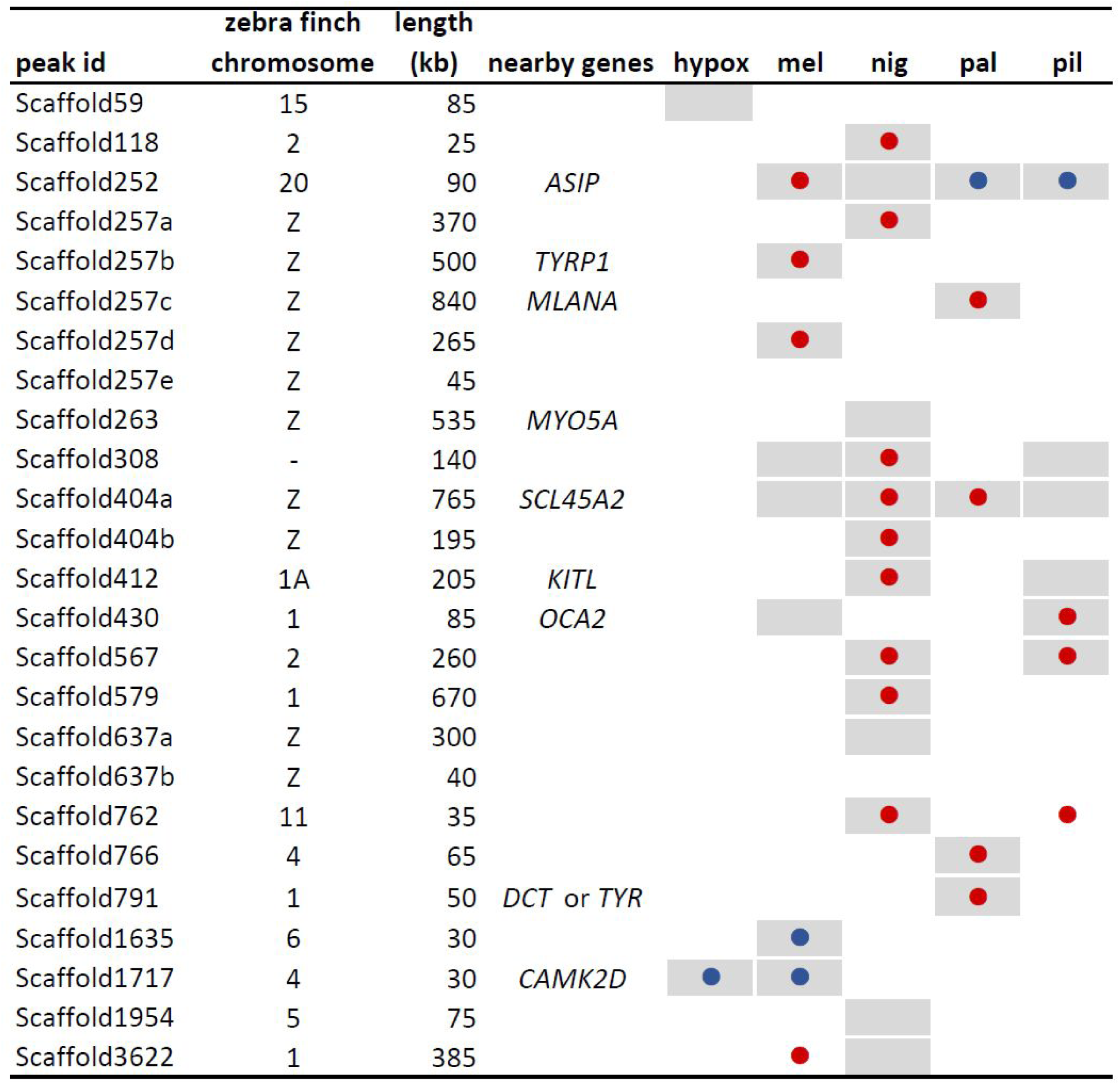
Summary of genealogical signatures of speciation. Results are for the 25 *F*_ST_ peaks previously identified in pairwise analyses of *S. hypoxantha* (hypox), *S. melanogaster* (mel), *S. nigrorufa* (nig), *S. palustris* (pal), and *S. pileata* (pil)^40^. Associated local trees were extracted from the *ARGweaver*-inferred ancestral recombination graph and examined for evidence of species differentiation (test 1: gray cells, *p*<0.0001), young clades indicating recent species-specific selective sweeps (test 2: red circles, *p*<0.0001), and deep separation indicating selection against gene flow (test 3: blue circles, quantile difference > 0.1). See **Methods** and **Supplementary Tables S2, S3 and S5** for details.

We further examined the remaining 17 cases of species enrichment that did not exhibit reduced TMRCAs. These could represent cases in which the *F*_ST_ peaks are driven by early selection against gene flow rather than recent selective sweeps, although they could equally well reflect weaker or older sweeps that failed to meet our criteria for significance. To distinguish between these possibilities, we devised a test for elevated cross-coalescence times between species (test 3), as expected in the case of selection against gene flow at the time of speciation but not in the case of sweeps (see **Methods** and **Supplementary Text**). We tested the statistical power of this method on simulated data (**Supplementary Table S4**), and controlled for false positives using genomic scaffolds that do not contain *F*_ST_ peaks. We found that most *F*_ST_ peaks had either no significant difference in cross-coalescence time with respect to flanking regions, or reduced—rather than elevated—cross-coalescence times (**Supplementary Table S5** and **Supplementary Text**). Nonetheless, three of the peaks did exhibit elevations of cross-coalescence times consistent with selection against gene flow (**Table 1**). The genomic signatures at these peaks are subtle but the genealogical evidence is compelling, particularly in the peaks on scaffolds 252 and 1635 (**Supplementary Figures S4**-**S6**). Notably, the region near the peak on scaffold 252, which is upstream of the gene encoding the Agouti-signaling protein (ASIP), contains multiple signatures of divergence. These include an apparent recent species-specific sweep in *S. melanogaster*, and a modestly recent sweep in *S. nigrorufa*, alongside deep separation for *S. palustris* and *S. pileata*. This deep separation could indicate selection against gene flow in these species, although it could also reflect other phenomena, such as ancestral population structure or balancing selection^46^ (**Supplementary Text**). Altogether, the observed patterns suggest possible selection against gene flow at a few loci of interest but its overall contribution to islands of differentiation appears to be limited compared to that of recent selective sweeps.

### Prediction of Selective Sweeps by Machine Learning

The previous analysis suggests that selective sweeps have played a major role in driving the islands of differentiation observed in these recently emerged southern capuchino species. To characterize these sweeps in more detail, we developed machine-learning methods to predict individual sweeps of various kinds across the available genome sequences. We adopted a prediction strategy based on supervised learning, using a diverse collection of standard population genomic summary statistics as features and a linear Support Vector Machine (SVM) for classification (**Figure 3** and **Methods**). Similar approaches are becoming increasingly widely used in population genomics, and have been shown to be particularly effective in integrating multiple disparate signals to address challenging prediction problems^47,48,44,49^. As in most previous applications, we trained our classifiers from simulated data, which has the advantage of providing essentially unlimited training data with error-free annotation of selection histories, but the potential disadvantage of being sensitive to the choice of simulation parameters (see **Discussion**). To make our method as robust as possible to this choice, we used a forward simulation scheme based on the SLiM^50^ simulator, making use of a previously inferred demographic model^41^ and allowing for ranges of values of key parameters, including selection coefficients, mutation and recombination rates, effective population sizes, and positions of beneficial mutations (see **Methods** and **Supplementary Text**).

**Figure 3:**
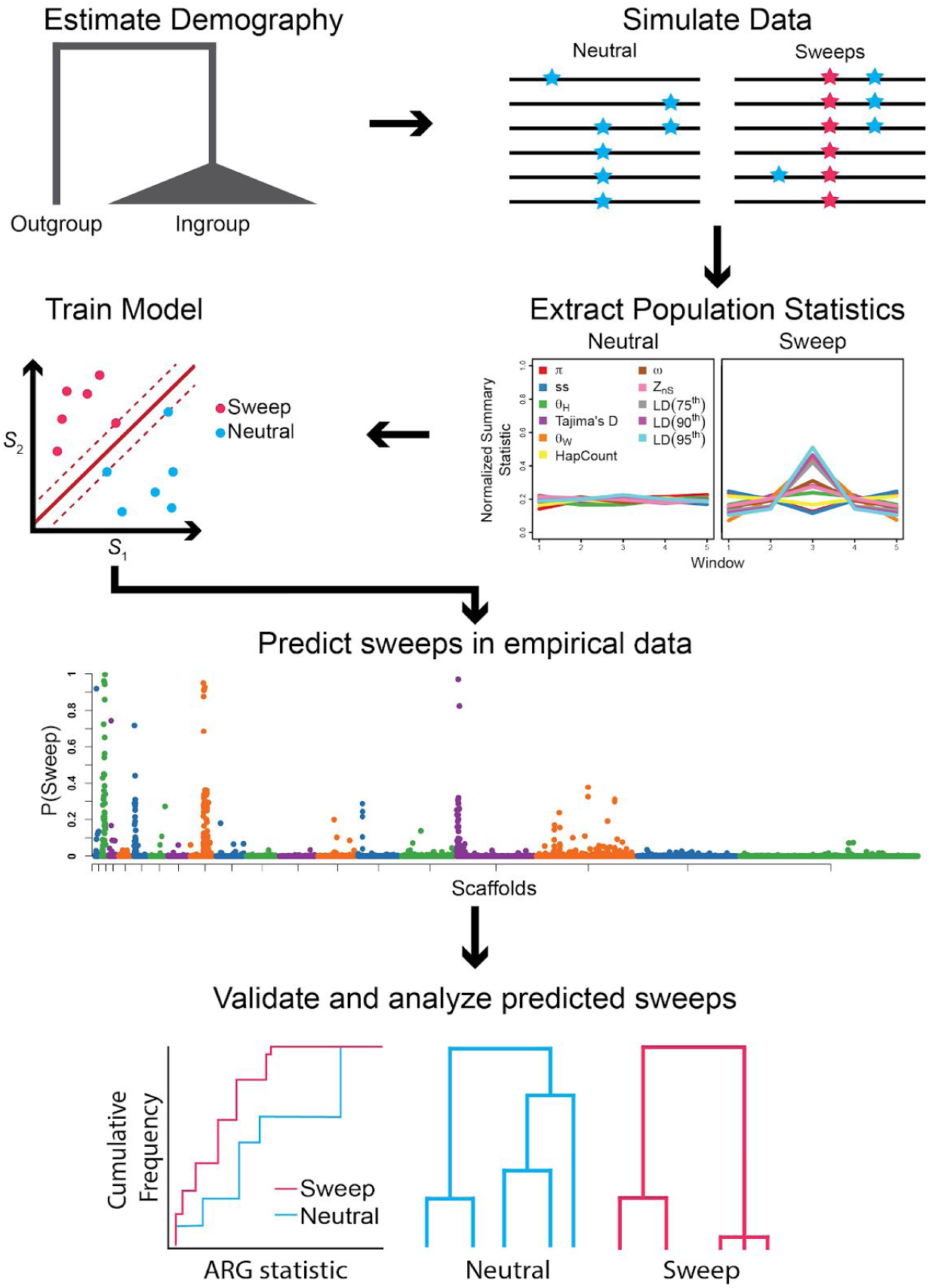
Illustration of machine-learning pipeline for prediction of selective sweeps. Based on an estimated demographic history^41^, SLiM^50^ is used to simulate both neutral genomic regions, and regions containing sweeps. Summary statistics specific to each species are then extracted from each simulated region. The summary statistics used were π (ref. ^78^), the number of segregating sites, Tajima’s *D*^71^, θ_w_ (ref. ^72^), θ_H_ (ref. ^70^), the number of distinct haplotypes^64^, *Z* (ref. ^73^), the maximum value of ω (ref. ^74^), and several statistics that summarize the linkage disequilibrium (LD) distribution (see **Methods**). All statistics were used as features in training a machine-learning model (a linear SVM) to discriminate a sweep from a neutral region. The model detects not only characteristic combinations of features, but also their patterns of variation across genomic windows. The learned model is applied to the empirical data to predict individual sweeps across genomic scaffolds. Finally, the predicted sweeps are analyzed using ARG-based measures (e.g. species-specific TMRCA and RTH), and local trees that map to those predictions are explored for recent clusters of coalescent events. Similar pipelines are used to distinguish between soft and hard sweeps, recent and ancient sweeps, and partial and complete sweeps (see **Methods**).

We were interested in several properties of the selective sweeps that contributed to the diversification of these species, including whether they tended to be “soft” or “hard” sweeps; to be recent or ancestral; or to be “complete” or “partial” sweeps^51,52^. To make these complex and intertwined questions more tractable, we sought first to address them individually, in sequence, and then later considered more complex cases combining multiple factors. Moreover, we focused first on the simple case of a pairwise analysis of present-day species, and simulations involving an ancestral population that diverged into two species, taking advantage of the fact that the five species under study are believed to have diverged from one another at roughly the same time, leading to a high degree of symmetry in the pairwise comparisons. In all cases, we simulated 48 haploid genomes (24 per species) consisting of 9000 regions of length 50 kb, using 8000 regions for training and the remaining 1000 for testing. Each genomic region was analyzed in five nonoverlapping 10 kb windows for signs of selection in the middle 10-kb window (see **Methods**).

### Soft Sweeps have been Common in the Diversification of Southern Capuchinos

We focused first on the distinction between soft and hard sweeps, training a three-way classifier to distinguish species-specific soft and hard sweeps from each other and from neutral regions (see **Methods**). On simulated test sets, the classifier displayed an overall accuracy of ∼93%, with most errors coming from the misclassification of soft sweeps as hard sweeps (∼9% of soft sweeps) or of hard sweeps as soft sweeps (∼6% of hard sweeps). (**Supplementary Table S6** and **Supplementary Figure S7**). In an extensive series of simulations, we found that the classifier was reasonably robust to differences in mutation and recombination rate, misspecified demographic models, misspecified selection coefficients, gene conversion, and other factors (see **Supplementary Text, Supplementary Figure S8-S14, and Supplementary Table S7**). We applied the method to the real data for *S. melanogaster* and *S. nigrorufa*, focusing at first on four scaffolds (252, 412, 404, and 1717) that contain top *F*_ST_ peaks and harbor known pigmentation-related genes (**Supplementary Table S8** and **Figure 4A**). The classifier predicted a total of 154 soft and 33 hard sweeps across these four scaffolds, including high-confidence (probability > 0.95) predictions of 28 soft and 3 hard sweeps. We observed similar patterns in three other pairwise analyses, selected based on their high levels of differentiation^40^ (**Supplementary Figure S15-17** and **Supplementary Table S8**). In general, we identified large numbers of apparent species-specific sweeps, many of which coincided with *F*_ST_ peaks or otherwise occurred nearby genes involved in the regulation of melanogenesis, consistent with the hypothesis that selective sweeps have been a major force behind the differentiation of these species. Furthermore, we consistently observed much larger numbers of predicted soft sweeps than hard sweeps. For example, in a broader analysis of sweeps in *S. melanogaster* (relative to *S. nigrorufa*) in the 19 scaffolds that contain *F*_ST_ peaks for this species, we observed 520 predicted soft sweeps and only 85 predicted hard sweeps, of which 86 and 8 were predicted with probability greater than 0.95, respectively (**Supplementary Table S9**).

**Figure 4:**
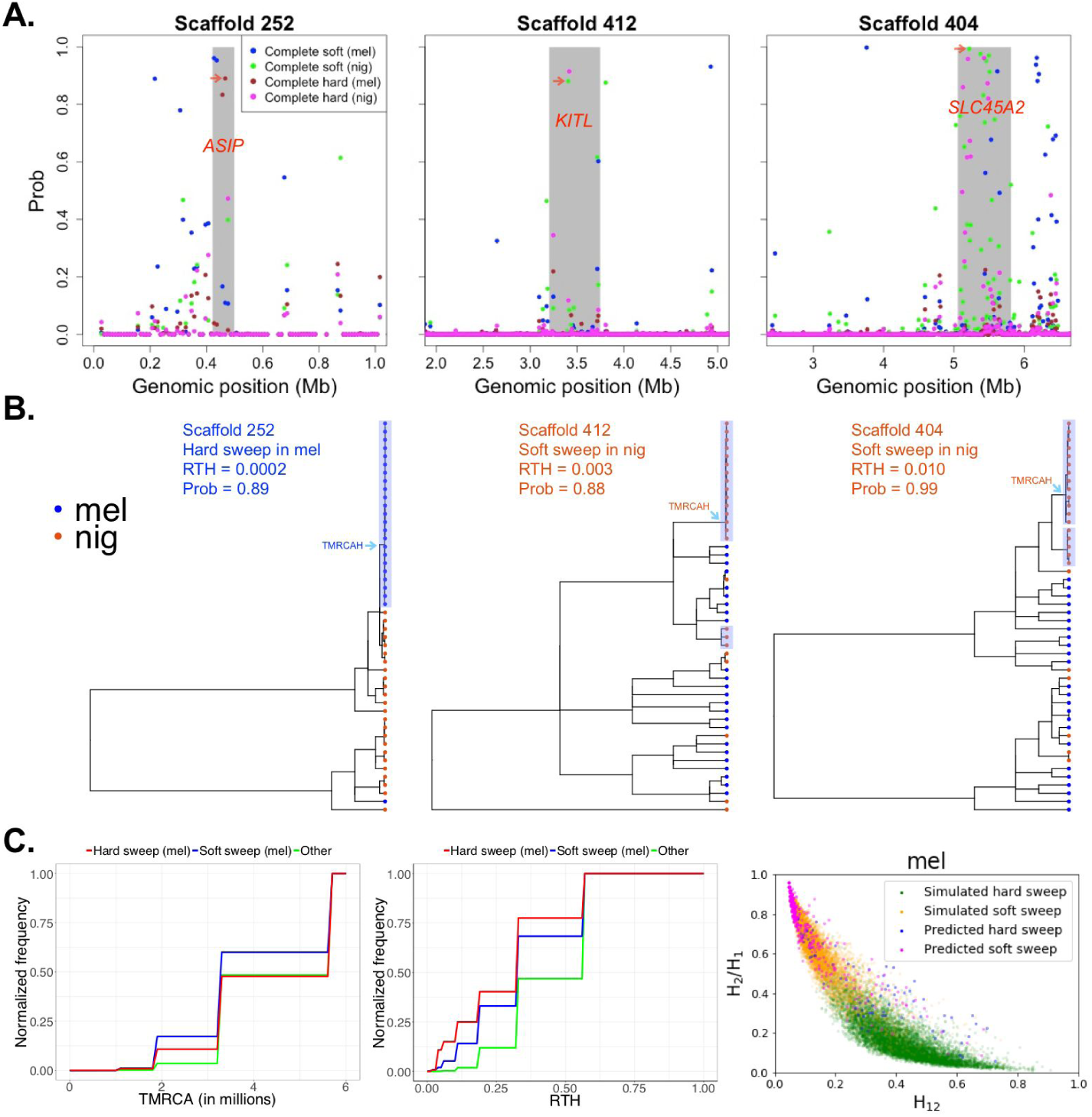
(**A**) Manhattan plots showing the prediction probabilities for soft and hard sweeps in *S. melanogaster* (mel) and *S. nigrorufa* (nig) across three scaffolds harboring top *F*_ST_ peaks and known pigmentation genes (labeled in *red*). (**B**) Local trees inferred by *ARGweaver* in 10 kb windows classified as soft or hard sweeps in *S. melanogaster* (mel) and *S. nigrorufa* (nig). The red arrow in panel A points to the classified window from which each local tree was extracted. Shown also is the RTH of the local tree and the prediction probability for soft or hard sweeps associated with the 10 kb window. Highlighted for each local tree are the two youngest clades containing at least three haploid samples from the target species. We also highlight the youngest clade that contains at least half the haploid samples for the target species (TMRCAH). (**C**) Cumulative distribution functions of *ARGweaver*-based estimates of species-specific TMRCA (*left*) and RTH (*middle*) for *S. melanogaster* in regions classified as hard sweeps (red), soft sweeps (blue), or other classes (green) (see **Supplementary Text**). These statistics are depleted by selective sweeps, owing to clusters of recent coalescent events. (*right*) Relationship between homozygosity-based statistics 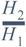 and *H*_12_ (see **Supplementary Text**) for simulated and real *S. melanogaster* data. Each dot in the plot corresponds to a 50 kb region, and its color indicates its predicted (or simulated) class.

To validate these predictions, we made use of several orthogonal analyses. First, we examined many individual predictions in detail, considering the local trees inferred by *ARGweaver* at both predicted hard and soft sweeps (**Figure 4B**). We found, in numerous cases, that the hard and soft sweeps had distinct genealogical features, with the hard sweeps displaying evidence of single derived haplotypes at high frequency, corresponding to unusually large and young clades, and the soft sweeps showing signatures of multiple potentially beneficial haplotypes all at elevated frequencies (**Supplementary Figure S18)**. Second, we carried out a more systematic analysis based on two measures derived from the inferred ARGs: (1) the time to most recent common ancestry (TMRCA) for the species experiencing the sweep, and (2) a related statistic called the relative TMRCA half-life (RTH), which is expected to be reduced in partial or soft sweeps^42^ (**Figure 4C** and **Supplementary Figure S19–20**). We found that both hard and soft sweeps exhibited reductions of both statistics relative to flanking neutral regions. Interestingly, hard sweeps show a lower RTH than soft sweeps, likely because of their recent origin on a single haplotype background.

Finally, to validate our ability to differentiate between soft and hard sweeps, we examined two haplotype homozygosity statistics, *H*_12_ and 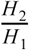, which are sensitive to differences in frequency between the two most prevalent haplotypes^54^ (see **Supplementary Text**). We found, as expected, that the predicted soft sweeps tended to have larger values of 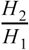 and smaller values of *H*_12_, while the hard sweeps showed the opposite pattern (**Figure 4C** and **Supplementary Figure S21**). Simulated hard and soft sweeps showed qualitatively similar behavior, although, unsurprisingly, they separated more cleanly than the empirical ones. We also verified that few (∼3%) of our predicted soft sweeps fall within the 20kb flanking regions of predicted hard sweeps (see **Supplementary Text**), indicating that our classifier is not frequently misled by the “soft shoulders” of hard sweeps^53^. Overall, our results suggest that the empirical data is indeed dominated by the soft sweeps, and if anything, the classifier has erred on the side of miscalling some soft sweeps as hard sweeps (consistent with tests on simulated data).

### Many of the Soft Sweeps are also Recent and Partial

We expanded our simulation scheme to consider other distinguishing characteristics of selective sweeps. First, focusing on the case of soft sweeps, we asked whether these sweeps tended to occur recently, after a pair of species diverged, or whether they may instead be ancestral, meaning that they began in the common ancestor of the two species. In simulation experiments, we found that we had some statistical power to distinguish these cases, with somewhat better classification accuracy for recent sweeps (∼98%) than ancestral ones (∼80%; see **Supplementary Table S10** and **Supplementary Figure S22**). In particular, the ancestral sweeps and neutral regions tended to be difficult to distinguish, probably owing to loss in older sweeps of the characteristic pattern of extended homozygosity^54^. We applied our classifier to the real data and did obtain predictions of 4–11 times as many recent, species-specific sweeps (per-species) as ancestral sweeps (see **Supplementary Table S11**), although this enrichment likely in part reflects differences in power.

Second, we attempted to ask whether species-specific soft sweeps tended to be “complete,” meaning that the favored allele has been driven completely to fixation, or “partial,” meaning that it has not been fixed. We had limited power to address this question (see **Supplementary Table S12** and **Supplementary Figure S23**) but found some evidence suggesting that a substantial fraction of soft sweeps are partial sweeps (see **Supplementary Table S13**). This observation is consistent with the fact that the reduction in RTH observed in predicted soft sweeps is more pronounced than the reduction in TMRCA (**Figure 4C**).

### An Expanded Analysis of all Five Species Further Supports Abundant Species-Specific Soft Sweeps

Having found that soft sweeps—often recent—have likely been common in these southern capuchino species, we extended our pairwise analyses to consider all five species simultaneously, allowing for a sweep to occur in any one of them. Our goal was to produce a comprehensive set of predictions encompassing all species. Based on simulated data, our multi-way classifier showed good accuracy in this setting (93–95% accuracy; **Supplementary Table S14** and **Supplementary Figure S24**). An analysis of the nineteen scaffolds containing *F*_ST_ peaks identified numerous species-specific soft sweeps (**Supplementary Figure S25** and **Supplementary Table S15**), consistent with our previous pairwise analyses. Furthermore, the inferred local trees associated with these predicted sweeps indicated reduced RTH statistics (**Figure 5** and **Supplementary Figures S26**). Together, these findings further support the prevalence of soft sweeps.

**Figure 5:**
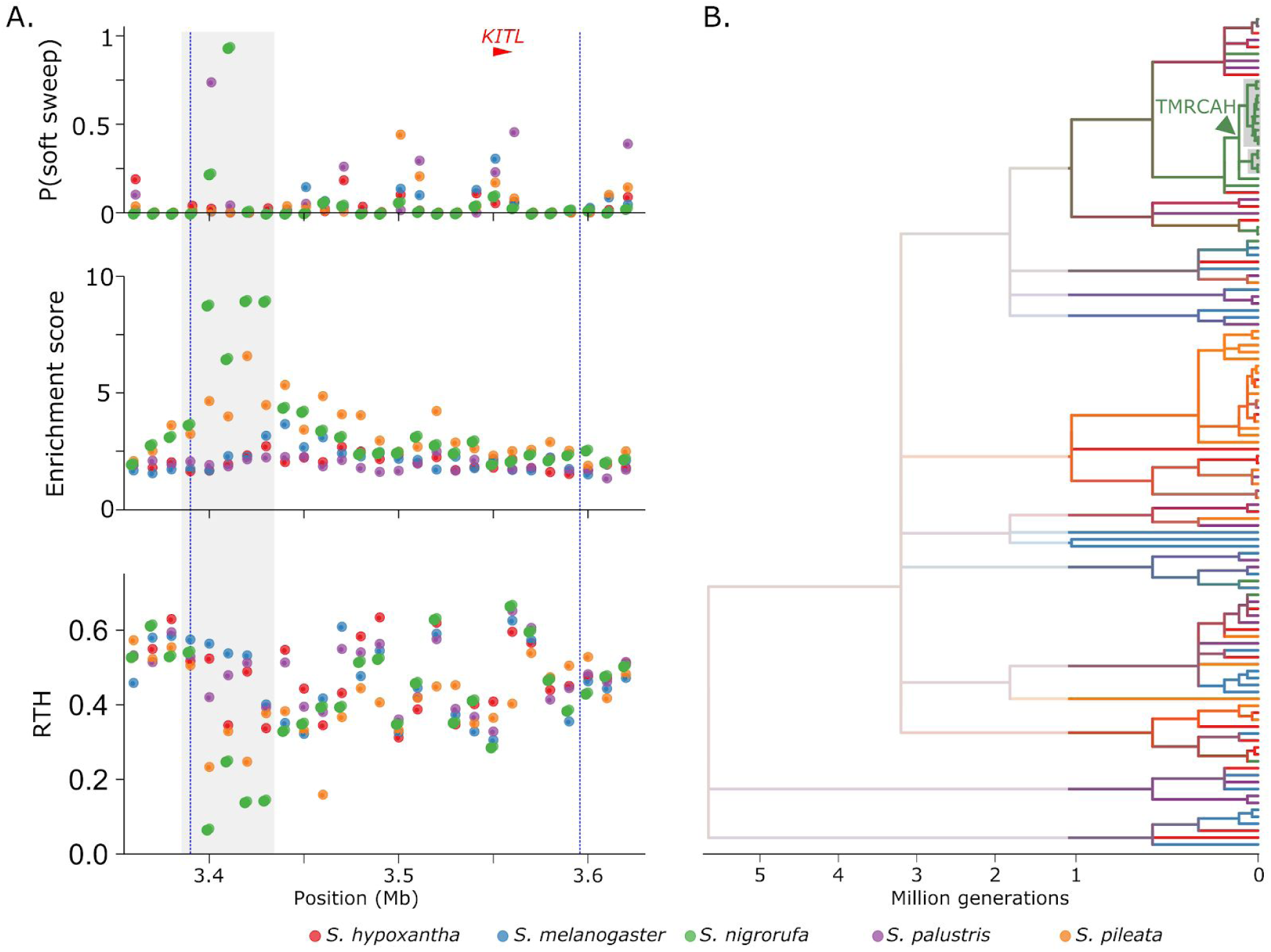
Predictions of the soft sweep classifier expanded to all five species and corresponding genealogical signatures. (**A**) Prediction probabilities for soft sweeps in each of the five species (top; see legend for species color code) in a region containing an *F*_ST_ peak (dashed vertical lines) upstream of the *KITL* gene (red) on scaffold 412. A high-confidence prediction for a soft sweep in S. *nigrorufa* (prediction probability 0.94) is inferred roughly 130 kb upstream of the *KITL* gene. The 50 kb region surrounding this predicted peak (gray bar) exhibits genealogical signatures of a sweep in two ARG-based statistics: (1) high enrichment scores for S. *nigrorufa* (middle) indicating that local trees in this region contain clades enriched for this species, and (2) low RTH (bottom) indicating that the clades enriched for S. *nigrorufa* are young. (**B**) A local tree inferred by *ARGweaver* in the region containing the inferred soft sweep (position 3.43 Mb). Tree tips are colored based on species label (see legend) and color of internal branches represents average over all offspring branches. The figure focuses on the last million generations, with deeper branches faded out and shown in half scale. The tree contains a young clade (∼105,000 generations) with 15 haploid samples, all from S. *nigrorufa* (out of the total 24 samples). This clade contains two major subclades (gray background), which could possibly correspond to two haplotype backgrounds for the inferred soft sweep.

To gain insight into the functional relevance of the predicted sweeps, we examined the Gene Ontology (GO) biological process of nearby genes, focusing on predictions of species-specific soft sweeps with probability >0.95. We identified only two significantly enriched GO processes: “melanin biosynthetic process from tyrosine” (*p* = 0.023 after FDR correction) and “pigmentation” (*p* = 0.022 after FDR correction). Notably, the predicted sweeps were mostly located in non-coding regions near the genes in question, and likely contain cis-regulatory elements that control gene expression. Prominent examples include predictions near the genes *ASIP* (induces melanocytes to synthesize pheomelanin instead of eumelanin), *KITL* (stimulates melanocyte proliferation) (**Figure 5**), and *SLC45A2* (transports substances needed for melanin synthesis).

## Discussion

In this article, we have presented an analysis of the genome sequences of 60 individuals representing five species of southern capuchino seedeaters. These birds serve as an excellent model for recent speciation in the presence of gene flow. Furthermore, because they differ primarily in male plumage and are mostly sympatric, they are a powerful system for studying the genetic effects of sexual selection in speciation^40,41^.

Our analysis focused in large part on previously identified genomic islands of differentiation^40^, defined by local peaks in *F*_ST_. There has been a great deal of interest for more than a decade in the identification of such islands using summary statistics such as *F*_ST_ and *d*_XY_. These summary statistics, however, provide limited information about the evolutionary processes underlying the separation of populations into distinct species. In this case, we further dissected the previously identified regions using a powerful new method for ancestral recombination graph (ARG) inference called *ARGweaver*. This method provides access to fully resolved genealogies, recombination breakpoints, and distributions over coalescence times, as opposed to the coarser averages represented by *F*_ST_ and *d*_XY_. Inspection of individual loci indicates that they often provide clearer indications of the evolutionary events underlying islands of differentiation (**Supplementary Figures S27-29**). Therefore, we designed a series of statistical tests that attempted to exploit this high-resolution information more generally. In addition, we combined these tests with machine-learning methods for the prediction of various types of selective sweeps, in order to gain deeper insights into the processes that led to the observed islands of differentiation.

We were particularly interested in comparing two competing models for the formation of genomic islands of differentiation: one in which selection acts during speciation to reduce gene flow in these genomic regions, and another in which selection acts to reduce within-species diversity through species-specific selective sweeps^32,33^. Notably, the “selective sweeps” model is a distillation of a larger family of models that allows for early sweeps, late sweeps, and adaptive introgression^33,36,45^. Nevertheless, we view these two general paradigms—early barriers to gene flow vs. recent sweeps—as representing a fundamental distinction between plausible models for the formation of islands, with the timing and specific nature of the sweeps being less essential.

At the same time, a binary choice between models is clearly an oversimplification—not only does it mask a diversity of scenarios within each paradigm, but it also obscures the possibility that both paradigms could be simultaneously at play. Importantly, the ARG-based statistics allow us to gain insight into the possible contributions of each model at each island of differentiation. For example, in the island upstream of the *ASIP* gene, we find evidence of a recent near-complete sweep in *S. melanogaster* alongside a deep separation between *S. pileata* and the other four species (**Supplementary Figures S4–S5** and **Supplementary Text**). Thus, both models likely contributed to differentiation in the regulatory sequence of this gene, but at different times and in different species. Notably, the distinction between the two paradigmatic models may not be absolute, since loci that experienced early barriers to gene flow could later undergo selective sweeps, and loci that underwent species-specific sweeps could lead to reduced hybrid fitness resulting in barriers to gene flow. Nevertheless, when we contrast these two extreme scenarios, we find much stronger support overall for *F*_ST_ peaks being associated with recent sweeps than with older barriers to gene flow.

It is worth noting that, while we have focused on selective sweeps as an alternative to barriers to gene flow, previous models have allowed for a reduction in within-species diversity owing to background selection (BGS) as well as selective sweeps^34,36,46^. We focused on selective sweeps because they are far more plausible as an explanation for dramatic reductions in within-species diversity that have occurred recently and in a species-specific manner. In addition, a recent study provides compelling evidence that BGS is unlikely to produce a significant local inflation of *F* ST under realistic population genetic parameters for vertebrates^55^. Moreover, we took care to use indicators of selective sweeps that should be fairly robust to BGS, to avoid mistaking signatures of background selection for sweeps. In particular, our ARG-based analysis made use of measures based on the degree of “clustering” of coalescence events in local trees, which are designed to be sensitive to sweeps to the exclusion of BGS^43^ (see **Methods**). Similarly, haplotype statistics such as *H* and 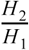 should be elevated by sweeps but not by BGS. Thus, our conclusions about the prominence of sweeps should be relatively unaffected by BGS.

It has recently been shown that *F*_ST_ outlier scans can indirectly enrich for regions of reduced recombination, because the *F*_ST_ statistic has increased variance in such regions^56^. Therefore, the *F*_ST_ peaks we have analyzed could potentially have lower average recombination rates than the background regions to which they were compared. We do not expect such a difference—if it does exist—to have a major impact on our conclusions, owing to our focus on the inferred genealogies in *F*_ST_ peaks and the relative times of coalescence events, which should not be highly sensitive to recombination rates. Moreover, even if the outlier effect were to enrich for distorted genealogies, we do not expect it produce a bias in differentiating between signatures of selection against gene flow (e.g., deep cross-coalescence) and signatures of recent sweeps (e.g., shallow within-species TMRCAs). Nevertheless, it is possible that locally reduced recombination rates in the *F*_ST_ peaks could have some indirect influences on our analysis. Improved recombination maps will be needed for the southern capuchinos or closely related species to enable this issue to be explored in more detail.

Consistent with other recent studies, we found machine-learning methods based on combinations of traditional population genetic summary statistics to be quite powerful in detecting selective sweeps. Notably, we went beyond previous work by characterizing sweeps not only as “soft” or “hard”, but also as ancient or recent, and partial or complete. Overall, we found abundant evidence for sweeps in these capuchino species, and indications that these sweeps are enriched for soft sweeps and recent population-specific sweeps. These conclusions were further supported by semi-independent evidence from ARG inference and haplotype statistics. At the same time, it was clear from our simulation experiments that some features of sweeps were much more difficult than others to discern from sequence data. For example, we had limited power to identify ancient sweeps and partial sweeps. We also attempted to distinguish between sweeps that were specific to a single species and sweeps that occurred in parallel at the same locus in multiple species, but found that our results were somewhat ambiguous, with true parallel sweeps being mixed together with cases of apparent shared sweeps or adaptive introgression (**Supplementary Table S16 and Supplementary Figures S30–S32**). In addition, our current machine-learning methods are still coarse-grained, in that they identify broad windows containing sweeps rather than specific causal variants, and they do not make direct use of informative ARG-derived features in classification. An important area for future work will be to develop improved high-resolution ARG-aware machine-learning predictors of sweeps^49^, taking advantage of the latest advances in genealogy and ARG inference^43,57,58^.

As in all studies that predict sweeps using machine-learning classifiers trained from simulated data, our methods are sensitive to biases stemming from our choices of parameters for simulation^44,49^. We attempted to mitigate this issue by simulating from a broad family of models, and by validating our predictions with independent methods where possible. In addition, we systematically evaluated the influence on our predictions of features such as misspecified demographic models, selection coefficients, mutation and recombination rates; the addition of gene conversion; and the “soft shoulders” phenomenon (**Supplementary Text**); and we found that our methods were fairly robust to these confounding factors. Nevertheless, training bias and limitations in both power and resolution have undoubtedly influenced our predictions to a degree.

Regardless of these caveats, the prediction of abundant soft sweeps—largely recent and population-specific—makes sense from first principles and previous findings. These capuchino species appear to have emerged from a large, highly polymorphic ancestral population, which would have provided high levels of standing variation to serve as the substrate for soft sweeps. Furthermore, these speciation events appear to have been quite recent, which might favor soft sweeps over hard sweeps, because they can occur more rapidly^59^. It has further been argued that standing genetic variants may be more likely than novel ones to drive speciation^60^, because novel mutations tend to be deleterious if not neutral^61^, whereas older variants have already been filtered and shaped by selection in their ecological context^62^. Notably, other recent empirical studies have similarly found a predominance of soft selective sweeps in both *Drosophila melanogaster*^63,64^ and humans^65^. The recent speciation event would also allow limited time for haplotypes under selection to reach fixation, which would favor partial sweeps over complete sweeps. Finally, the likely role of sexual selection in driving many of these sweeps is consistent with selected alleles for features such as plumage having been present in the ancestral population, likely at low frequency, and having swept to high frequency relatively recently, in a species-specific manner. Sexual selection would also be consistent with our observations suggesting that many loci have experienced sweeps in parallel in different species. We note that there is now strong evidence—including genetic, behavioral, coloration, song and captive breeding data^66–69^—that these are truly distinct species, not artificial taxonomic constructs based on superficial morphological features.

Overall, the genetic changes underlying any speciation event are complex and unlikely to be fully explained by any simple model. They are undoubtedly highly context-dependent, depending on diverse features such as population size and structure, rates and patterns of gene flow, degrees of sympatry, differences in local environments, and the strength of sexual selection. Nevertheless, our analysis of these capuchino species demonstrates that prominent genomic islands of differentiation can be explained largely through recent, species-specific selective sweeps. Furthermore, we have shown that the analysis of these islands can benefit substantially by going beyond summary statistics such as *F*_ST_ and *d*_XY_ and making full use of the ancestral recombination graph (ARG), as well as machine-learning methods for prediction of selective sweeps. Together, our observations help to fill in the picture of how selection, recombination, and drift act together to shape the genomes of distinct species, with broad implications across the tree of life.

## Materials and Methods

### Obtaining local trees from genome sequence data

All of our analyses are based on the genome sequence data published by Campagna et al.^40^. We applied standard filtering and executed *ARGweaver* on the entire data set, producing an inferred local tree for every position along the sequenced genome scaffolds (**Supplementary Text**).

### Summarizing genealogical signatures from the inferred ARG

Genealogical information was summarized using several measures extracted from each local tree (see below). Average values for each measure were computed in non-overlapping 20 kb windows tiling all scaffolds (**Supplementary Text**). A significance threshold was determined for each measure separately by considering the distribution of values observed in the 39,699 windows that cover the 576 scaffolds that do not contain *F*_ST_ peaks. We applied thresholds associated with a strict empirical *p*-value of 0.0001. Moreover, to account for the use of extreme values (maximum or minimum) in long genomic blocks, we examined the distribution of values for each measure in a collection of 1,376 non-overlapping 500 kb blocks from the same 576 scaffolds. The block length was selected to be conservative relative to the length distribution of *F*_ST_ peaks (mean length 243 kb and only four peaks longer than 500 kb; **Supplementary Table S1**).

### A test for species differentiation (test 1 in Table 1)

Scores corresponding to species differentiation in local trees were determined using a hypergeometric test, as follows. For a subtree with *n* leaves out of which *k* are mapped to a certain species, we computed the probability of this observation under a hypergeometric distribution, and defined the associated enrichment score as -log_10_(P_*K=24,N=120*_(*X*≥*k*|*n*)). The enrichment score associated with a given species in a given site is defined as the maximum score associated with that species in a subtree of the local tree inferred in that position. Thus, a high enrichment score is obtained when the local tree contains a subtree enriched for individuals from that species. Enrichment scores were averaged in non-overlapping 20 kb windows, and an empirical *p*-value of 0.0001 was determined separately for each species by considering the top four values observed in the set of 39,699 windows used for control (see above). An *F*_ST_ peak was considered to exhibit significant differentiation for a given species (gray cells in **Table 1**), if the peak contained a 20 kb window with an enrichment score for that species that exceeded its empirical significance threshold (See **Supplementary Table S2** for the species-specific thresholds and a complete set of results). Notably, the enrichment scores based on the hypergeometric distribution reflect a simplifying assumption of exchangeability of all lineages across species; however, the empirical control ensures that violations of this assumption do not produce a bias in the identified peaks.

### A test for reduction in within-species time to most recent common ancestry (TMRCA) (test 2 in Table 1)

We based this test on the time to the most recent common ancestor of half the haploid samples (*n*=12) from a given species (TMRCAH). Requiring only half the samples allows us to consider partial sweeps and provides robustness to the inherent uncertainty in the inferred local trees. To account for variation in age estimates stemming from mutation rate variation and/or background selection, we used a relative version of TMRCAH, which, following ref. ^42^, we denote as the Relative TMRCA Halflife (RTH). Originally, RTH was defined as the TMRCAH of a given species divided by the full TMRCA of that species, but for test 2, we used a slightly modified measure (RTH’) obtained by dividing the TMRCAH by the age of the youngest subtree that contained at least half of all samples, not only those samples from the species in question (*n*=60). This measure provides improved robustness to errors in tree inference and potentially captures a wider variety of selective sweeps (**Supplementary Text**). RTH’ is related to the species-specific sequence diversity (π_Within_) that has been used in other studies of islands of differentiation, but we expect RTH’ to be more sensitive to partial sweeps and less affected by mutation rate variation and background selection (**Supplementary text** and **Supplementary Figure S28**). Values of RTH’ were averaged in non-overlapping 20 kb windows, and an empirical *p*-value of 0.0001 was determined separately for each species by considering the lowest four values observed in the set of 39,699 windows used for control (see above). An *F*_ST_ peak was considered to exhibit a significant reduction in TMRCA in a given species (red circles in **Table 1**), if the peak contained a 20 kb window with an RTH’ for that species that is below its empirical significance threshold (See **Supplementary Table S3** for the species-specific thresholds and a complete set of results).

### A test for elevation in cross-coalescence time (test 3 in Table 1)

For a given local tree and pair of species, we considered the ten most recent cross-coalescent events between the two species and normalized these ages, as in test 2, by the age of the youngest subtree that contains at least half of the total number of haploid samples. These measures are related to *d*_XY_, but are expected to be more sensitive to recent changes in gene flow and less affected by mutation rate variation and background selection (**Supplementary Text** and **Supplementary Figure S29**). Rather than computing average values across 20 kb windows, we computed for each species-pair the distribution of normalized recent cross coalescence times in every *F*_ST_ peak and compared this distribution to the one observed in flanking regions. Quantile differences were used to measure the difference between the two distributions, and a significance threshold associated with an empirical *p*-value of 0.01 was set based on examination of 1,367 genomic regions from scaffolds that do not contain an *F*_ST_ peak (**Supplementary text**).

### A test for possible selection against gene flow

Because elevated cross coalescence times could be caused by a variety of phenomena (**Supplementary Text**), we applied additional conditions to detect deep clades enriched for lineages ancestral to a particular species, as expected from selection against gene flow. In particular, we infer possible selection against gene flow for a given species in a given *F*_ST_ peak (blue circles in **Table 1**) if these conditions hold:

1. The species has a significantly high enrichment score in the peak (passes test 1).
2. The species does not have a significantly low RTH’ in the peak (fails test 2).
3. The species has significantly elevated cross-coalescence times with some other species relative to the flanking regions (passes test 3).

### Summary statistics used as features for prediction of different models for selective sweeps

sSeveral sequence-based species-level summary statistics were used as features for a machine learning approach to distinguish among different models of selection (including neutral drift). All statistics were collected in five consecutive 10 kb windows with the objective of identifying possible sweeps induced by a positively selected mutation in the third (middle) window. Some of these summary statistics corresponded to standard measures of diversity, such as the number of segregating sites, π (ref. ^70^), Tajima’s D (ref. ^71^), θ_W_ (ref. ^72^), θ_H_ (ref. ^70^), the number of distinct haplotypes (ref. ^64^), Z_nS_ (ref. ^73^), and maximum value of ω (ref. ^74^). For each of these statistics, we computed an average value for each of the five 10 kb windows for each analyzed species separately and jointly for all species together. We also extracted statistics based on the distribution of the coefficient of linkage disequilibrium (*D*_*AB*_) of each of the five windows with the middle window—where the site under selection is assumed to be located. *D*_*AB*_ was computed for every variant site in the middle window with each other variant site across all five windows. Then, for each of the five windows, we extracted the 75th, 90th, and 95th percentiles of the distribution of *D*_*AB*_ values relative to the middle window. Each of these LD-based statistics was recorded for each analyzed species separately. Finally, each summary statistic was normalized by dividing the value recorded for a given window by the sum of values across all five windows (see **Figure 3**).

### Simulated datasets used for training and testing the selective sweep classifier

Training and testing data sets were generated using SLiM^50^ by simulating 9,000 regions of length 50 kb for each model we considered: (e.g., “neutral”, “soft sweep”, or “hard sweep”). Of these regions, 8,000 were used for training and 1,000 were used for testing (see below). The number of simulated species and sampled sequences was set to match the analyzed data set. Thus, in the species-pair analyses, a total of 48 haploid sequences were sampled at the end of the simulation (24 per species), and in the expanded analysis the entire set of five species was simulated and 120 haploid sequences were sampled. Simulations used a demographic model based on one inferred previously from ddRAD data^41^. To demonstrate how well the simulations fit the empirical data at hand, we applied PCA to the summary statistics extracted from both the empirical data and the simulations based on the demographic model inferred from RAD-seq data (see **Supplementary Figure S33**). In the expanded analysis we used the complete model, and in the species-pair analyses we used the appropriate two-species derivatives of this model (**Supplementary Text**). In non-neutral simulations, selection was applied to a single focal site located in the middle 10 kb window. We explored a range of values for the main parameters corresponding to demography and selection: (1) mutation rate, (2) recombination rate, (3) effective population sizes, (4) selection coefficient, (5) onset time of selection, and (6) position of beneficial mutation. Furthermore, for soft sweeps (partial or complete), we also varied the initial derived allele frequency at which a mutation switches from evolving under drift to becoming beneficial, and for partial sweeps we varied the target derived allele frequency at which a mutation switches back to being neutral (**Supplementary Text**). For each simulated region, we recorded the set of features used by the classifier (see above). If one of these statistics fell outside of the range of values observed in the genomic data, the simulated 50 kb segment was excluded from the set and replaced by another segment (to maintain a total of 9,000 simulated segments per class). The comparison to genomic data was done based on the 19 scaffolds that contain *F*_ST_ peaks and the specific species being analyzed by the classifier. Thus, slightly different filtering was applied for each of the four species-pair analyses in each classification task (see below).

### Training a linear SVM to classify different modes of selection

A linear support vector machine (SVM) was applied to the simulated training data sets to learn a classification model for each task separately. The linear SVM classifies samples into two categories (e.g., “neutral” vs. “non-neutral”) by finding the hyperplane that maximizes the separation of the data from the two different classes^75,76^ (**Figure 3**). Since most of our prediction tasks had more than two categories (e.g. “neutral”, “soft sweep”, and “hard sweep”), we used a one-vs-rest approach. Thus, a separate classifier was trained for each class, with the 8,000 regions simulated for that class as positive training examples, and regions simulated under other models as negative training examples. We used the C-Support Vector Classification tool (*sklearn*.*svm*.*SVC*) from the Python *sklearn* package to train and test the classifier. Default operating parameters were used in all cases. Classification into more than two categories was obtained by combining the results from multiple binary classifiers using the *predict_proba* function in *sklearn*. This function applies a logistic function to the individual score produced by each classifier, thus transforming it into a value reflecting the probability that the binary classifier assigns to its “positive” class. It then divides the probability obtained (separately) for each class by the sum of all probabilities, to obtain a normalized prediction probability for each class. The resulting multi-way classifier was then tested using 1,000 regions simulated for each class. The results of these tests were summarized in a confusion matrix describing the frequency of different types of prediction errors when the predicted class is set as the one with the highest probability. We also computed a receiver operating characteristic (ROC) curve for every pair of classes, to provide a more complete summary of the behavior of different types of errors. We report the calibration curves in **Supplementary Figure S34** to assess the calibration of the probabilities generated by the classifier.

### Robustness study

Our machine-learning approach requires making subjective decisions about which types of examples to simulate and will naturally be biased towards the assumed scenarios. Therefore, it is important to test the model with simulations representing alternative scenarios. We have carried out a fairly extensive analysis of the robustness of our approach, considering not only alternative demographic parameters (such as ancestral Ne, derived species Ne, divergence time), but also alternative parameters for recombination rate, mutation rate, selection coefficients, and gene conversion. In all cases, we took care to test our prediction methods under parameters well outside the range used in training. This analysis is summarized in the **Supplementary Text**. We found that our method was fairly robust to alternative parameter values, although, as expected, performance did degrade somewhat under severely misspecified models (see **Supplementary Figures S8-14**).

### Species-pair analyses and expanded analysis of all five species

We started by distinguishing among different types of selective sweeps using a series of classification tasks, each focused on two types of sweeps and two species. The pairs of sweep categories considered were (1) soft vs. hard sweeps, (2) recent vs. ancestral sweeps, (3) partial vs. complete sweeps, and (4) species-specific vs. parallel sweeps (see **Supplementary Text** for more details). In each case, we aimed to distinguish the two types of sweeps from each other and from neutrally evolving regions. Each of these four classification tasks was applied to the four species-pairs that overall exhibited the highest *F*_ST_ values genome-wide: (1) *S. melanogaster* vs. *S. nigrorufa*, (2) *S. nigrorufa* vs. *S. pileata*, (3) *S. pileata* vs. *S. palustris*, and (4) *S. hypoxantha* vs. *S. melanogaster*. Note that for each species-pair, simulated segments were filtered using slightly different empirical ranges (see above). Based on the outcome of the four classification tasks in all four species pairs, we designed an expanded analysis of all five capuchino species. In particular, in this analysis we allowed any one of the five species to undergo a species-specific complete soft sweep.

### Application of machine learning to genomic data

We analyzed the 19 scaffolds that contain *F*_ST_ peaks using each of our trained classifiers. Each scaffold was scanned by a sliding 50 kb window along the scaffold with a step size of 10 kb. Summary statistics were extracted, as described above, for each 50 kb window using the genome samples appropriate for the specific classification task at hand—24 genomes in each species-pair analyses and all 60 genomes in the expanded analysis. These summary statistics were then provided as input to the trained binary classifiers and their classification outputs were combined to provide a normalized probability for each class (see above). The middle 10 kb of the 50 kb window was then assigned the class with the highest score, and Manhattan plots were used to show the distribution of class assignments and their normalized probabilities across each scaffold (e.g., Figure 4A).

## Acknowledgments

The authors would like to acknowledge Noah Dukler for help with figure preparation. This research was supported by US National Science Foundation grants (NSFDEB) 1555769, 1555754, US-Israel Binational Science Foundation grant BSF2015523, and US National Institutes of Health grant R35-GM127070. The content is solely the responsibility of the authors and does not necessarily represent the official views of the US National Institutes of Health or the US National Science Foundation. The bird illustrations in Figure 1 are original drawings done by Jillian Ditner.

