## Supplemental File for "Genomic islands of differentiation in a rapid avian radiation have been driven by recent selective sweeps"

### Table of contents

**Supplementary Text** **3**
1. The data set 3
2. Supplementary methods for the ARG-based analysis 3
3. Supplementary results and discussion for the ARG-based analysis 11
4. Supplementary methods of models for selective sweeps using machine learning 16
5. Robustness of the selective sweep classifier 20

**Supplementary Tables** **23**

**Supplementary Figures** **35**

**References** **68**

### Supplementary Text

#### 1. The data set.

**Whole genome data set.** Our analysis made use of the reference genome previously assembled for S. *hypoxantha*[1]. The assembly covers 1.17 Gb of sequence partitioned into 5,120 scaffolds. The population-level sequencing data from that paper consisted of 72 individuals from nine capuchino seedeater species. We applied stringent filtering by masking sites with the following quality parameters: *QD*<2, *FS*>40.0, *MQ*<20.0, or *HaplotypeScore*>12.0. Variants were subsequently filtered if they satisfied one of the following conditions: (1) had more than two alleles, (2) had a minor allele frequency smaller than 0.1, (3) had mean sequencing depth smaller than 2, (4) had mean sequencing depth greater than 50, (5) had genotypes for less that 80% of the sequenced individuals (57 out of 72). This filtering pipeline produced 11,530,110 SNPs genotyped for 72 individuals of the nine southern capuchino species. Since four of the nine species—*S. ruficollis*, *S. hypochroma*, *S. cinnamomea*, and the outgroup *S. bouvreuil*—were represented by only three individuals, we focused our analysis on the five species for which there are 12 individuals: *S. hypoxantha, S. melanogaster*, *S. nigrorufa, S. palustris*, and *S. pileata*. The same filtered sequence data was used in both parts of the analysis: the ARG-based analysis and the SVM-based predictions of selective sweeps.

**$F_{ST}$ peaks.** We considered the 25 $F_{ST}$ peaks defined in ref. [1] (**Supplementary Table S1**). These regions were determined as stretches of two or more non-overlapping 25 kb windows with average $F_{ST}$ above 0.2 in at least one of the ten pairwise comparisons, and containing at least one variant site with $F_{ST}$ above 0.85. Windows with fewer than ten variant sites were not considered (see ref. [1] for more details).

#### 2. Supplementary methods for the ARG-based analysis.

**Using *ARGweaver* to infer local trees.** One of the main challenges in extracting reliable signatures of local genealogies in population-level genomic data sets is that recombination frequently modifies the local tree along the genome. As a result, local trees cannot simply be inferred just by using sequence data from the sites that evolved under these trees. Reliable inference of local genealogies can be done by making use of the ancestral recombination graph

(ARG), which captures the correlation between nearby trees, through an explicit model of coalescence and recombination. Several methods for ARG or tree inference from individual genome sequence data have been developed in recent years [2–5], but *ARGweaver* [4] is the only available method that can reconstruct complete ARGs (containing recombination events and local trees), that can accommodate unphased data, and that scales to data sets of the size considered here. The method accepts a collection of long unphased aligned sequences, and it generates samples of ARGs for these sequences via a Markov chain Monte Carlo (MCMC) algorithm. While the Bayesian sampling algorithm is a strength of *ARGweaver*, for simplicity we base all of our analyses here on a single high-probability sample of the ARG. As discussed below, we extract various features of local genealogies, haplotypes, and recombination events from this reconstructed ARG.

**ARG blocks.** To facilitate effective execution of *ARGweaver*, we filtered and partitioned the sequence scaffolds according to the following process. First, due to the low confidence in local genealogy inference near the edges of the aligned sequences, only scaffolds of length at least 120 kb were considered. This filter eliminated roughly 110 Mb of sequence, resulting in 1.06 Gb of sequence analyzed by *ARGweaver*. Among the remaining 595 scaffolds, those shorter than 2 Mb were analyzed as is, and longer scaffolds were split for computational efficiency into blocks of similar length (< 2 Mb) with an overlap of 100 kb. This resulted in 1,003 genomic blocks (ARG blocks) which were analyzed via separate parallel runs of *ARGweaver*.

**Executing *ARGweaver*.** We applied *ARGweaver* to probabilistic genotypes specified in the previously generated sequence VCF file[1]. Genotypes based on sequence coverage less than 5 or greater than 30 were masked using an individual mask file, as well as genotypes with certainty below 80% (`--mask-uncertain 0.8`). We used a site compression of 5 when possible (`-c 5`), and reduced compression in ARG blocks where the frequency of variant sites was too high. To make the analysis more robust to mutation rate variation, we supplied *ARGweaver* with local mutation rates estimated using two individuals sequenced by Campagna *et al.* (2017)[1] from two species not analyzed in this study: L20.3 from *S. hypochroma* and L15.3 from *S. bouvreuil* (individuals from these two species were not used in our subsequent analyses). The absolute divergence between these two individuals was computed in 100 kb windows with 10 kb overlaps using probabilistic genotypes given in the sequence VCF file. Windows with fewer than 20,000 unfiltered bases were assigned the genome-wide average rate. The mutation rate at each genomic position was set by scaling the assumed average mutation rate of $10^{-9}$/bp/gen by the absolute divergence in the 100 kb window centered at that position divided by the average divergence across all windows. In addition, we used the following options for setting the effective population size, recombination rate, time discretization, random genotype phasing, and MCMC sampling: `--popsize 1000000 -r 1e-9 --ntimes 20 --maxtime 1e7 --delta 0.005 --randomize-phase 1.0 --unphased --sample-phase 10 --resample-window-iters 1 --resample-window 10000 -n 1000`. *ARGweaver* was executed on each ARG block for 1000 MCMC iterations, and the ARG sampled in the last iteration was used to extract local trees for all subsequent analyses. In one of the ARG blocks that does not overlap an $F_{ST}$ peak (the fourth ARG block for scaffold 329), *ARGweaver* was halted after 700 iterations due to time constraints. While *ARGweaver* was executed on the entire data set, consisting of 72 diploid individuals, all of our downstream analyses made use of the 60 individuals from the five focal species of this study. Thus, every local tree was collapsed to a tree containing 120 leaves, each associated with a haploid sample from the 60 individuals (two haploid samples per individual).

**Significance test used in Figure 2 and Supplementary Figures S3** and **S4 for young and large clades.** In each $F_{ST}$ peak, we considered the site with the largest $F_{ST}$ value and examined the local tree inferred by *ARGweaver* at that site. For illustrative purposes in the figures, we simply manually selected a subtree (clade) in the local tree that appeared to be a plausible candidate for a species-specific (partial) sweep (i.e., by being fairly large, young in age, and enriched for haploid samples from one of the species exhibiting a high local $F_{ST}$ value). To determine the significance of the young clade age relative to its size, the age and size were compared to a background distribution based on clades of identical size 10,000 local genealogies sampled uniformly at random along the same scaffold outside of the $F_{ST}$ peak. If a sampled tree did not contain a clade of the target size, it was discarded, and if it contained multiple such clades, the age of the youngest one was recorded. A one-side empirical *p*-value was computed for the age of the target clade as the probability of observing an age at least as young as the observed one in the background distribution.

**Normalizing coalescence ages in local trees.** Several of our ARG-based measures, such as TMRCAH and cross-coalescence times, are based on times of coalescence events in local trees. The coalescence times inferred by *ARGweaver* could be influenced by local mutation rate and reduced local effective population size due to background selection. We thus normalized these ages to account for these effects and reduce the variance in ages observed in the genomic background. In most cases, we divided the time of interest (TMRCAH or cross coalescence time) by the age of the youngest subtree that contained at least half the total number of samples (n=60). This normalization appears to reduce much of the variation in coalescence times observed along the genome (**Supplementary Figures S28-S29**). We note that RTH', obtained by this normalization, is reduced by a wide variety of selective sweeps, such as partial sweeps shared by multiple species, or complete species-specific sweeps. A slightly different normalization approach (RTH), obtained by dividing the TMRCAH of a given species by the TMRCA of that species, was used in the validation of the soft sweeps inferred by the SVM (**Supplementary Text 4**). This normalization is designed to be more sensitive to partial or soft species-specific sweeps.

**Summarizing ARG-based measures in 20 kb windows.** Each sequence scaffold was tiled by non-overlapping 20 kb windows after masking out the 50 kb near each edge (due to inference uncertainty in the edges of every ARG block). In every 20 kb window, we computed an average value for each measure by using 40 representative local trees from that window, equally spaced 500 bp apart starting 250 bp from the edge of the window. To assess significance thresholds, we considered the distribution of each measure in 39,699 windows that belong to one of the 576 sequence scaffolds that do not contain $F_{ST}$ peaks. We set a significance threshold with an empirical p-value of 0.0001 by requiring that only three control windows exceed the threshold (tests 1 and 2 in Table 1).

**A test for elevation in cross-coalescence time (test 3 in Table 1).** To detect possible selection against gene flow, we looked for an elevation in times of inter-species cross coalescence (CC) within $F_{ST}$ peaks relative to times of CC observed in nearby genomic regions. In the three sections below, we describe the measure we used to capture recent CC times, the measure we used for difference in CC times between a peak and its flanking region, and a method to control for false positives. **Estimating recent times of cross coalescence.** For a simple and robust measure of cross coalescence (CC) time (used in **Figure 2** and **Supplementary Figure S4**), we used the average of the times of all CC events in a given local tree. Note that an individual coalescence event may be counted multiple times toward this distribution if it is the most recent common ancestor of multiple pairs of samples from the two species. Thus, this measure could be viewed as an ARG-based version of $d_{XY}$. In the detailed analysis of $F_{ST}$ peaks (test 3 in **Table 1** and **Supplementary Table S5**), we used a slightly different version that considered only the ten youngest CC events divided by the age of the youngest subtree that contains at least half of the total number of haploid samples. This version is designed to be more sensitive to recent changes in the amount of gene flow (due to the focus on recent events), and less sensitive to the influence of mutation rate variation and background selection (due to the normalization; see **Supplementary Figure S29**).

**Comparing cross coalescence times in $F_{ST}$ peaks to flanking regions.** For every $F_{ST}$ peak and every species-pair, we computed the distribution of normalized recent CC times in the peak and compared this distribution to the one observed in flanking regions. For peaks longer than 200 kb, we focused on the 200 kb region that contains the highest species enrichment scores. To account for possible fluctuations in coalescence times between sequence scaffolds, we used a local control set consisting of the 200,000 sites flanking the $F_{ST}$ peak. When possible, we selected 100 kb from each side of the peak, unless the peak was close to the edge of a sequence scaffold. The peak on scaffold 308 was discarded from this analysis, because there are less than 200 kb flanking the peak on the sequence scaffold. The difference between the distribution of normalized CC times in an $F_{ST}$ peak and that recorded in the flanking region was quantified using quantile differences. The quantile difference between two distributions was computed by taking the median value of one distribution, examining the quantile of this median value in the second distribution, and taking the difference in absolute value between this quantile and the value ½. This calculation is done in both directions ($F_{ST}$ peak versus flanking region and vice versa), and the difference associated with every peak is taken to be the maximum of the two. See **Supplementary Table S5** for the quantile differences computed for all $F_{ST}$ peaks and all species pairs.

**Controlling false positive rates in tests for elevated cross coalescence times.** We empirically determined thresholds for statistically significant quantile differences in CC times by running a similar analysis on a collection of control regions. For this control, we used the 1,376 genomic blocks of length 500 kb taken from the 576 scaffolds that do not contain $F_{ST}$ peaks. In each block, we used the middle 200 kb to model a region of interest (paralleling an $F_{ST}$ peak), and the two adjacent 100 kb blocks as flanking regions. For each of the ten species pairs and 1,376 genomic blocks, we computed the quantile difference between the distributions of recent CC times in the region of interest relative to the flanking regions, similar to our analysis of $F_{ST}$ peaks. The top and bottom percentiles of this distribution of quantile differences was obtained at thresholds of ±0.21. Importantly, we observed only minor differences between the distributions of quantile differences obtained for the different species pairs. Thus, thresholds of ±0.21 provide robust empirical *p*-values of 0.01 for significant elevation or reduction in CC times.

**Assessing the statistical power of our method for detecting selection against gene flow.** According to the demographic model inferred by Campagna *et al.* (2015)[6], the five capuchino species diverged less than 50,000 generations ago and had ancestral effective population sizes as large as 15,000,000. Under this model, we expect large variation in coalescence times even in the absence of selection, which potentially poses a major challenge for our method, since it depends on identifying local regions with unusually deep cross coalescence. We therefore sought to use simulation experiments to determine how early selection against gene flow had to have started so that our method could detect it. In the five sections below, we describe the simulation framework we developed to address this question, we provide details about our experiments, summarize the results, and conclude with a brief discussion about the implication of these results to our analysis of selection against gene flow in capuchino seedeaters.

**A general framework of simulations used in the tests for statistical power.** Because no existing population genetic simulation software package has an explicit model for selection against gene flow, we designed a custom simulation framework to mimic such a scenario. We simulated a collection of genomic segments, some of which represented the genomic background, while others represented genomic regions under selection against gene flow. All simulations included 24 diploid individuals split evenly between two species, to mimic one of our pairwise comparisons. The genomic background segments were simulated using demographic parameters taken from Campagna *et al*. (2015)[6] (see details below), and segments modeling selection against gene flow were simulated with a deeper divergence time, while keeping the remaining demographic parameters the same. In this way, we mimicked a scenario of complete selection against gene flow during the interval between the deeper and shallower divergence times. We simulated a large collection of segments with a series of different divergence times and then applied our method to each of these segments using the background segments as flanking regions. Finally, for each divergence time, we recorded the proportion of segments simulated with that time for which our method estimated significant elevation in CC times (an estimate of the power of our test).

**Detailed description of simulations.** We simulated 450 separate 100 kb segments: 300 segments were simulated for the genomic background, and the remaining 150 segments were partitioned into five sets with various deep divergence times. The 300 genomic background segments were simulated using a divergence time of 43,000 generations, as inferred by Campagna *et al*. (2015)[6], and each of the other five sets was simulated with a divergence time in {100, 200, 300, 400, 500} x 1,000 generations. Simulations were carried out using SLiM[7], and followed similar guidelines to the simulations used to generate the training and test sets for the SVM (see **Section 4 in Supplementary Text**). In particular, we scaled down effective population sizes and divergence times by a factor of 100 to improve computational efficiency. Per-generation mutation and recombination rates were drawn from a uniform distribution in the intervals [1e-9, 1e-7] and [1e-6, 2e-6], respectively. We started with an initial population containing 144,500 individuals, and split this population into two sister species after T1 generations. The effective population size for each derived species was then sampled uniformly in the interval [1100, 2000], and each derived species was allowed to evolve for an additional T2 generations. T1 and T2 were set so that T1+T2 = 18,550, and T2 takes one of the following values: 430 (background), 1000, 2000, 3000, 4000, and 5000. Each simulation thus ran for a total of 18,550 generations before 24 haploid samples were sampled from each of the two derived species at “present time”.

**Analysis of simulated data.** We executed *ARGweaver* on each of the 450 simulated segments for 1000 MCMC iterations, and recorded the ARG sample generated in the 1000th iteration. We estimated a significance threshold for quantile difference in CC times, mimicking the approach we took in the genomic analysis. We partitioned the background segments into 150 pairs, and used their inferred ARGs to compute the quantile difference between CC times. Because the distribution of these quantile differences is expected to be symmetric around 0 (both segments were generated by the same model), we estimated its upper tail by examining the absolute values of these 150 quantile differences, and we used the tail of this distribution to determine a threshold (0.23) associated with a *p*-value of 0.02. Note that this threshold is slightly higher than the one we computed using actual genomic segments, likely because of the fact that we are not modeling linkage between a segment and its “flanking” region (see discussion below). Finally, each of the 150 segments simulated with deep divergence was randomly paired up with a different background segment as its flanking region. For each such pair, we used the two inferred ARGs to compute the quantile difference between CC times, as done in the genomic analysis, where values above 0.23 were considered statistically significant ($p<0.02$; see above).

**Results of power analysis.** We recorded for each of the five sets in our simulation experiment the number of sets (out of 30) for which the quantile difference was above the significance threshold of 0.23. We also recorded the number of sets for which the quantile difference attained its maximum value of 0.5. To provide an indication for the level of sequence differentiation observed in each set, we also computed average $F_{ST}$ values across the 30 segments simulated for each set. Results are summarized in **Supplementary Table S4**. The statistical power of our method is estimated to be 27% in the set simulated with divergence time set to 100,000 generations. However, the average $F_{ST}$ for segments in this set is less than the threshold value 0.2 used for determining the 25 $F_{ST}$ peaks we analyzed in this study. Thus, if differentiation in one of these peaks is solely attributed to selection against gene flow, then this would have had to start earlier than 100,000 generations ago to cause the observed level of $F_{ST}$. Importantly, in all of the sets that have average $F_{ST}$ above 0.2, the statistical power of our method is estimated to be larger than 90%. We conclude from this analysis that despite the recent divergence and large ancestral population sizes, if one or more of the 25 $F_{ST}$ peaks we analyzed was caused by selection against gene flow, our method based on elevated CC times should be able to detect it with high probability.

**Discussion related to power analysis.** The simulation experiments indicate reasonable statistical power to detect selection against gene flow in our setting, but this framework has two main limitations. The first limitation is that we model recombination only within each region, and we do not model linkage between regions (each region is simulated separately). This stems from a fundamental limitation of all existing simulation tools, which do not allow explicitly modeling selection against gene flow, or modeling changes in divergence times along a sequence. The lack of linkage eliminates the correlation that we observe in real genomic data between the CC times inferred in a region of interest and those inferred in its flanking regions. This correlation likely results in quantile differences that are somewhat closer to 0 than the ones we observed in simulations, as can be attested to by the slightly lower significance thresholds: 0.21 for a $p$-value of 0.01 in the genomic comparisons, and 0.23 for a $p$-value of 0.02 in the simulations. However, because these differences are quite small, and because we control the false positive rate in the simulations (by setting a higher threshold), we expect similarly high statistical power even when linkage is present. The second limitation of our simulations is that selection against gene flow was assumed to be very strong, leading to complete isolation at the time of initiation. Thus, these experiments provide a lower bound on the time that selection against gene flow started. If selection is weak enough to allow for some gene flow, then it would have had to start earlier than 200,000 generations ago to result in an average $F_{ST}$ above 0.2 and to allow detection by our method with high probability.

#### 3. Supplementary results and discussion for the ARG-based analysis.

**Species differentiation in local trees in $F_{ST}$ peaks (test 1 in Table 1).** Species enrichment scores were computed across the 25 $F_{ST}$ peaks for all five species, and a maximum value was recorded for every peak and species (**Supplementary Table S2**). Significance thresholds were determined separately for each species using the control set of 39,699 windows and an empirical *p*-value of 0.0001. A given genomic region is said to pass test 1 for certain species, if the region contains a 20 kb window with a significant enrichment score for that species. A total of 23 of the 25 peaks (92%) pass test 1 for at least one species, and seven of these (28%) pass the test for more than one species. To assess the significance of this result, we also examined a collection of 1,376 non-overlapping 500 kb blocks from the 576 scaffolds that do not contain $F_{ST}$ peaks (the same scaffolds used to compute empirical *p*-values; see **Methods**). Only seven of these 1,376 genomic blocks (0.5%) pass test 1, and this is always for a single species. This confirms the expectation of a strong association between high $F_{ST}$ values and differentiation between species in the local trees, as illustrated in the examples shown in **Supplementary Figure S27**. Interestingly, despite the fact that we used a separate threshold for each species, different species appear to contribute differently to divergence. The largest contribution comes from *S. nigrorufa*, for which 14 peaks pass test 1, and *S. melanogaster*, for which 8 peaks pass the test. On the other hand, only two peaks pass the test for *S. hypoxantha*, but its significance threshold was also the highest (4.3), which could suggest that regions associated with its divergence may fall outside the $F_{ST}$ peaks determined by Campagna *et al.* (2017)[1].

**Reduced RTH' in $F_{ST}$ peaks (test 2 in Table 1).** We computed minimum RTH' values for each of the 25 $F_{ST}$ peaks for the five species (**Supplementary Table S3**). Significance thresholds were determined separately for each species using the control set of 39,699 windows and an empirical *p*-value of 0.0001. A given genomic region is said to pass test 2 for certain species if the region contains a 20 kb window with a significantly low RTH' in that species. A total of 20 cases pass test 2, 18 of which coincide with cases that pass also test 1 (see **Table 1** for combined results). Thus, out of the 35 cases that pass test 1 (**Supplementary Table S2**), more than half also pass test 2 and are thus apparently associated with recent selective sweeps in the differentiated species. Two $F_{ST}$ peaks (on scaffolds 567 and 762) passed test 2 for two species, suggesting parallel or shared sweeps (see, for example, **Supplementary Figures S6** and **S7**). For comparison, only five of the 1,376 long genomic blocks selected for the control (0.4%) passed test 2, and this was always for a single species.

We note that some of the decrease in RTH' observed in $F_{ST}$ peaks is a direct result of the existence of species-enriched clades in the local trees in these regions. This is because enriched clades reduce the size, and thus the age, of clades that contain half the samples of the enriched species. To deal with this possible confounding factor, we computed for every local tree a relative time for the first coalescence of any 12 lineages (RT12) (from possibly different species). This relative age was obtained by taking the age of the youngest subtree with at least $n$=12 leaves, and dividing it by the age of the subtree that contained at least half the total number of samples. A reduced RT12 corresponds to a recent sweep affecting at least 12 haploid samples, which is the same number of samples required in the subtree defining RTH'. Average RT12 values were computed in 20 kb windows and a significance threshold was set using a relaxed empirical $p$-value of 0.001 based on the 39,699 windows selected for control. RT12, which should not a-priori be correlated with species enrichment scores, was significantly reduced ($p$<0.001) in eight of the 25 $F_{ST}$ peaks (32%), whereas only five of the 1,376 long genomic control blocks (0.4%) contained windows with RT12 below the same threshold. Together, these results suggest that much of the species differentiation observed in $F_{ST}$ peaks is a result of recent selective sweeps, and most of these are species-specific.

**Examining inter-species cross coalescence times in $F_{ST}$ peaks (as part of test 3 in Table 1).** We computed the distribution of normalized inter-species cross coalescence times (CC) for all 10 species pairs in all 24 $F_{ST}$ peaks and flanking regions (excluding the peak on scaffold 308 due to insufficient flanking sites; see **Supplementary Text 2**). A significance threshold of 0.21 was determined using the control set of 1,376 regions of length 500 kb and an empirical $p$-value of 0.01. **Supplementary Table S5** specifies quantile differences between the distribution in peaks and the distribution in the associated flanking regions and highlights cases that pass the significance threshold. Interestingly, in 43 out of the 58 cases (74%) where we see quantile differences larger than 0.21, this difference is a result of reduced CC times in a peak relative to its flanking regions. This is likely a consequence of selective sweeps, some of which contain individuals from multiple species, thus decreasing inter-species coalescence times (see, for example, **Supplementary Figures S31** and **S32**). Hence, on average, CC times are not elevated in $F_{ST}$ peaks, as observed by inspection of $d_{XY}$ statistics done by Campagna *et al.* (2017)[1] (see also **Supplementary Figure S2**). Nonetheless, there are six $F_{ST}$ peaks in which we observe significantly elevated CC times for at least one pair of species. However, elevated CC times could also be caused by other factors, as discussed in detail in the three sections below.

**Elevated cross coalescence times caused by shared selective sweeps.** In the $F_{ST}$ peak on scaffold 762, we observe significantly elevated CC times between either *S. nigrorufa* or *S. pileata* and each of the other three species (other than the pair *S. pileata* vs. *S. hypoxantha*, where we observe elevation below the significance threshold). However, both *S. nigrorufa* and *S. pileata* have significantly low RTH' in this peak and only *S. nigrorufa* is significantly enriched (**Supplementary Tables S2** and **S3**). Examining a representative local tree in this peak (**Supplementary Figure S31**), we observe a young clade (originating roughly 250,000 generations ago) that contains 35 haploid samples, out of which 21 are from *S. nigrorufa* and 13 samples from *S. pileata*. This, together with the enrichments observed in this peak, implies a recent selective sweep shared by the two species (possibly due to adaptive introgression; see **Discussion** and **Supplementary Figure S30**). Such a shared sweep could explain the observed elevation in CC times, because it reduces by nearly 67% the number of lineages from *S. nigrorufa* and *S. pileata* that are free to coalesce with lineages from the remaining three species in the time before the sweep began. We thus conclude that elevated CC times in this peak are likely a result of a recent shared sweep in *S. nigrorufa* and *S. pileata*, and not a result of selection against gene flow, which would not have led to low RTH' for the two species. This is why for test 3 we require that the species does not pass test 2.

**Elevated cross coalescence times unrelated to species differentiation.** In the first peak on scaffold 637 we observe elevated CC times between *S. hypoxantha, S. melanogaster*, and *S. palustris*, and none of these three species pass test 1 in this peak (**Supplementary Tables S2** and **S5**). Similarly, in the $F_{ST}$ peak on scaffold 766, we observe elevated CC times between *S. pileata* and *S. hypoxantha*, but none of these two species passes test 1 in the peak. We thus conclude that the elevated CC times observed in these two peaks is likely unrelated to species differentiation, and could be a result of other phenomena, such as balancing selection or ancestral polymorphism. This is why we require that a species pass test 1 in order to pass test 3.

**Three peaks pass test 3 in Table 1.** The three $F_{ST}$ peaks on scaffolds 252, 1635, and 1717 satisfy the conditions of test 3, since each of them has at least one species with significantly elevated CC times that passes test 1 and does not pass test 2. Below, we illustrate in detail the genealogical signatures observed in these peaks, which are consistent with selection against gene flow. **The peak on scaffold 252** exhibits a combination of different factors contributing to species differentiation, as described in detail in the section below. One of these factors appears to be early selection against gene flow, starting roughly 1 million generations ago, between *S. palustris* and the other four species. Note that the divergence time of ~45,000 generations inferred for the five species by Campagna *et al.* (2015)[6] reflects the time at which the genomes started diverging. However, local barriers to gene flow could indeed start long before that. **The peak on scaffold 1635** shows evidence of a deep clade enriched for *S. melanogaster* (elevated CC between *S. melanogaster* and *S. palustris*). Thus, early selection against gene flow (starting roughly 1 million generations ago) between *S. melanogaster* and the other four species likely explains its differentiation in this peak (**Supplementary Figure S6**). **The peak on scaffold 1717** is consistent with possible contribution to differentiation from selection against gene flow, but evidence is somewhat weaker. We find evidence of a deep clade enriched for *S. melanogaster* (elevated CC between *S. melanogaster* and either *S. hypoxantha* or *S. nigrorufa*). However, there are signs of a selective sweep shared by *S. hypoxantha* and *S. nigrorufa* in this region (**Supplementary Figure S32**), which could also contribute to the observed elevation in CC (see explanation above). Thus, in this case we conclude that if there was selection against gene flow for *S. melanogaster* in this region, it had a modest effect on sequence divergence.

**Genealogical patterns of differentiation in the ASIP locus.** The peak on scaffold 252, which is upstream of the gene encoding for the Agouti-signaling protein (ASIP) is particularly interesting, since divergence in this region is likely caused by a combination of various different factors. The species *S. melanogaster* appears to have undergone a very recent species-specific complete sweep, as suggested by the very high enrichment score (18.34) and very low RTH' (0.043) (**Supplementary Tables S2** and **S3**). The representative tree shown in **Supplementary Figure S5** has a clade whose age is approximately 10,000 generations, which contains 21 out of the 24 samples from this species and no samples from other species. The species *S. nigrorufa* appears to have undergone a more ancient sweep, indicated by its high enrichment score (7.46), larger RTH' (0.592) and the age of its enriched clade in the representative tree (approximately 200,000 generations). This divergence may have been maintained by selection against gene flow, but evidence for this is somewhat weak, since this species has higher divergence levels genome-wide, and its current geographic range has little overlap with other species ranges. The species *S. pileata* is most striking in this respect, since it has a very high enrichment score (8.81) and a high RTH' (0.967). This RTH' is, indeed, higher than what we expect in a genomic region with high species enrichment. The significantly elevated CC times between this species and three other species (**Supplementary Table S5**) provides additional evidence for prolonged separation. Moreover, the representative tree shown in **Supplementary Figure S5** contains a clade whose age is roughly a million generations, which contains 20 samples, 18 of which are from *S. pileata*. Such deep separation is likely maintained by selection against gene flow. A similar case can be made also for *S. palustris*, but since its enrichment score is much lower (3.19), it is unclear whether its deep separation is not primarily a result of the separation of the other three species.

#### 4. Supplementary methods of models for selective sweeps using machine learning

**Pairwise experiments.** We conducted a series of pairwise analyses of Southern Capuchino species to better understand the nature of the selective sweeps that affect scaffolds that contain $F_{ST}$ peaks. These experiments consisted of four classification tasks designed to compare the following types of sweeps:

1. Soft vs. hard sweeps. A hard sweep involves an increase in frequency of a newly arising beneficial mutation, together with its haplotype background. In a hard sweep scenario, there is almost no genetic variation left in the haplotype background associated with the beneficial mutation. A soft sweep involves an increase in frequency of a standing genetic variant, together with the associated haplotype backgrounds, when that variant becomes beneficial (for example due to an environmental change). In a soft sweep scenario, there is some genetic variation left in the haplotype background associated with the beneficial mutation.
2. Recent vs. ancestral sweeps. A recent sweep occurs after the split of the ancestral population into five capuchino species while an ancestral sweep involves a beneficial mutation that occurs in the ancestral population.
3. Complete vs. partial sweeps. A complete sweep involves a beneficial mutation that increases in frequency together with its haplotype background until it reaches fixation, while a partial sweep involves an increase in frequency of a beneficial mutation without the beneficial mutation reaching fixation.
4. Species-specific vs. parallel sweeps. A parallel sweep is defined as two independent sweeps occurring in the two derived species.

**Expanded analysis**. Based on the binary outcomes of these pairwise experiments, we designed an expanded analysis involving all five species. The binary outcomes of the pairwise-based experiments pointed to soft sweeps being the predominant mode of adaptation in these capuchino species as shown using the machine learning classifier, ARG-based summary statistics, local genealogies extracted from the ARG, and homozygosity-based statistics. Furthermore, these soft sweeps tend to mostly be complete and recent. We thus constructed an expanded analysis of all five species to identify species-specific complete soft sweeps, allowing for a sweep to occur in any one of the species.

**Demographic parameters used when generating simulated data with SLiM.** All simulations start with an initial population containing 144,500 individuals. This size reflects the effective size inferred by Campagna *et al.* (2015)[6] for the population ancestral to all five species and sister to the outgroup S. *bouvreuil*, scaled down by a factor of 100. Scaling down the size of the simulated population(s) reduces the running time of the simulations, but requires scaling all other parameters appropriately. Thus, all time durations were also scaled down by a factor of 100 relative to the values inferred by Campagna *et al.* (2015)[6], and the mutation and recombination rates were scaled up by the same factor, relative to the values expected in these bird species. Consequently, per-generation mutation and recombination rates were drawn from a uniform distribution in the intervals [1e-9, 1e-7] and [1e-6, 2e-6], respectively. The ancestral population was allowed to evolve for 18,120 generations, and then simultaneously split into either two or five derived species (depending on whether we did a species-pair analysis or an expanded analysis of all five species). The effective population size for each derived species was sampled uniformly in the interval [1100, 2000], and each derived species was allowed to evolve for an additional 430 generations, reflecting the divergence time and range of effective population sizes inferred by Campagna *et al.* (2015)[6] for the five capuchino species (scaled down by a factor of 100). Each simulation thus ran for a total of 18,550 generations, reflecting the time since divergence from *S. bouvreuil* [6], before 24 haploid samples were sampled from each derived species. Finally, to demonstrate how well the simulations fit the empirical data at hand, we applied PCA to the summary statistics extracted from both the empirical data and the simulations based on the demographic model inferred from RAD-seq data. We find that the top two principal components for the two data sets largely overlap (**Supplementary Figure S33**), suggesting that the inferred demographic model fits the genomic data reasonably well.

**Selection parameters used when generating simulated data with SLiM.** When simulating data under various models for selective sweeps, a single focal site was designated as the one affected by positive selection. The position of this site within the 50 kb simulated sequence was sampled uniformly at random in the interval [20,000, 30,000], that is, in the middle 10 kb of the simulated region. The scaled selection coefficient for this site was drawn uniformly from the interval [0.75, 5]. The onset of selection was drawn uniformly between generations 280 and 330 after the species split, reflecting a sweep that started roughly 10,000-15,000 generations ago (considering the 100-fold scaling ; see above). For data simulated under hard sweeps, the onset of selection is defined as the time at which the beneficial mutation occurs in the focal site, and for data simulated under soft sweeps, it is defined as the time that the focal site is switched from neutral drift to positive selection. Furthermore, for data simulated under soft sweeps, we introduced a prior on the derived allele frequency of the focal site at the onset of selection, such that it ranged between 0.01 and 0.1. For data simulated under partial sweeps, the focal site was set to switch back to neutral drift when it reached a derived allele frequency drawn uniformly from interval [0.1, 0.99]. In all of the cases above, selection was set to act only in one of the derived species. For data simulated under ancestral sweeps, the onset time of selection was drawn uniformly in the 20 generations immediately before the species split, reflecting a sweep that started roughly 44,000 generations ago.

**Validation of inferred sweeps using ARG-based measures.** The inferred soft and hard sweeps were validated using summary statistics extracted from the ARG inferred by *ARGweaver*. In particular, we explored two measures relating to the within-species TMRCA, which are all expected to be reduced in species-specific selective sweeps (see also **ARG-based measures associated with sweeps** in Methods): (1) TMRCA - the age of the youngest clade that contains all 24 haploid samples from a given species and (2) RTH - the age of the youngest clade that contains at least half of the haploid samples from a given species (TMRCAH), divided by the TMRCA of that species. The distribution of these values was recorded for regions classified by the linear SVM into each of the three selection categories—soft sweeps, hard sweeps, and neutral—to examine observed differences. As additional validation, we used representative local trees extracted from the ARG in regions predicted to undergo sweeps by either the species-pair analyses or the expanded five species analysis. Within a given 10 kb window classified as a sweep in a given species, we extract as a representative tree the local tree with the lowest RTH.

**Validation of soft versus hard sweeps using homozygosity statistics.** To differentiate soft from hard sweeps, we used the haplotype homozygosity statistics, $H_1$, $H_2$, and $H_{12}$ (ref. [8]). The haplotype homozygosity, $H_1$, is defined as the probability of observing a completely homozygous haplotype (50 kb long), which is computed by taking the sum of squares of all haplotype frequencies. Similarly, $H_2$ is the haplotype homozygosity computed by ignoring the most frequent haplotype, and $H_{12}$ is the haplotype homozygosity computed by combining the two most frequent haplotypes. In hard sweeps, one haplotype rises to much higher frequency than all others, so $H_1$ and $H_{12}$ are both expected to be high, but the ratio $\frac{H_2}{H_1}$ is expected to be low. However, in soft sweeps, the ratio $\frac{H_2}{H_1}$ is expected to be much higher, and $H_{12}$ is typically lower (depending on how "soft" the sweep is). Thus, the two statistics $\frac{H_2}{H_1}$ and $H_{12}$ provide effective separation between the two types of sweeps, as demonstrated in ref. [8].

**High-confidence prediction of sweeps.** For high-confidence predictions, we considered those associated with a normalized probability 0.95 or higher. In addition we assigned each prediction an empirical *p*-value based on a set of 10,000 regions simulated under neutral evolution. Specifically, for each trained classifier and each non-neutral class assignment, we obtained a null distribution of prediction probabilities based on the application of the classifier to the neutral regions. Then, the *p*-value associated with each non-neutral prediction was taken to be the relative frequency of values in the corresponding null distribution that equal or exceed the given prediction probability. Finally, the set of *p*-values for each non-neutral class of each classifier were corrected using the false-discovery-rate (FDR) method of Benjamini & Hochberg[77], and we confirmed that calls with prediction probability above 0.95 were associated with an FDR-corrected *p*-value below 0.05.

#### 5. Robustness of the selective sweep classifier

We evaluated the robustness of our approach by considering alternative demographic parameters (such as ancestral Ne, derived species Ne, and divergence time) as well as alternative parameters for recombination rate, mutation rate, selection coefficients, and gene conversion. Our general approach was to apply the previously trained SVM to simulated data sets (with 200 replicates per model condition) that were generated under various alternative parameter settings, with values well outside the ones used for training. In this way, we tested our model on data sets for which it was deliberately misspecified, and we could systematically measure the degree to which prediction performance was consequently degraded. For simplicity, we focused on the classification task of differentiating soft and hard sweeps in a two-species setting (see above under **Pairwise experiments**), which we believe is reasonably representative of all of our tasks.

Our initial model used simulations starting with an ancestral population size of 14,450,000, as inferred for the population ancestral to all five derived species. In the misspecified datasets, we varied the ancestral population size according to the following alternative parameter values: 7,500,000, 10,000,000, 12,500,000, and 17,500,000. **Supplementary Figure S8** reports the performance difference in confusion matrices between the misspecified dataset relative to the correctly specified test set. We observed minimal performance difference (<5%) in the classification accuracy for the different classes and in cases of misclassification. Furthermore, in our trained model, the effective population size for each derived species was sampled uniformly in the interval [110000, 200000]. Using a similar approach, we varied the derived species population size according to the following alternative parameter values: 75,000, 100,000, 225,000, and 250,000. **Supplementary Figure S9** reports the performance difference in confusion matrices for each misspecified experiment. For the misspecified experiments with the derived species population size values of 100,000, 225,000, and 250,000, the performance difference between the misspecified dataset relative to the correctly specified test set ranged between -12% and +6%. For the experiment with the derived species population size value of 75,000, the performance difference was still small, but ranged between -22% to +5%, with the highest error resulting in the misclassification of soft sweeps as hard, which would not weaken our main conclusion that sweeps are predominantly soft. Overall, these results suggest that our prediction methods are generally not highly sensitive to the derived species population size, but when these population sizes become too small, random effects dominate the simulations, contributing to an increase in error.

In the simulations used to train our model, we assumed that the sampled populations diverged 43,000 generations ago. However, divergence might actually be deeper, especially if it wasn't instantaneous and involved some gene flow. To examine the possible effect of our assumption on divergence time we simulated additional sets using deeper divergence times of 105,000, 155,000, and 355,000. The only classification error whose rate increased in these data sets by more than 8% is the classification of soft sweeps as hard sweeps (**Supplementary Figure S10**). This rate of this classification error grows by 12% when the actual divergence time is more than twice that assumed in training (105,000), and grows by 52% when the actual divergence time is eight times larger (355,000), which is a fairly extreme case. Importantly, this type of classification error would not weaken our main conclusion that the sweeps are predominantly soft.

Furthermore, we explored two alternative values of the selection coefficient, 0.005 and 0.075, which fall outside the range [0.0075, 0.05] in which we sample selection coefficients when training the SVM. Results are reported in **Supplementary Figure S11**. In general, our classifier performed very well under these conditions, but it did appear to have limited sensitivity when selection is weak (in this case, due to misclassification of hard as soft sweeps).

Although our trained model accommodated variation in mutation and recombination rates, we created multiple misspecified datasets varying either the per-generation mutation or recombination rate such that the alternative values used are outside the range of simulations in the training set. We used the following alternative parameter values: $\boldsymbol{\mu}$ = 3e-09, 4e-09, or 5e-09 (compared to training values $\boldsymbol{\mu}$ sampled from [1e-11, 1e-09]) and $\boldsymbol{\rho}$ = 4e-08, 5e-09, and 2e-09 (compared to training values $\boldsymbol{\rho}$ sampled from [1e-08, 2e-08]). Results are reported in **Supplementary Figures S12** and **S13**. The model seems highly robust to changes in mutation rate. It is somewhat more sensitive to changes in recombination rate, particularly when it decreases from 4e-8 to 2e-09, presumably because the LD-based signatures of sweeps are strongly distorted at very low recombination rates. In general, however, we found that our classifier was not highly sensitive to reasonable departures from the parameter values assumed during training.

In addition, we carried out an analysis to see the effect of gene conversion, given that it is thought to occur at higher rates than crossover, which is the only form of recombination simulated in our training and test sets. In particular, it is possible that gene conversion could tend to "soften" a hard sweep, by causing the haplotype backgrounds of selected alleles to appear more heterogeneous. To test whether this phenomenon could have substantially influenced our predictions, we included gene conversion in our robustness analysis, by simulating it within 50kb segments using SLiM. We set the two relevant SLiM parameters as follows: 1) nonCrossoverFraction, which describes the fraction of recombination events that do not result in crossover, was set to 0.8; and 2) the mean length of gene conversion tracts, which are assumed to be geometrically distributed, was set to 5,000bp. Results are reported in **Supplementary Figure S14**, and suggest that gene conversion had little effect on the overall performance of our classifier, and had little impact in particular on the misclassification of hard sweeps as soft.

Finally, it has been shown[9] that the flanking regions of hard sweeps can be mistakenly identified as soft sweeps (referred to as "soft shoulders"). We addressed the possibility of "soft shoulders" by simulating a set of 50kb regions in which the middle window (20kb-30kb) is not under selection but the flanking part of each region (0kb-10kb or 40kb-50kb) is under selection (soft or hard sweep). We found that our classifier was generally able to correctly predict these middle windows as neutral, with an average accuracy of 77% (**Supplementary Table S7**). Thus, it appears that our model is generally able to approximately pinpoint the location of the causal mutation for a sweep, without being confused by the patterns at the shoulders of sweeps. Furthermore, we re-examined the high-confidence soft sweep predictions that our classifier makes, and only 3% of them fall in a flanking region (20kb upstream and downstream) of a predicted hard sweep. Therefore, it seems unlikely that our predictions are strongly influenced by miscalling the flanking regions of hard sweeps as soft sweeps.

#### Supplementary Tables

**Supplementary Table S1:** The 25 $F_{ST}$ peaks considered in our analysis. These regions were determined by Campagna *et al.* (2017)[1] as stretches of two or more non-overlapping 25-kb windows with average $F_{ST}$ above 0.2 in at least one of the ten pairwise comparisons and at least one variant site with $F_{ST}$ above 0.85 (windows with fewer than ten variant sites were not considered; see Campagna *et al.* (2017)[1] for more details). The peak id is based on the sequence scaffold it belongs to. Sequence scaffolds are also mapped to chromosomes of the zebra finch genome. Nearby melanogenesis genes are indicated as in Table 1 of Campagna *et al.* (2017)[1].

| peak id | scaffold | start position | end position | length (kb) | zebra finch chromosome | melanogenesis genes |
|---|---|---|---|---|---|---|
| Scaffold59 | 59 | 5750000 | 5835000 | 85 | 15 | |
| Scaffold118 | 118 | 7165000 | 7190000 | 25 | 2 | |
| Scaffold252 | 252 | 420000 | 510000 | 90 | 20 | *ASIP* |
| Scaffold257a | 257 | 5800000 | 6170000 | 370 | Z | |
| Scaffold257b | 257 | 21245000 | 21745000 | 500 | Z | *TYRP1* |
| Scaffold257c | 257 | 23975000 | 24815000 | 840 | Z | *MLANA* |
| Scaffold257d | 257 | 28665000 | 28930000 | 265 | Z | |
| Scaffold257e | 257 | 31310000 | 31355000 | 45 | Z | |
| Scaffold263 | 263 | 20000 | 555000 | 535 | Z | *MYO5A* |
| Scaffold308 | 308 | 55000 | 195000 | 140 | unknown | |
| Scaffold404a | 404 | 5050000 | 5815000 | 765 | Z | *SLC45A2* |
| Scaffold404b | 404 | 10730000 | 10925000 | 195 | Z | |
| Scaffold412 | 412 | 3390000 | 3595000 | 205 | 1A | *KITL* |
| Scaffold430 | 430 | 10995000 | 11080000 | 85 | 1 | *OCA2* |
| Scaffold567 | 567 | 2515000 | 2775000 | 260 | 2 | |
| Scaffold579 | 579 | 335000 | 1005000 | 670 | 1 | |
| Scaffold637a | 637 | 6015000 | 6315000 | 300 | Z | |
| Scaffold637b | 637 | 6850000 | 6890000 | 40 | Z | |
| Scaffold762 | 762 | 1660000 | 1695000 | 35 | 11 | |
| Scaffold766 | 766 | 2000000 | 2065000 | 65 | 4 | |
| Scaffold791 | 791 | 9905000 | 9955000 | 50 | 1 | *DCT* or *TYR* |
| Scaffold1635 | 1635 | 3710000 | 3740000 | 30 | 6 | |
| Scaffold1717 | 1717 | 930000 | 960000 | 30 | 4 | *CAMK2D* |
| Scaffold1954 | 1954 | 2825000 | 2900000 | 75 | 5 | |
| Scaffold3622 | 3622 | 975000 | 1360000 | 385 | 1 | |

**Supplementary Table S2:** Maximum enrichment scores for the five *Sporophila* species in each of the 25 $F_{ST}$ peaks. Gray shade indicates scores that exceed thresholds associated with an empirical *p*-value of 0.0001 (test 1; gray cells in Table 1, see **Methods**). Thresholds were determined separately for every species: *S. hypoxantha* (hypox) 4.3; *S. melanogaster* (mel) 3.7; *S. nigrorufa* (nig) 4.1; *S. palustris* (pal) 3.1; *S. pileata* (pil) 3.4. Scores that exceed the maximum enrichment score observed in the control set (5.46) are indicated in bold.

| peak id | length (kb) | hypox | mel | nig | pal | pil |
|---|---|---|---|---|---|---|
| Scaffold59 | 85 | 4.39 | 2.39 | 1.97 | 1.78 | 1.97 |
| Scaffold118 | 25 | 1.97 | 1.96 | 4.9 | 2.26 | 1.84 |
| Scaffold252 | 90 | 3.01 | **18.34** | **7.46** | 3.19 | **8.81** |
| Scaffold257a | 370 | 2.29 | 2.62 | **7.39** | 2.29 | 2.56 |
| Scaffold257b | 500 | 2.16 | **10.79** | 4.09 | 2.69 | 2.5 |
| Scaffold257c | 840 | 3.83 | 2.93 | 3.88 | **7.9** | 2.92 |
| Scaffold257d | 265 | 2.23 | **6.98** | 2.72 | 2.71 | 2.59 |
| Scaffold257e | 45 | 2.33 | 2.4 | 2.79 | 2.37 | 2.3 |
| Scaffold263 | 535 | 2.14 | 2.38 | 4.24 | 2.05 | 2.75 |
| Scaffold308 | 140 | 2.2 | 4.43 | **6.14** | 2.04 | 4.95 |
| Scaffold404a | 765 | 3.64 | 4.2 | **6.36** | 3.21 | 4.55 |
| Scaffold404b | 195 | 2.47 | 2.41 | **10.07** | 2.09 | 2.89 |
| Scaffold412 | 205 | 2.58 | 3.41 | **7.73** | 2.24 | 5.28 |
| Scaffold430 | 85 | 2.29 | 4.3 | 3.16 | 2.19 | 4.61 |
| Scaffold567 | 260 | 2.29 | 2.16 | **6.81** | 2.04 | 5.39 |
| Scaffold579 | 670 | 2.84 | 3.45 | 5.06 | 2.61 | 2.69 |
| Scaffold637a | 300 | 2.08 | 2.27 | **6.03** | 2.15 | 2.9 |
| Scaffold637b | 40 | 1.97 | 1.83 | 4.05 | 2.38 | 2.38 |
| Scaffold762 | 35 | 2.3 | 2.36 | **9.37** | 2.3 | 2.9 |
| Scaffold766 | 65 | 2.45 | 1.98 | 2.35 | **7.75** | 2.01 |
| Scaffold791 | 50 | 2.1 | 1.92 | 2.06 | 5.18 | 2.02 |
| Scaffold1635 | 30 | 2.45 | 4.96 | 1.93 | 2.05 | 2.28 |
| Scaffold1717 | 30 | 4.63 | 3.7 | 2.77 | 1.75 | 2.51 |
| Scaffold1954 | 75 | 1.94 | 2.28 | 4.39 | 2.36 | 1.98 |
| Scaffold3622 | 385 | 3.24 | 3.48 | **6.32** | 2.08 | 2.3 |

**Supplementary Table S3:** Minimum RTH' values for the five *Sporophila* species and minimum RT12 in each of the 25 $F_{ST}$ peaks. Gray shade indicates scores below the significance thresholds (test 2; red circles in Table 1, See **Methods**). Thresholds were determined separately for every species based on an empirical *p*-value of 0.0001: *S. hypoxantha* (hypox) 0.2; *S. melanogaster* (mel) 0.3; *S. nigrorufa* (nig) 0.5; *S. palustris* (pal) 0.5; *S. pileata* (pil) 0.5. RTH' values below the minimum value observed in the control set (0.188) are indicated in bold. For RT12, a separate threshold of 0.03 was used, based on a relaxed empirical *p*-value of 0.001.

| peak id | length (kb) | RTH' hypox | RTH' mel | RTH' nig | RTH' pal | RTH' pil | RT12 |
|---|---|---|---|---|---|---|---|
| Scaffold59 | 85 | 0.38 | 0.84 | 0.79 | 0.90 | 0.95 | 0.064 |
| Scaffold118 | 25 | 0.60 | 0.36 | 0.50 | 0.71 | 0.93 | 0.041 |
| Scaffold252 | 90 | 0.86 | **0.04** | 0.59 | 0.83 | 0.97 | 0.011 |
| Scaffold257a | 370 | 0.81 | 0.72 | **0.17** | 0.83 | 0.80 | 0.048 |
| Scaffold257b | 500 | 0.82 | 0.27 | 0.59 | 0.82 | 0.75 | 0.081 |
| Scaffold257c | 840 | 0.80 | 0.63 | 0.84 | **0.04** | 0.74 | 0.016 |
| Scaffold257d | 265 | 0.89 | **0.17** | 0.83 | 0.84 | 0.76 | 0.044 |
| Scaffold257e | 45 | 0.90 | 0.84 | 0.83 | 0.81 | 0.92 | 0.106 |
| Scaffold263 | 535 | 0.87 | 0.89 | 0.79 | 0.92 | 0.88 | 0.134 |
| Scaffold308 | 140 | 0.92 | 0.68 | 0.48 | 0.90 | 0.71 | 0.114 |
| Scaffold404a | 765 | 0.50 | 0.84 | 0.30 | 0.48 | 0.63 | 0.047 |
| Scaffold404b | 195 | 0.82 | 0.85 | **0.08** | 0.91 | 0.75 | 0.023 |
| Scaffold412 | 205 | 0.82 | 0.79 | 0.43 | 0.88 | 0.55 | 0.059 |
| Scaffold430 | 85 | 0.93 | 0.33 | 0.92 | 1.00 | 0.39 | 0.047 |
| Scaffold567 | 260 | 0.88 | 0.81 | 0.35 | 0.94 | 0.25 | 0.028 |
| Scaffold579 | 670 | 0.72 | 0.38 | 0.37 | 0.60 | 0.86 | 0.033 |
| Scaffold637a | 300 | 0.82 | 0.85 | 0.62 | 0.82 | 0.82 | 0.104 |
| Scaffold637b | 40 | 0.97 | 0.90 | 0.63 | 0.95 | 0.86 | 0.105 |
| Scaffold762 | 35 | 0.92 | 0.87 | **0.05** | 0.99 | 0.45 | 0.022 |
| Scaffold766 | 65 | 0.92 | 0.71 | 0.96 | **0.06** | 0.93 | 0.024 |
| Scaffold791 | 50 | 0.80 | 0.85 | 0.92 | 0.45 | 0.92 | 0.119 |
| Scaffold1635 | 30 | 0.99 | 0.66 | 0.96 | 0.99 | 1.09 | 0.117 |
| Scaffold1717 | 30 | 0.61 | 0.79 | 0.80 | 0.97 | 0.89 | 0.023 |
| Scaffold1954 | 75 | 0.86 | 0.74 | 0.63 | 0.79 | 0.98 | 0.112 |
| Scaffold3622 | 385 | 0.35 | 0.27 | 0.63 | 0.54 | 0.78 | 0.017 |

**Supplementary Table S4:** Results of simulation experiments for assessing statistical power of our ARG-based method for detecting selection against gene flow. To assess the statistical power of our method for detecting selection against gene flow, we considered five different values for the divergence time, and for each value we simulated 30 segments of length 100 kb. An additional set of 300 background segments was simulated to compute a threshold corresponding to an empirical *p*-value of 0.02, and to act as flanking regions to the remaining segments. The background regions were simulated with divergence time of 43,000 generations, as inferred by Campagna *et al.* (2015)[6] (**Supplementary Text**). For each set, we measured the average $F_{ST}$, the fraction of segments for which we measure significantly elevated cross-coalescence times in inferred ARGs (quantile difference at least 0.23), and the fraction of segments for which the quantile difference attained a maximum value of 0.5.

| Simulated divergence time (generations) | Average FST | Segments with quantile diff. ≥ 0.23 | Segments with quantile diff. = 0.5 |
|---|---|---|---|
| (background) 43,000 | 0.07 | 1.3% | 0.7% |
| 100,000 | 0.16 | 26.7% | 0% |
| 200,000 | 0.27 | 93.3% | 0% |
| 300,000 | 0.33 | 93.3% | 6.7% |
| 400,000 | 0.38 | 96.7% | 33.3% |
| 500,000 | 0.41 | 100% | 53.3% |

**Supplementary Table S5:** Quantile differences between the distribution of normalized recent cross-coalescence times in $F_{ST}$ peaks relative to flanking regions (**Supplementary Text**). The quantile difference is defined as the difference between ½ and the quantile in one distribution associated with the median value of the second distribution. The quantile difference is computed in both directions ($F_{ST}$ peaks versus flanking region and vice versa), and the maximum value is recorded for each of the 10 species pairs in 24 $F_{ST}$ peaks, excluding the one on scaffold 308, which had insufficient flanking sites. The sign indicates whether cross-coalescence times in divergence peaks are elevated (+) or reduced (-). Quantile differences that exceed 0.21 in absolute value correspond to an empirical *p*-value smaller than 0.01 (based on an analysis of scaffolds that do not contain $F_{ST}$ peaks), and are highlighted in bold. Significant positive values are further highlighted with gray background, and are used as one of the three conditions for test 3 in **Table 1** (see **Methods)**.

| | length | hypox | | | | mel | | | nig | | pal |
|---|---|---|---|---|---|---|---|---|---|---|---|
| **peak id** | **(kb)** | **mel** | **nig** | **pal** | **pil** | **nig** | **pal** | **pil** | **pal** | **pil** | **pil** |
| Scaffold59 | 85 | 0.16 | **-0.22** | **-0.38** | -0.05 | -0.01 | 0.06 | -0.17 | **-0.24** | -0.04 | -0.05 |
| Scaffold118 | 25 | **-0.47** | -0.03 | **-0.46** | **-0.26** | -0.10 | **-0.50** | **-0.44** | **-0.29** | **-0.49** | **-0.45** |
| Scaffold252 | 90 | -0.06 | -0.08 | -0.13 | **0.23** | -0.01 | -0.09 | **0.42** | 0.10 | 0.14 | **0.27** |
| Scaffold257a | 370 | -0.04 | -0.16 | -0.02 | -0.08 | -0.03 | -0.15 | -0.08 | -0.11 | -0.10 | -0.08 |
| Scaffold257b | 500 | -0.11 | -0.19 | -0.05 | -0.10 | -0.09 | -0.12 | -0.12 | -0.03 | -0.12 | -0.03 |
| Scaffold257c | 840 | -0.09 | **-0.35** | -0.04 | **-0.18** | -0.08 | -0.14 | -0.20 | -0.19 | 0.09 | **-0.34** |
| Scaffold257d | 265 | -0.09 | -0.09 | -0.07 | -0.04 | 0.08 | -0.04 | -0.02 | 0.13 | -0.05 | 0.10 |
| Scaffold257e | 45 | **-0.26** | -0.07 | -0.01 | -0.05 | -0.01 | **-0.35** | -0.12 | 0.13 | 0.11 | 0.19 |
| Scaffold263 | 535 | 0.18 | -0.12 | -0.12 | -0.17 | -0.07 | 0.12 | 0.15 | -0.03 | -0.03 | -0.16 |
| Scaffold308 | 140 | - not enough flanking sites in sequence scaffold - | | | | | | | | | |
| Scaffold404a | 765 | -0.12 | **-0.25** | -0.08 | -0.08 | -0.07 | **-0.31** | -0.10 | 0.00 | -0.03 | -0.09 |
| Scaffold404b | 195 | **-0.21** | **-0.29** | **-0.26** | -0.09 | -0.15 | -0.15 | -0.10 | -0.11 | 0.07 | -0.14 |
| Scaffold412 | 205 | -0.04 | **-0.24** | -0.17 | -0.15 | 0.14 | -0.16 | -0.05 | 0.09 | -0.18 | -0.09 |
| Scaffold430 | 85 | -0.05 | 0.13 | -0.07 | 0.14 | -0.02 | 0.14 | **-0.39** | -0.08 | 0.18 | 0.10 |
| Scaffold567 | 260 | -0.14 | -0.18 | -0.16 | -0.16 | -0.10 | -0.13 | -0.14 | 0.05 | -0.18 | -0.19 |
| Scaffold579 | 670 | -0.12 | -0.15 | -0.18 | **-0.38** | -0.09 | **-0.33** | **-0.41** | -0.17 | -0.20 | -0.13 |
| Scaffold637a | 300 | **0.22** | 0.05 | **0.24** | 0.03 | -0.02 | 0.05 | 0.00 | 0.05 | 0.08 | -0.04 |
| Scaffold637b | 40 | **-0.29** | -0.17 | -0.02 | -0.09 | **-0.32** | **-0.22** | -0.04 | -0.05 | -0.16 | -0.01 |
| Scaffold762 | 35 | -0.01 | **0.23** | -0.10 | 0.14 | **0.32** | 0.15 | **0.29** | **0.26** | **-0.50** | **0.23** |
| Scaffold766 | 65 | -0.04 | -0.06 | 0.08 | **0.21** | -0.08 | **-0.48** | **-0.20** | 0.09 | 0.10 | **-0.41** |
| Scaffold791 | 50 | -0.10 | -0.01 | -0.08 | -0.12 | -0.08 | **-0.27** | -0.09 | -0.17 | 0.13 | -0.13 |
| Scaffold1635 | 30 | 0.13 | 0.05 | 0.06 | 0.00 | **0.30** | 0.20 | **-0.34** | 0.04 | -0.12 | **0.25** |
| Scaffold1717 | 30 | **0.37** | **-0.47** | **-0.42** | **-0.23** | **0.32** | -0.16 | -0.08 | **-0.38** | -0.13 | **-0.27** |
| Scaffold1954 | 75 | -0.06 | -0.10 | **-0.24** | 0.07 | 0.17 | -0.17 | -0.11 | 0.17 | -0.06 | -0.03 |
| Scaffold3622 | 385 | **-0.42** | 0.06 | **-0.28** | **-0.43** | 0.07 | **-0.40** | **-0.22** | 0.07 | 0.03 | 0.04 |

**Supplementary Table S6:** Confusion matrix for the classification task for soft versus hard sweeps in a two-species analysis. The classification task involves five classes: (1) neutral, (2-3) soft sweep in species #1 or #2, and (4-5) hard sweep in species #1 or #2. A five-way linear SVM classifier was trained using a training set of 40,000 simulated regions of length 50 kb (8,000 per class; see **Methods**). The classifier was then tested on a separate set comprised of 5,000 regions (1,000 per class). The cell in row *i* and column *j* of the confusion matrix reports the fraction of test regions simulated for class *i*, which the SVM assigned to class *j*. Thus, the values on the diagonal (in bold) represent the classification accuracy for the different classes, and off-diagonal entries correspond to cases of mis-classification. The average classification accuracy for this task (across all cases) is 92.84%, and the most common mis-classifications are: (1) soft sweeps being mis-classified as hard sweeps in the same species (9.15%), and (2) hard sweeps being mis-classified as soft sweeps in the same species (6.1%).

| | **Neutral** | **Soft (#1)** | **Soft (#2)** | **Hard (#1)** | **Hard (#2)** |
|---|---|---|---|---|---|
| **Neutral** | **98.70%** | 0.70% | 0.60% | 0.00% | 0.00% |
| **Soft (#1)** | 1.00% | **89.70%** | 0.00% | 9.30% | 0.00% |
| **Soft (#2)** | 2.00% | 0.20% | **88.80%** | 0.00% | 9.00% |
| **Hard (#1)** | 0.40% | 5.70% | 0.10% | **93.80%** | 0.00% |
| **Hard (#2)** | 0.20% | 0.00% | 6.50% | 0.10% | **93.20%** |

**Supplementary Table S7:** Confusion matrix for the robustness analysis involving simulating a set of 50kb regions where the middle window (20kb-30kb) is not under selection but the flanking part of each region (0kb-10kb or 40kb-50kb) is under selection (i.e. soft or hard sweep) in a two-species analysis. The classification task involves five classes: (1) neutral, (2-3) soft sweep in species #1 or #2, and (4-5) hard sweep in species #1 or #2. A five-way linear SVM classifier was trained using a training set of 40,000 simulated regions of length 50 kb (8,000 per class; see **Methods**). The classifier was then tested on a separate set comprised of 800 regions (200 per neutral class with a flanking soft or hard sweep in species #1 or #2). The cell in row *i* and column *j* of the confusion matrix reports the fraction of misspecified regions simulated for class *i*, which the SVM assigned to class *j*. The values in the 1st column (in bold) represent the classification accuracy (middle window) for the different neutral cases while the other column entries correspond to cases of mis-classification. The average classification accuracy for this task (across all cases) is 77%, and the most common mis-classification is neutral with a flanking hard sweep being mis-classified as a hard sweep (27-31%).

| | **Neutral** | **Soft (#1)** | **Soft (#2)** | **Hard (#1)** | **Hard (#2)** |
|---|---|---|---|---|---|
| **Neutral with flanking soft in #1** | **94%** | 3% | 1% | 3% | 0% |
| **Neutral with flanking soft in #2** | **92%** | 0% | 7% | 0% | 2% |
| **Neutral with flanking hard in #1** | **58%** | 12% | 0% | 31% | 0% |
| **Neutral with flanking hard in #2** | **63%** | 0% | 11% | 0% | 27% |

**Supplementary Table S8:** Numbers of soft and hard sweeps predicted across the four scaffolds (252, 412, 404, and 1717) that contain the top $F_{ST}$ peaks and harbor known pigmentation-related genes. Four separate species-pair analyses were executed, with the classification task involving five classes: (1) neutral, (2-3) soft sweep in species #1 or #2, and (4-5) hard sweep in species #1 or #2. A five-way linear SVM classifier was trained on simulated data and then applied to the genomic data using a sliding-window approach that classified each 10-kb window into one of the five classes (see **Methods**). The table reports the number of windows classified into the two "soft sweep" classes (2$^{nd}$ column) and into the two "hard sweep" classes (3$^{rd}$ column). The numbers of predictions associated with normalized probability 0.95 or higher are given in the 4$^{th}$ and 5$^{th}$ columns (FDR-adjusted *p*-value < 0.05; see **Section 4 in Supplementary Text**). Results suggest widespread species-specific soft sweeps across these four scaffolds.

| Pairwise analysis | # soft sweeps | # hard sweeps | # soft sweeps (Prob > 0.95) | # hard sweeps (Prob > 0.95) |
|---|---|---|---|---|
| **mel-nig** | 154 | 33 | 28 | 3 |
| **nig-pil** | 118 | 31 | 10 | 4 |
| **pil-pal** | 115 | 11 | 14 | 2 |
| **hypox-mel** | 158 | 13 | 17 | 1 |

**Supplementary Table S9:** Numbers of soft and hard sweeps predicted across all 19 scaffolds that contain an $F_{ST}$ peak. Four separate species-pair analyses were executed, with the classification task involving five classes: (1) neutral, (2-3) soft sweep in species #1 or #2, and (4-5) hard sweep in species #1 or #2. A five-way linear SVM classifier was trained on simulated data and then applied to the genomic data using a sliding window approach that classified each 10 kb window into one of the five classes (see **Methods**). The table reports the number of windows classified into each of the four non-neutral classes (3$^{rd}$ and 4$^{th}$ columns). The numbers of predictions associated with normalized probability 0.95 or higher are given in the 5$^{th}$ and 6$^{th}$ columns (FDR-adjusted *p*-value < 0.05). Results suggest widespread species-specific soft sweeps across these 19 scaffolds.

| Pairwise analysis | Species | # soft sweeps | # hard sweeps | # soft sweeps (Prob > 0.95) | # hard sweeps (Prob > 0.95) |
|---|---|---|---|---|---|
| **mel-nig** | mel | 520 | 85 | 86 | 8 |
| | nig | 324 | 116 | 41 | 12 |
| **nig-pil** | nig | 398 | 86 | 49 | 7 |
| | pil | 291 | 105 | 38 | 12 |
| **pil-pal** | pil | 277 | 44 | 28 | 7 |
| | pal | 389 | 98 | 43 | 9 |
| **hypox-mel** | hypox | 349 | 47 | 45 | 7 |
| | mel | 510 | 115 | 74 | 10 |

**Supplementary Table S10:** Confusion matrix for the classification task for species-specific versus ancestral soft sweeps in a two-species analysis. The classification task involves four classes: (1) neutral, (2-3) soft sweep in species #1 or #2, and (4) soft sweep in the population ancestral to species #1 and #2. Table layout and description are otherwise similar to Supplementary Table S6. The average classification accuracy for this task (across all cases) is 91%, and the most common mis-classifications are: (1) ancestral soft sweeps being mis-classified as neutral regions (18.30%) and (2) neutral regions being mis-classified as ancestral soft sweeps (10.10%).

| | **Neutral** | **Soft (#1)** | **Soft (#2)** | **Ancestral** |
|---|---|---|---|---|
| **Neutral** | **88.80%** | 0.60% | 0.50% | 10.10% |
| **Soft (#1)** | 0.90% | **98.20%** | 0.00% | 0.90% |
| **Soft (#2)** | 0.80% | 0.00% | **97.00%** | 2.20% |
| **Ancestral** | 18.30% | 0.50% | 1.10% | **80.10%** |

**Supplementary Table S11:** Numbers of species-specific and ancestral soft sweeps predicted across all 19 scaffolds that contain an $F_{ST}$ peak. Four separate species-pair analyses were executed, with the classification task involving four classes: (1) neutral, (2-3) soft sweep in species #1 or #2, and (4) soft sweep in the population ancestral to species #1 and #2. A four-way linear SVM classifier was trained on simulated data and then applied to the genomic data using a sliding window approach that classified each 10-kb window into one of the four classes (see **Methods**). The table reports the number of windows classified into each of the three non-neutral classes with normalized probability 0.95 or higher (FDR-adjusted *p*-value < 0.05; see **Section 4 in Supplementary Text**). Results suggest widespread species-specific soft sweeps across these 19 scaffolds.

| **Pairwise analysis** | **Species** | **# soft sweeps (Prob > 0.95)** | **# ancestral sweeps (Prob > 0.95)** |
|---|---|---|---|
| **mel-nig** | mel | 228 | 20 |
| | nig | 113 | |
| **nig-pil** | nig | 166 | 26 |
| | pil | 104 | |
| **pil-pal** | pil | 98 | 16 |
| | pal | 136 | |
| **hypox-mel** | hypox | 111 | 28 |
| | mel | 172 | |

**Supplementary Table S12:** Confusion matrix for the classification task for complete versus partial soft sweeps in a two-species analysis. The classification task involves five classes: (1) neutral, (2-3) complete soft sweep in species #1 or #2, and (4-5) partial soft sweep in species #1 or #2. Table layout and description are otherwise similar to Supplementary Table S6. The average classification accuracy for this task (across all cases) is 75.2%, and the most common mis-classifications are: (1) partial soft sweeps in both species being mis-classified as neutral regions (30.15%), (2) partial soft sweeps being mis-classified as complete soft sweeps in the same species (10.5%), (3) complete soft sweeps being mis-classified as partial soft sweeps in the same species (9.15%), and (4) neutral regions being mis-classified as partial soft sweeps in one of the two species (8.65%).

| | **Neutral** | **Complete(#1)** | **Complete(#2)** | **Partial (#1)** | **Partial (#2)** |
|---|---|---|---|---|---|
| **Neutral** | **82.60%** | 0.00% | 0.10% | 8.30% | 9.00% |
| **Complete(#1)** | 0.50% | **91.00%** | 0.00% | 8.40% | 0.10% |
| **Complete(#2)** | 0.60% | 0.20% | **89.20%** | 0.10% | 9.90% |
| **Partial (#1)** | 29.10% | 11.10% | 0.00% | **56.40%** | 3.40% |
| **Partial (#2)** | 31.20% | 0.20% | 9.90% | 2.00% | **56.70%** |

**Supplementary Table S13:** Numbers of complete and partial soft sweeps predicted across all 19 scaffolds that contain an $F_{ST}$ peak. Four separate species-pair analyses were executed, with the classification task involving five classes: (1) neutral, (2-3) complete soft sweep in species #1 or #2, and (4-5) partial soft sweep in species #1 or #2. A five-way linear SVM classifier was trained on simulated data and then applied to the genomic data using a sliding window approach that classified each 10 kb window into one of the five classes (see **Methods**). The table reports the number of windows classified into each of the four non-neutral classes with normalized probability 0.95 or higher (FDR-adjusted *p*-value < 0.05; see **Section 4 in Supplementary Text**). Results suggest slightly more prevalent partial soft sweeps, compared to complete soft sweeps in these 19 scaffolds.

| **Pairwise analysis** | **Species** | **# complete soft sweeps (Prob > 0.95)** | **# partial soft sweeps (Prob > 0.95)** |
|---|---|---|---|
| **mel-nig** | mel | 26 | 35 |
| | nig | 16 | 18 |
| **nig-pil** | nig | 17 | 22 |
| | pil | 10 | 18 |
| **pil-pal** | pil | 15 | 15 |
| | pal | 12 | 20 |
| **hypox-mel** | hypox | 19 | 11 |
| | mel | 10 | 27 |

**Supplementary Table S14:** Confusion matrix for the classification task for neutral regions and regions experiencing species-specific complete soft sweeps in the expanded five-species analysis. The classification task involves six classes: (1) neutral and (2-6) species-specific complete soft sweep in species #1 to #5. Table layout and description are otherwise similar to Supplementary Table S6. The average classification accuracy for this task (across all cases) is 94.3%, and the most common mis-classifications are soft sweeps being mis-classified as neutral regions (4.48%).

| | **Neutral** | **Soft (#1)** | **Soft (#2)** | **Soft (#3)** | **Soft (#4)** | **Soft (#5)** |
|---|---|---|---|---|---|---|
| **Neutral** | **93.25%** | 1.75% | 1.75% | 1.38% | 0.25% | 1.63% |
| **Soft (#1)** | 4.50% | **94.38%** | 0.13% | 0.25% | 0.63% | 0.13% |
| **Soft (#2)** | 4.50% | 0.50% | **94.38%** | 0.25% | 0.13% | 0.25% |
| **Soft (#3)** | 4.50% | 0.13% | 0.25% | **94.75%** | 0.25% | 0.13% |
| **Soft (#4)** | 4.50% | 0.00% | 0.50% | 0.00% | **94.75%** | 0.25% |
| **Soft (#5)** | 4.38% | 0.38% | 0.25% | 0.38% | 0.13% | **94.50%** |

**Supplementary Table S15:** Numbers of soft sweeps predicted across all 19 scaffolds that contain an $F_{ST}$ peak in the expanded five species analysis. The classification task involved six classes: (1) neutral, and (2-6) species-specific complete soft sweep in each of the five species. A six-way linear SVM classifier was trained on simulated data and then applied to the genomic data using a sliding window approach that classified each 10 kb window into one of the six classes (see **Methods**). The table reports the number of windows classified into each of the five non-neutral classes (2$^{nd}$ column). The numbers of predictions associated with normalized probability 0.95 or higher are given in the 3$^{rd}$ column (FDR-adjusted *p*-value < 0.05; see **Section 4 in Supplementary Text**).

| **Species** | **# soft sweeps** | **# soft sweeps (Prob > 0.95)** |
|---|---|---|
| **hypox** | 652 | 7 |
| **mel** | 1087 | 5 |
| **nig** | 731 | 10 |
| **pil** | 1051 | 13 |
| **pal** | 691 | 4 |

**Supplementary Table S16:** Numbers of species-specific and parallel soft sweeps predicted across all 19 scaffolds that contain an $F_{ST}$ peak. Four separate species-pair analyses were executed, with the classification task involving four classes: (1) neutral, (2-3) species-specific soft sweep in species #1 or #2, and (4) parallel soft sweeps occurring separately and independently in both species (#1 and #2). A four-way linear SVM classifier was trained on simulated data and then applied to the genomic data using a sliding window approach that classified each 10 kb window into one of the four classes (see **Methods**). The table reports the number of windows classified into each of the three non-neutral classes (with "# soft sweeps" indicating the total number of species-specific soft sweeps predicted for the two species).

| **Pairwise analysis** | **# soft sweeps** | **# parallel soft sweeps** |
|---|---|---|
| **mel-nig** | 99 | 149 |
| **nig-pil** | 80 | 119 |
| **pil-pal** | 63 | 124 |
| **hypox-mel** | 77 | 177 |

#### Supplementary Figures

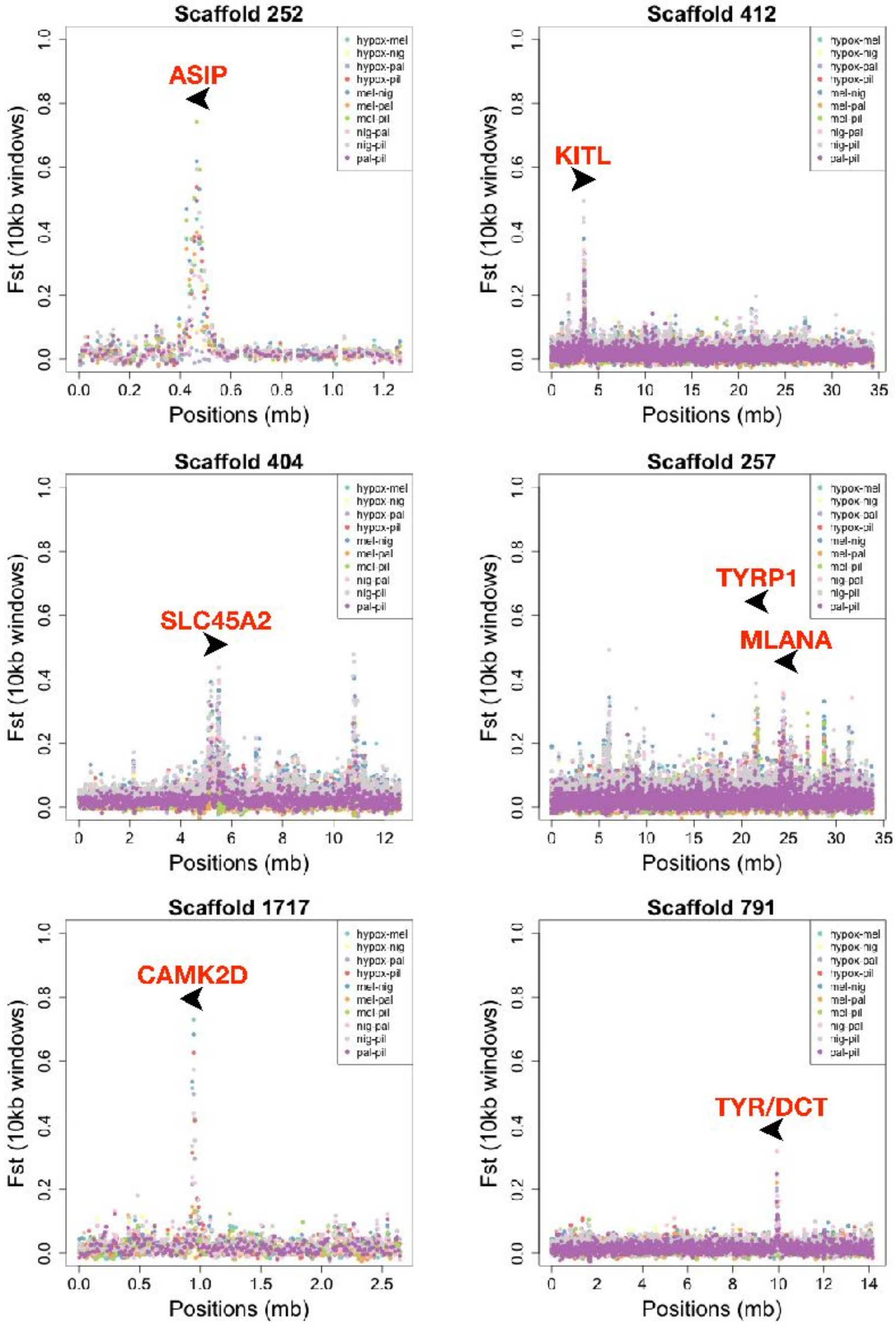


**Supplementary Figure S1:** Distribution of average $F_{ST}$ computed across six scaffolds reported by Campagna *et al.* (2017)[1] to contain the top $F_{ST}$ peaks near pigmentation-related genes. A complete summary of all peaks can be found in Campagna *et al.* (2017)[1]. Values are computed for all 10 pairwise comparisons (see legend), and averaged in non-overlapping 10 kb windows. Pigmentation genes involved in the melanogenesis pathway are indicated in red.

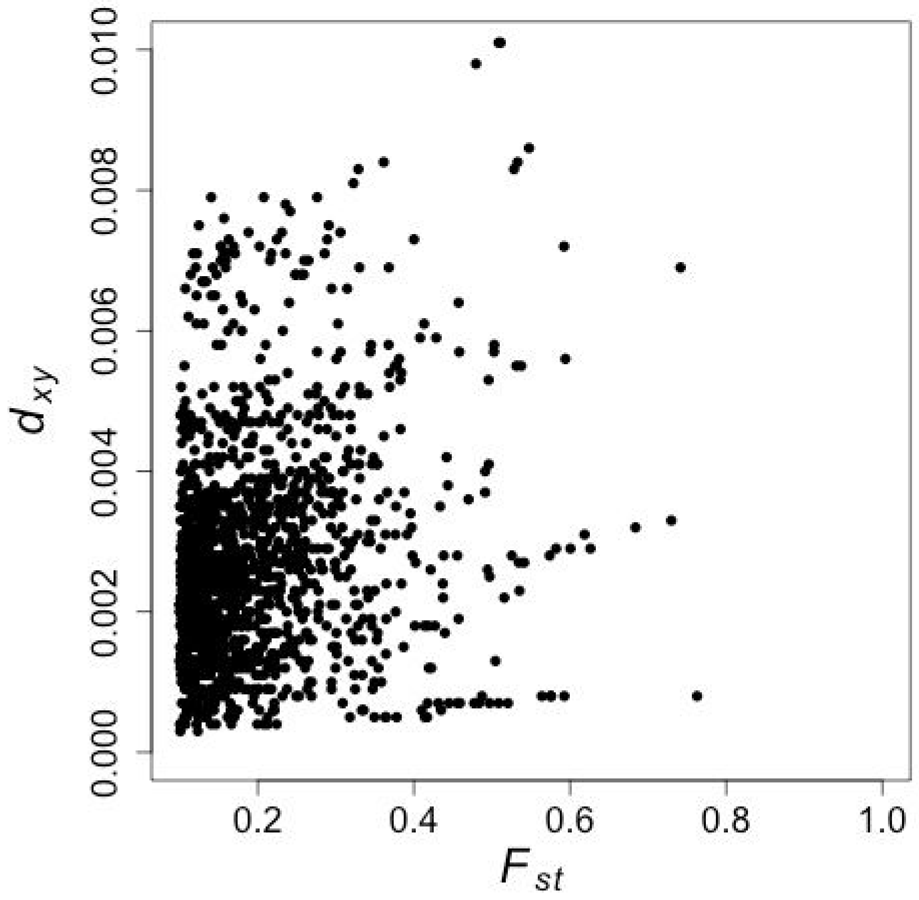


**Supplementary Figure S2:** Relationship between $F_{ST}$ and $d_{XY}$ in $F_{ST}$ peaks. Average values of $F_{ST}$ and $d_{XY}$ were computed for each of the ten species-pairs in non-overlapping 10 kb windows that intersect with one of the 25 $F_{ST}$ peaks. Every data point corresponds to a 10 kb window and a pair of species. To focus on cases of elevated relative sequence differentiation, we restrict the plot to data points with $F_{ST}$>0.1. We find a relatively low correlation between $F_{ST}$ and $d_{XY}$ overall (Pearson's $r$=0.197), contrary to what we would expect if selection against gene flow was a dominant factor in the formation of the $F_{ST}$ peaks (see Figure 1).

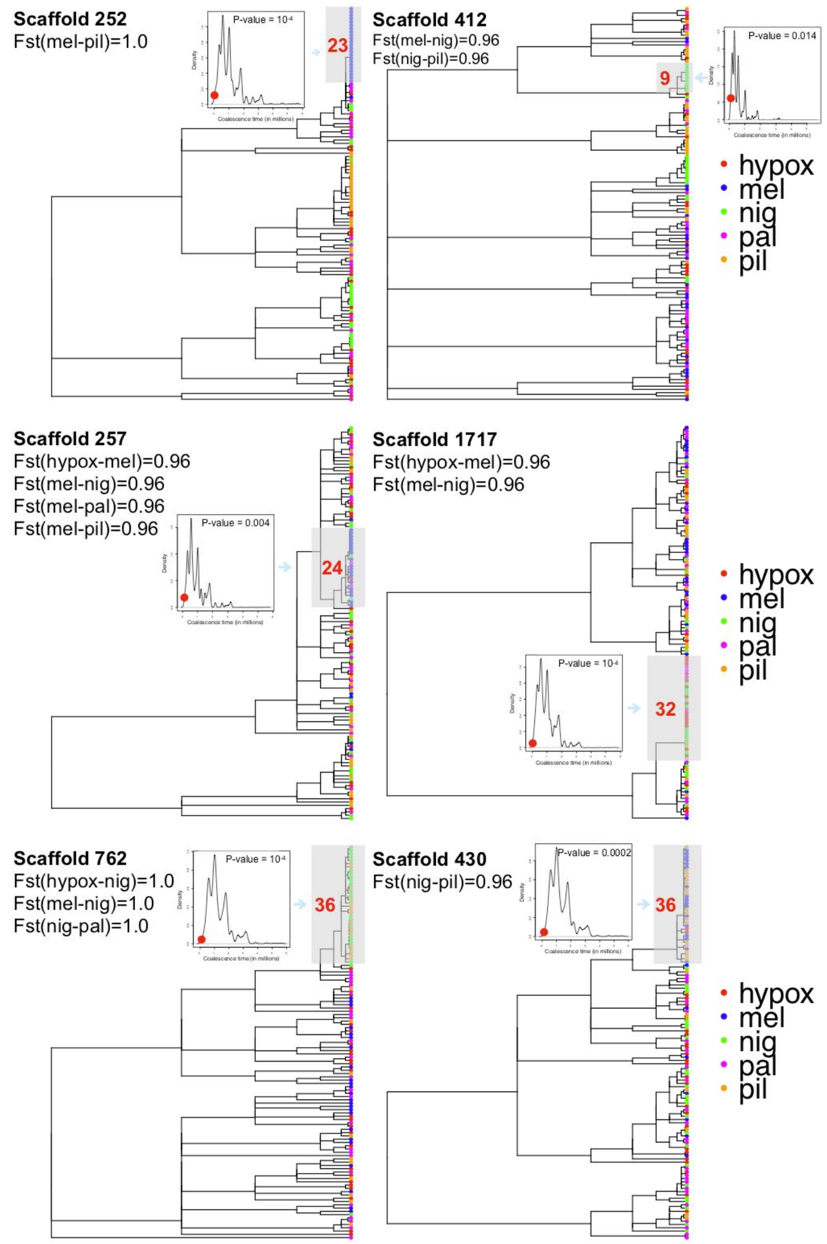


**Supplementary Figure S3:** Genealogical indications of species-specific selective sweeps in six $F_{ST}$ peaks. In each peak, we examined the local tree inferred by *ARGweaver* at the site with the highest $F_{ST}$ value in the peak. For each peak, we specify the highest $F_{ST}$ value (attained in the focal site) along with the pairwise comparison(s) it originated from. The tips of the trees are colored based on their species labels (see color legend). In each tree, we focused on a single clade that exhibited a “burst” of coalescence events (gray highlight; **Supplementary Text**). The age of this clade was compared to the age distribution of clades of identical size (indicated in red) recorded in local trees extracted from the same scaffold, but outside the $F_{ST}$ peak. The background age distribution is shown in the inset, with the red dot indicating the age of the focal clade observed in the $F_{ST}$ peak. The *p*-value resulting from this comparison is also specified in the inset. All six peaks contain focal clades that are significantly younger than clades in the flanking regions ($p$<0.015), with four of them being particularly striking ($p$<0.001).

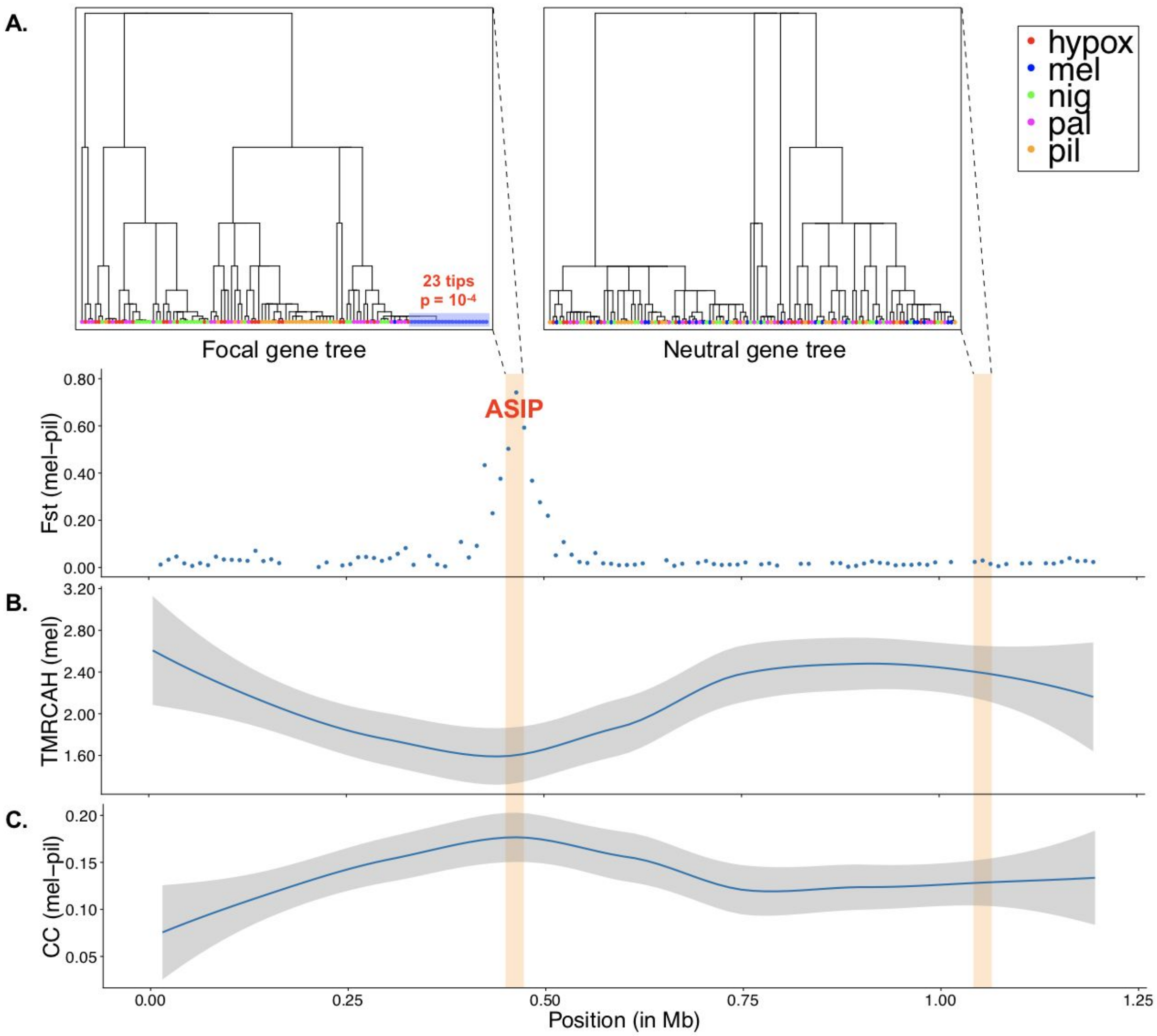


**Supplementary Figure S4:** Comparison of inferred genealogies in an $F_{ST}$ peak and flanking neutral regions of scaffold 252. (**A**) Average $F_{ST}$ values in non-overlapping 10 kb windows between *S. melanogaster* (mel) and *S. pileata* (pil), showing a pronounced peak near the pigmentation-related gene *ASIP*. Local genealogies within the peak (*top left*) and in a flanking neutral region (*top right*), representing the relationships among 120 haploid samples from five species (see *legend*). The tree within the $F_{ST}$ peak has a very young clade (*gray box*) which is enriched for *S. melanogaster* and has a more recent TMRCA than expected given its size ($p$=$10^{-4}$, **Supplementary Text**). By contrast, the flanking tree is characterized by abundant deep coalescence events and incomplete lineage sorting. (**B**) Times of most recent common ancestry (in millions of generations) for half of the haploid samples (TMRCAH) from *S. melanogaster*. Values are reduced in the $F_{ST}$ peak compared to flanking regions, suggesting species-specific selective sweeps. (**C**) Cross-coalescence times (CC) in millions of generations between the two species, as computed from the inferred ARG. Elevated cross coalescence times are observed in the $F_{ST}$ peak, indicating possible influence of selection against gene flow on this peak (see also **Supplementary Figure S5** and **Supplementary text**). The smoothing method used in panels B and C was local polynomial regression fitting along with a 95% confidence interval.

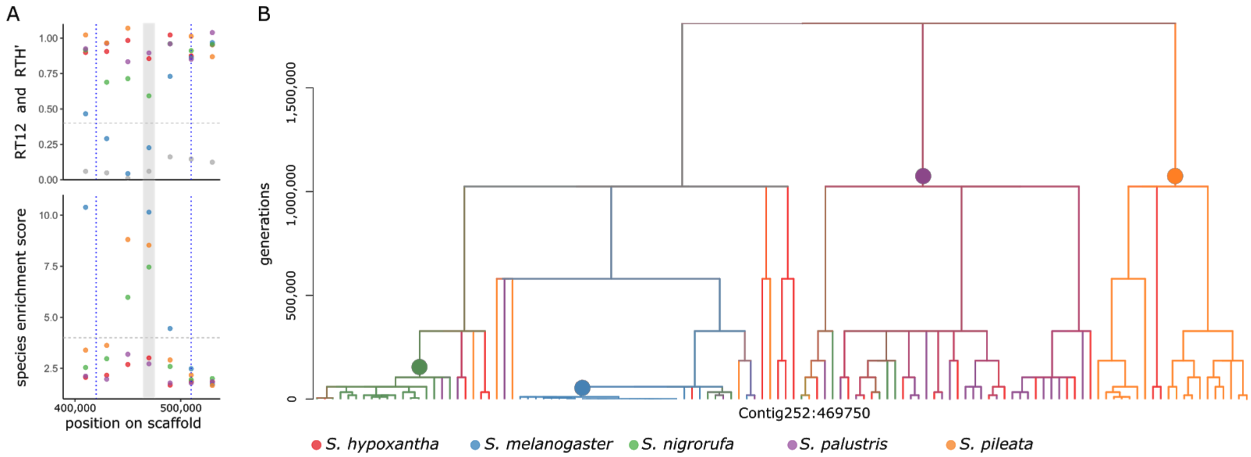


**Supplementary Figure S5:** Genealogical statistics and a representative tree for the $F_{ST}$ peak on scaffold 252. (A) RTH', RT12, and species enrichment scores averaged across non-overlapping 20 kb windows in the genomic region containing the divergence peak. The peak boundaries are indicated by vertical dashed lines. Species-specific scores are marked by color (see legend), and RT12 is marked by gray circles. Horizontal dashed lines correspond to mean significance thresholds across species (see **Supplementary Tables S2** and **S3** for species-specific thresholds). (B) A representative tree extracted from the inferred ARG inside the 20 kb window with the highest enrichment scores within the peak (gray vertical bar in panel A). The color of leaf branches corresponds to the species label of the haploid sample (see legend), and the color of internal branches corresponds to the average color across all samples in the clade below the branch. In this peak we observe significantly high enrichment scores for *S. melanogaster*, *S. nigrorufa*, *S. palustris*, and *S. pileata*, and a significantly low RTH' for *S. melanogaster* (**Table 1**, and **Supplementary Tables S2** and **S3**). The four corresponding species-enriched clades are indicated by colored circles at their roots. The young clade enriched for *S. melanogaster* suggests a very recent species-specific selective sweep for this species in this region. On the other hand, the deep clade enriched for *S. pileata* suggests selection against gene flow between this species and the other four. This observation is also supported by elevated cross coalescence times (**Supplementary Table S5**, see **Supplementary Text** for more details).

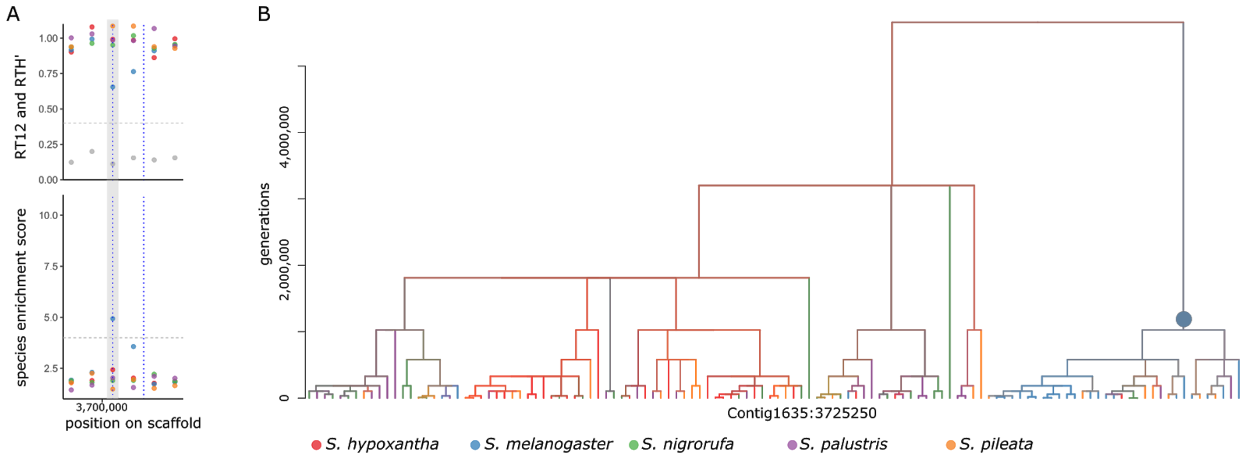


**Supplementary Figure S6:** Genealogical statistics and a representative tree for the $F_{ST}$ peak on scaffold 1635. (A) RTH', RT12, and species enrichment scores averaged across non-overlapping 20 kb windows in the genomic region containing the divergence peak. (B) A representative tree extracted from the inferred ARG inside the 20 kb window with the highest enrichment scores within the peak (gray vertical bar in panel A). See caption of **Supplementary Figure S5** for complete specification of color code and vertical and horizontal dashed lines in panel A. In this peak we observe a significantly high enrichment score for *S. melanogaster*, and no significantly reduced RTH' (**Table 1**, and **Supplementary Tables S2** and **S3**). A clade enriched for *S. melanogaster* is indicated by a colored circle at its root. This clade is roughly one million generations old and contains 33 samples, out of which 21 are from *S. melanogaster*. This deep enriched clade suggests selection against gene flow between *S. melanogaster* and the other four species, possibly excluding *S. pileata*, which has six samples within this clade. This observation is also supported by elevated cross coalescence times between *S. melanogaster* and *S. nigrorufa* (**Supplementary Tables S5**).

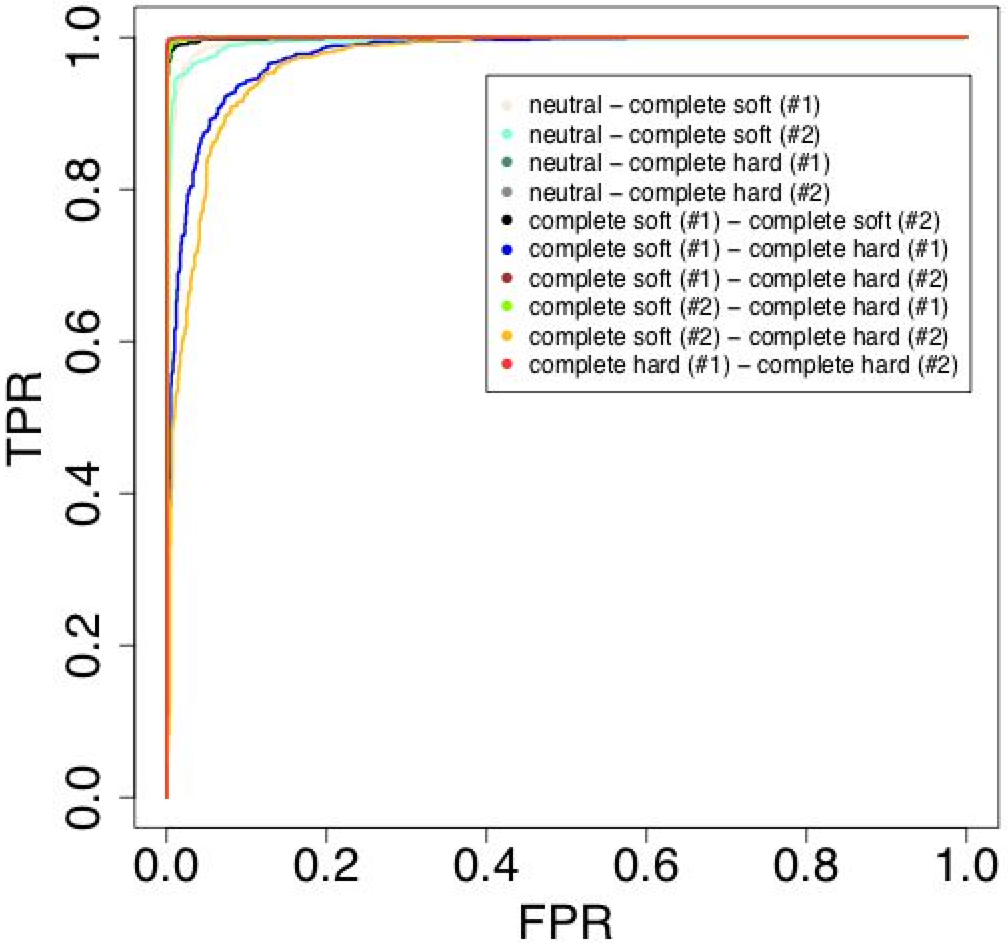


**Supplementary Figure S7:** Receiver Operating Characteristic (ROC) curves for the classification task for soft versus hard sweeps in a two-species analysis. The classification task involves five classes: (1) neutral, (2-3) soft sweep in species #1 or #2, and (4-5) hard sweep in species #1 or #2. A five-way linear SVM classifier was trained using a training set of 40,000 simulated regions of length 50 kb (8,000 per class; see **Methods**). The classifier was then tested on a separate set comprised of 5,000 regions (1,000 per class). Here, we consider all pairs of classes (see legend), and the ROC curve for each pair records the true positive rate (TPR) as a function of the false positive rate (FPR). The curve associated with the pair "class A – class B" is obtained by varying the prediction threshold from 0 to 1 and recording for each threshold the number of test regions from class A and class B that the classifier assigns to class B (with prediction probability above the threshold). The Area Under Receiver Operating Characteristics (AUROC) for these curves ranged between 0.96 and 0.99.

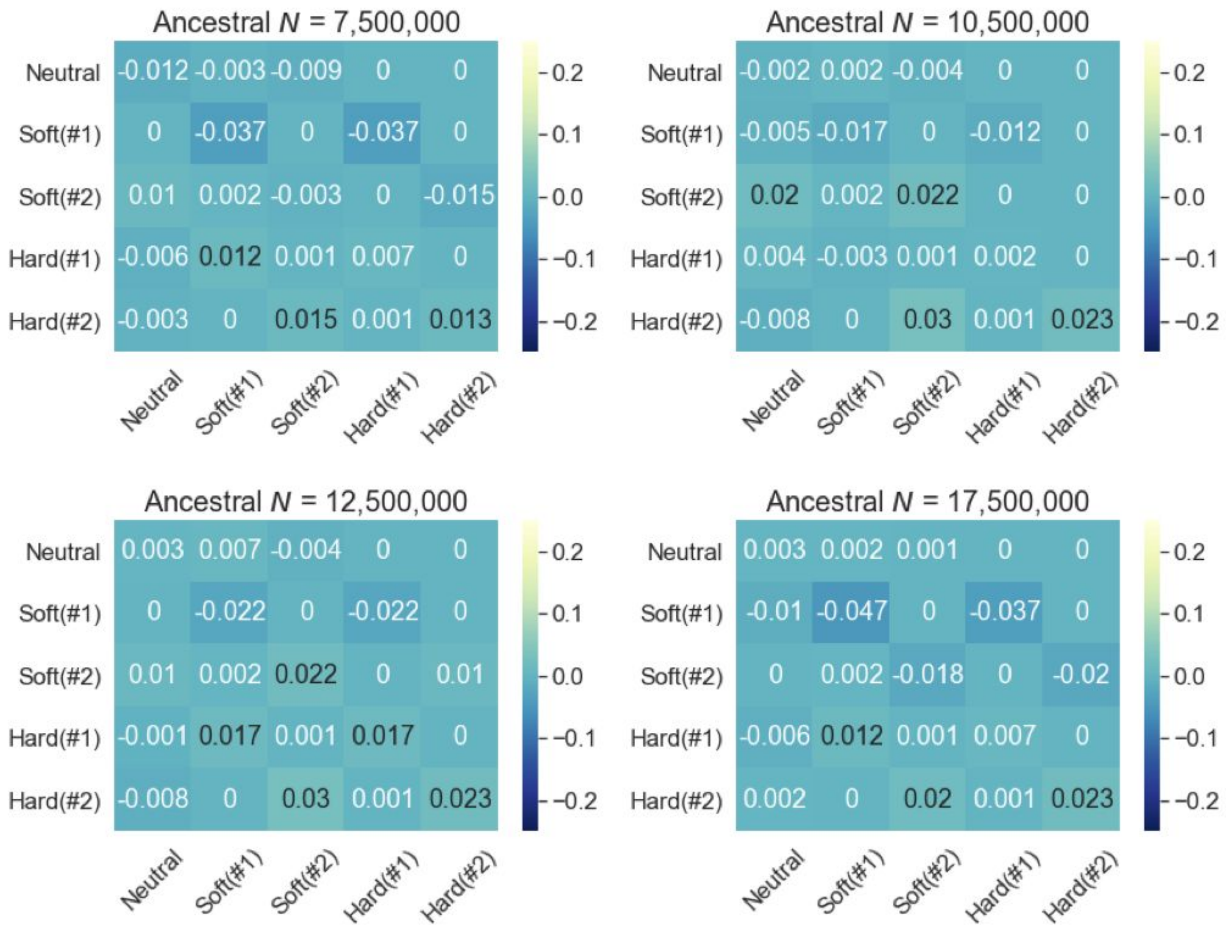


**Supplementary Figure S8:** Relative performance of the classifier on a simulated data set in which the model is misspecified in comparison to one in which the model is correctly specified. The evaluation is done on the classification task for soft versus hard sweeps in a two-species scenario. The classification task involves five classes: (1) neutral, (2-3) soft sweep in species #1 or #2, and (4-5) hard sweep in species #1 or #2. A five-way linear SVM classifier was trained using a training set of 40,000 simulated regions of length 50 kb (8,000 per class; see **Methods**). The classifier was then applied separately to a correctly-specified dataset and a misspecified dataset (as detailed in the Supplemental Text). A standard confusion matrix was generated for each data set, such that the cell in row *i* and column *j* reports the fraction of regions simulated for class *i* that the SVM assigned to class *j*. The values shown above represent the differences between these two confusion matrices. A negative value indicates a decrease in performance owing to model misspecification and a positive value indicates an increase in performance. Notice that the values on the diagonal represent differences in classification accuracy, and off-diagonal entries represent differences in various mis-classification rates.

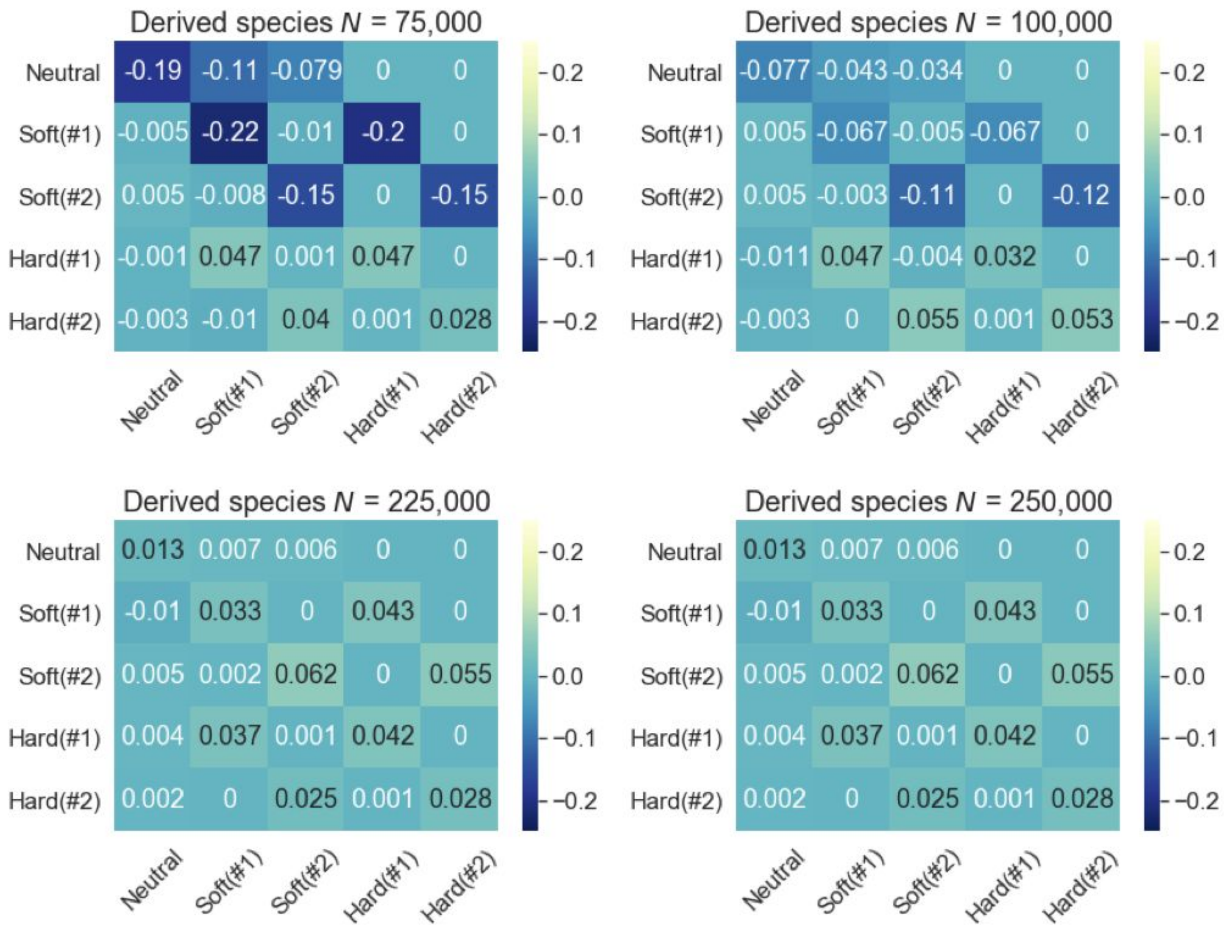


**Supplementary Figure S9:** Relative performance of the classifier on a simulated data set in which the derived species population size is misspecified in comparison to one in which the model is correctly specified. Figure layout and description are similar to Supplementary Figure S8.

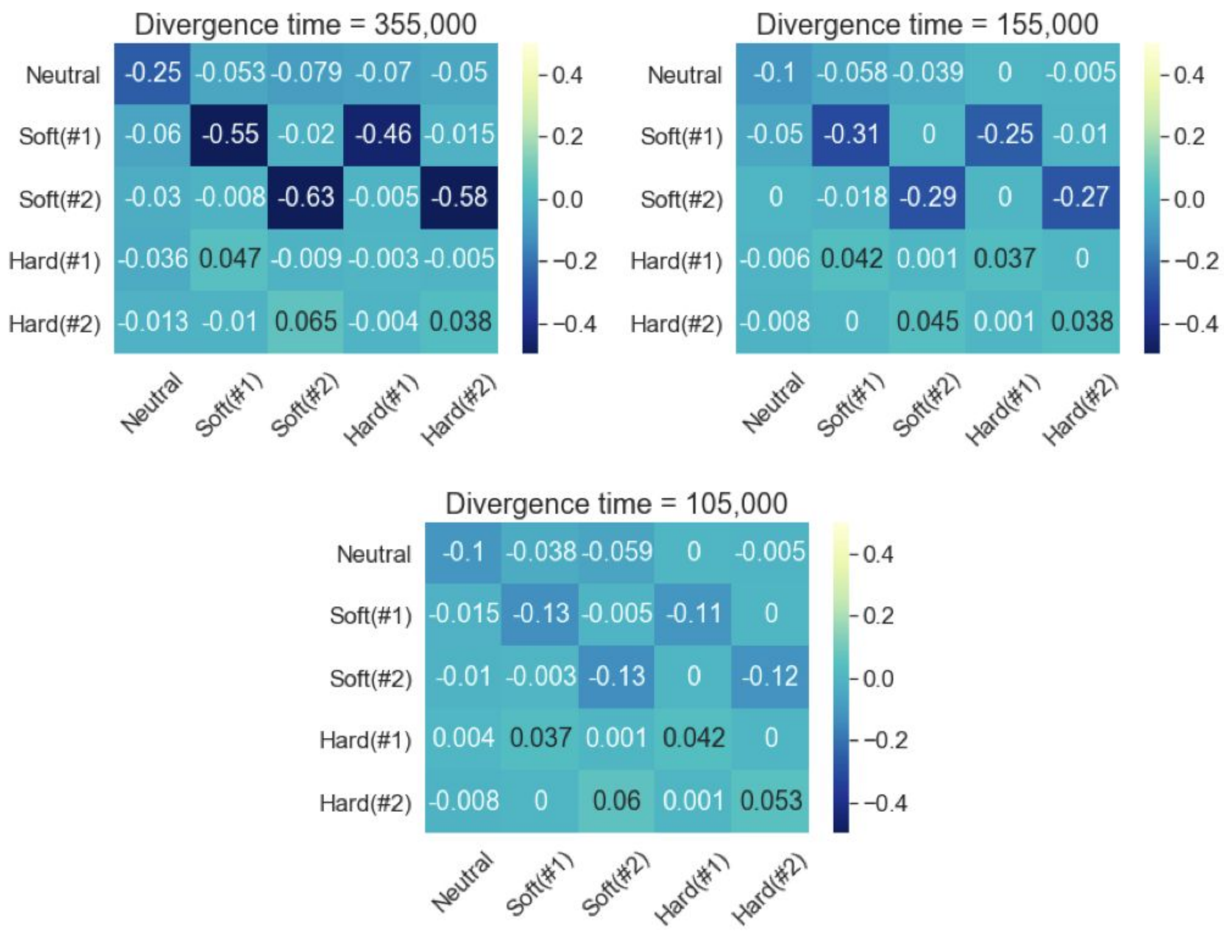


**Supplementary Figure S10:** Relative performance of the classifier on a simulated data set in which the divergence time is misspecified in comparison to one in which the model is correctly specified. Figure layout and description are similar to Supplementary Figure S8.

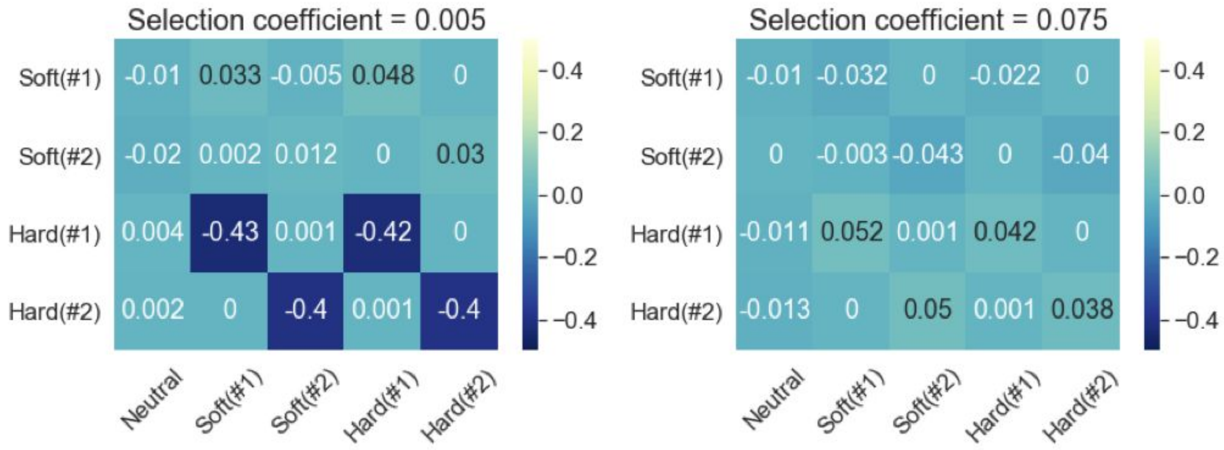


**Supplementary Figure S11:** Relative performance of the classifier on a simulated data set in which the selection coefficient is misspecified in comparison to one in which the model is correctly specified. Figure layout and description are similar to Supplementary Figure S8.

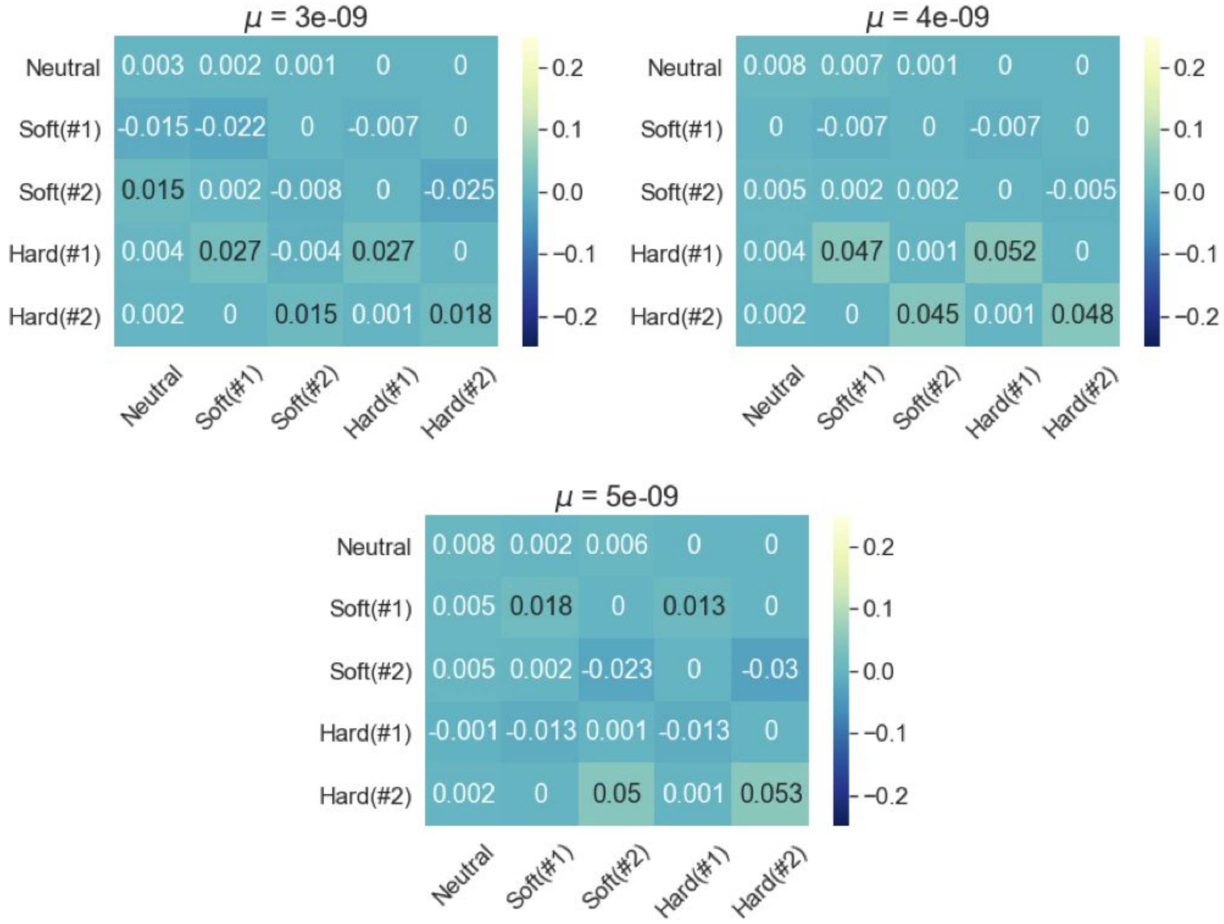


**Supplementary Figure S12:** Relative performance of the classifier on a simulated data set in which the mutation rate is misspecified in comparison to one in which the model is correctly specified. Figure layout and description are similar to Supplementary Figure S8.

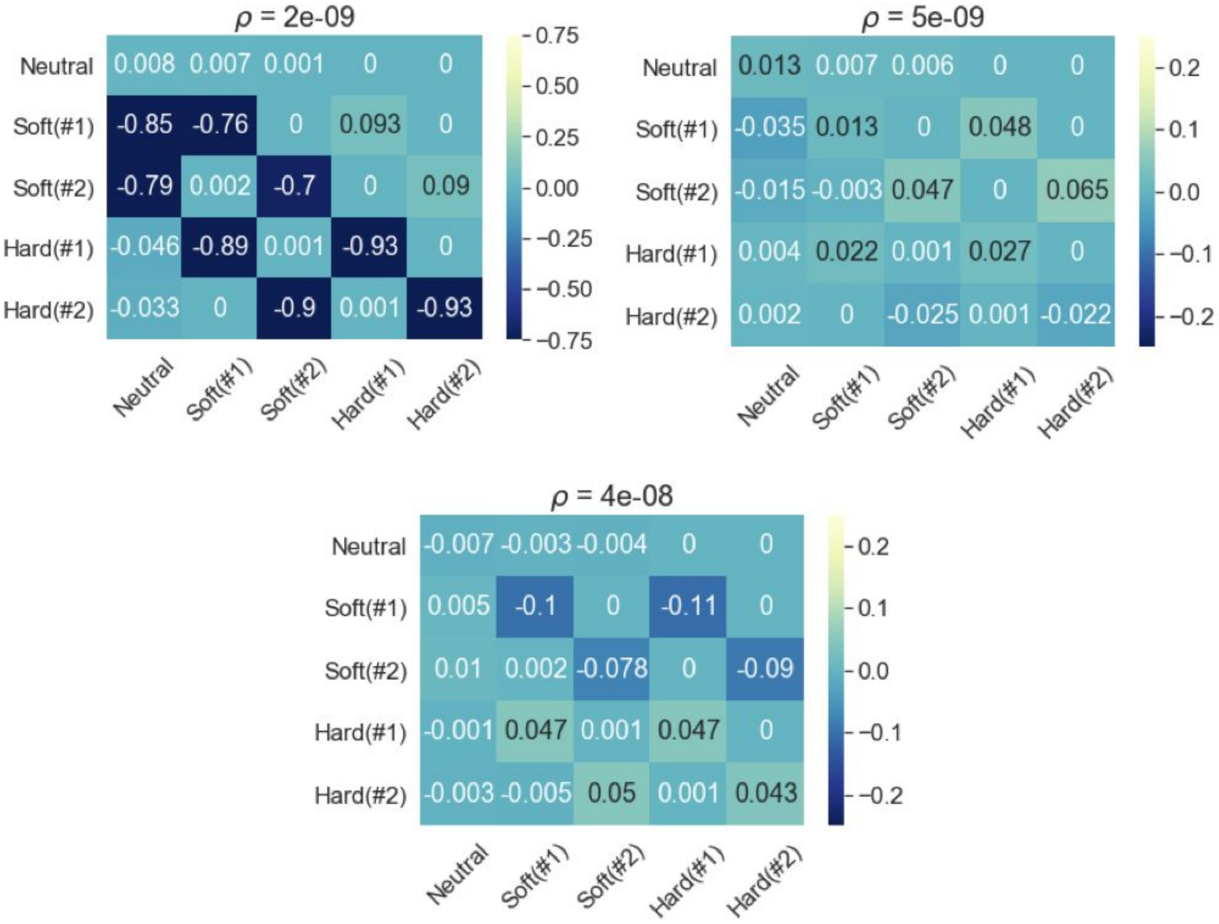


**Supplementary Figure S13:** Relative performance of the classifier on a simulated data set in which the recombination rate is misspecified in comparison to one in which the model is correctly specified. Figure layout and description are similar to Supplementary Figure S8.

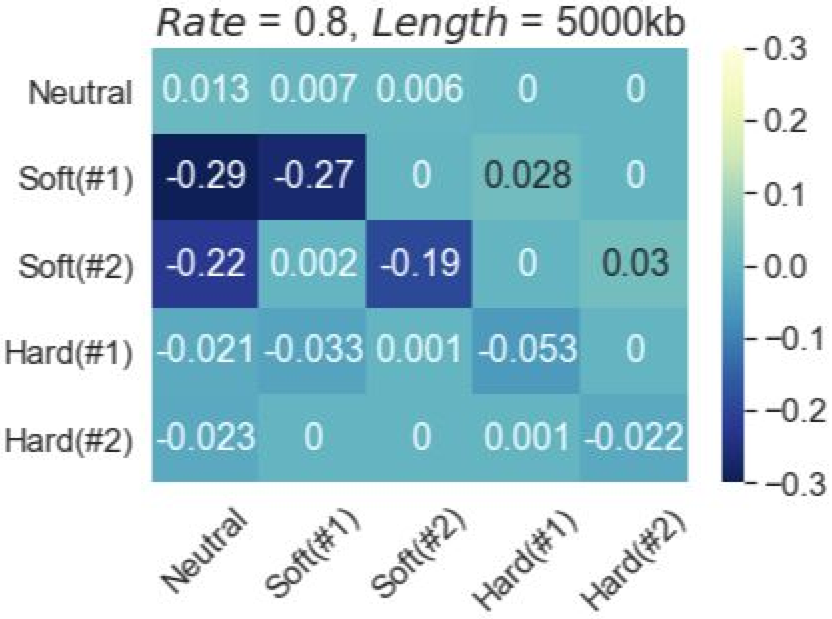


**Supplementary Figure S14:** Relative performance of the classifier on a simulated data set in which gene conversion is included in comparison to one in which the model is correctly specified not to include gene conversion (see details in **Supplemental Text**). Figure layout and description are similar to Supplementary Figure S8.

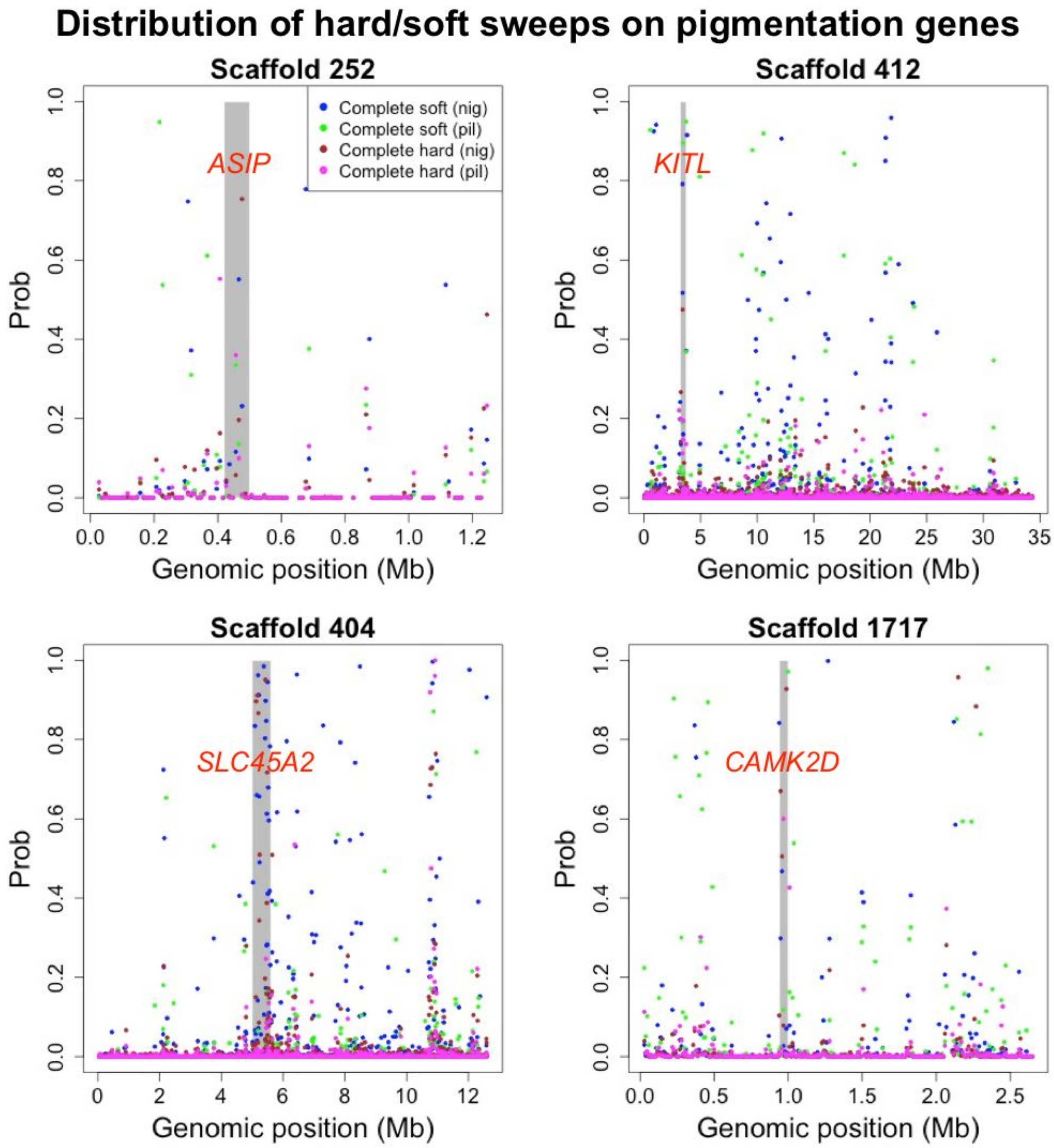


**Supplementary Figure S15:** Manhattan plots showing predicted soft and hard sweeps for *S. nigrorufa* (nig) and *S. pileata* (pil) across four scaffolds harboring top $F_{ST}$ peaks and known pigmentation genes (labeled in *red*). Each dot indicates the normalized probability assigned by the five-way classifier in a 10 kb window to one of the four non-neutral classes (see legend).

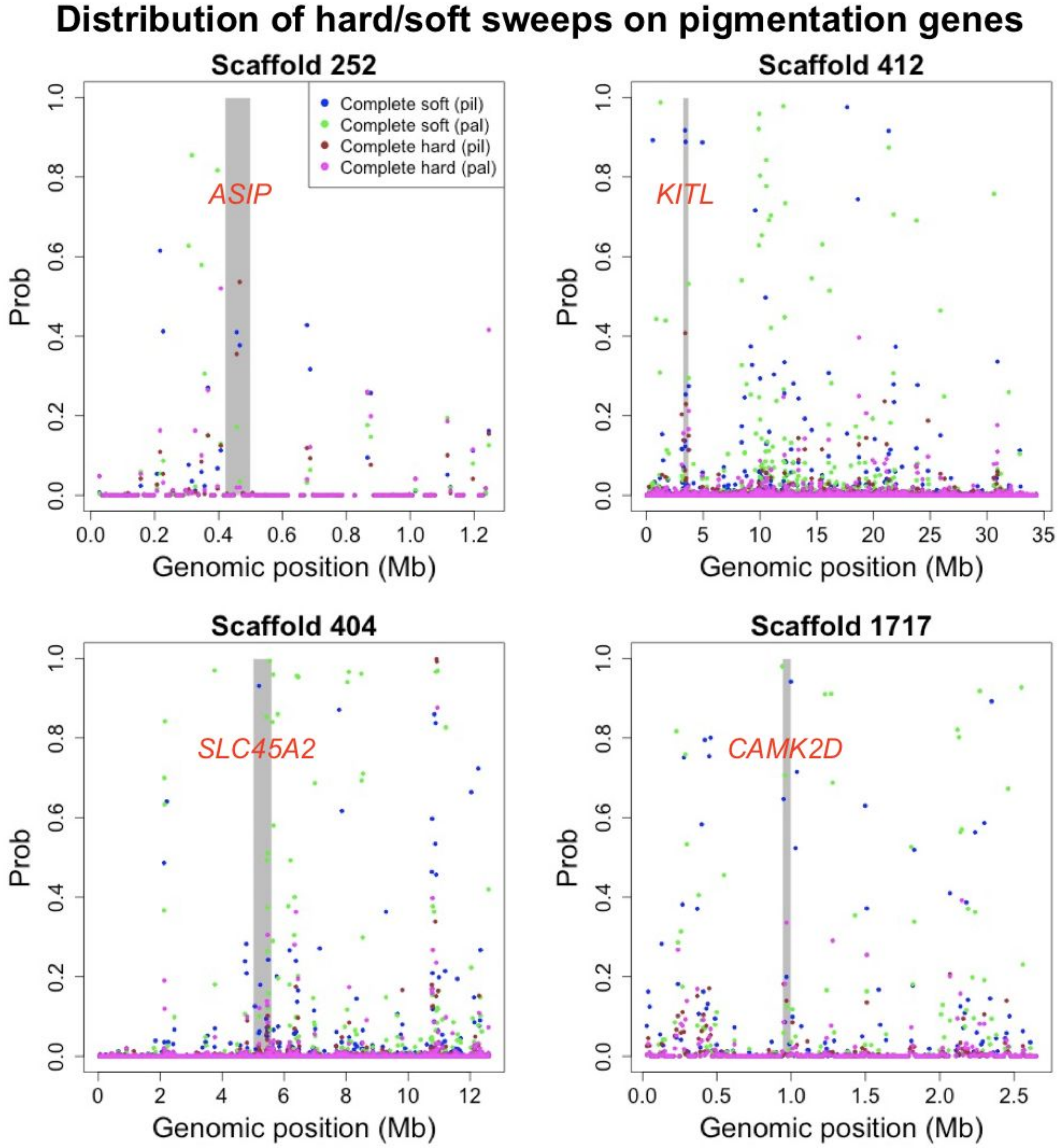


**Supplementary Figure S16:** Manhattan plots showing predicted soft and hard sweeps for *S. pileata* (pil) and *S. palustris* (pal) across four scaffolds harboring top $F_{ST}$ peaks and known pigmentation genes (labeled in *red*). Each dot indicates the normalized probability assigned by the five-way classifier in a 10 kb window to one of the four non-neutral classes (see legend).

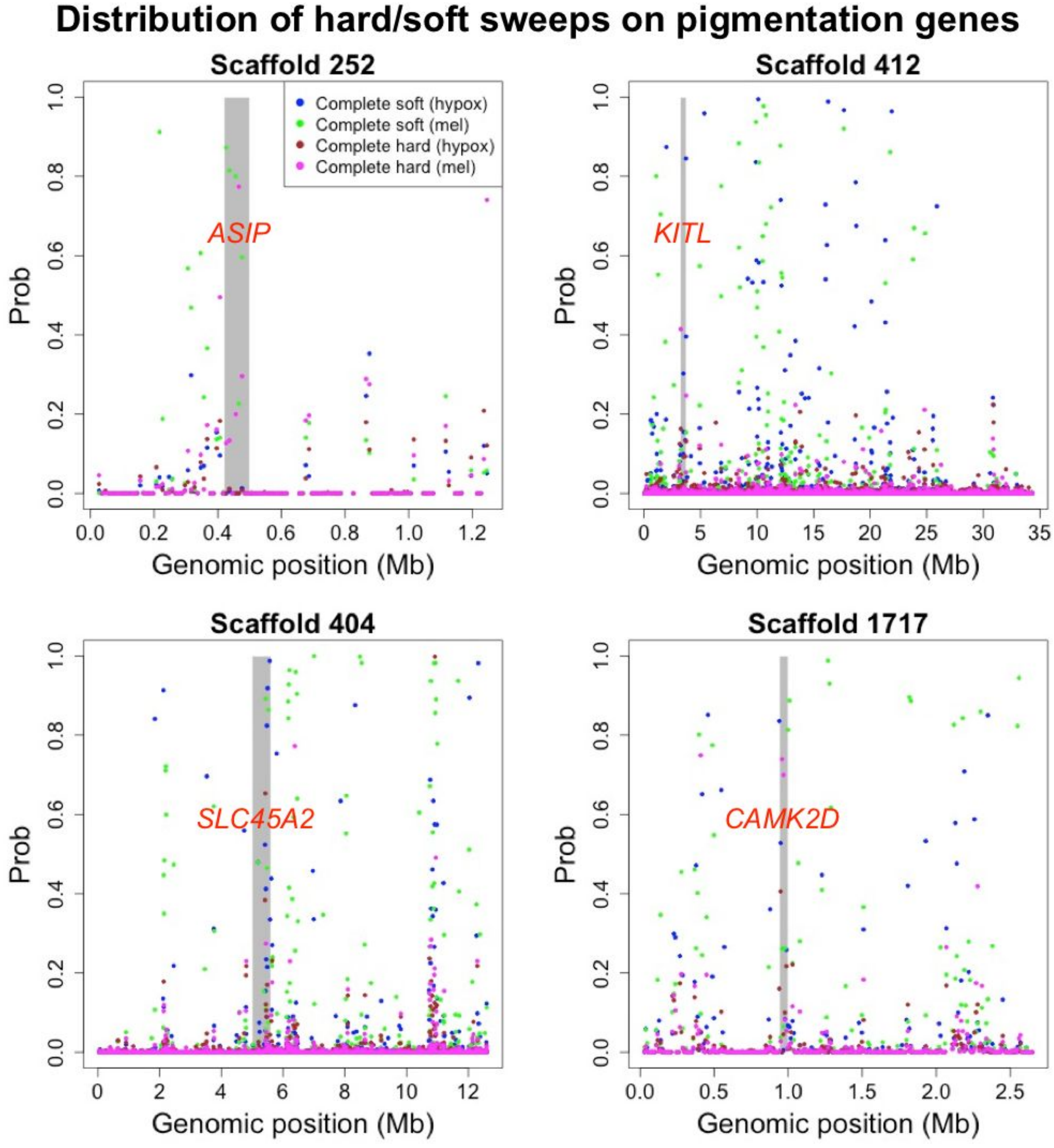


**Supplementary Figure S17:** Manhattan plots showing predicted soft and hard sweeps for *S. hypoxantha* (hypox) and *S. melanogaster* (mel) across four scaffolds harboring top $F_{ST}$ peaks and known pigmentation genes (labeled in *red*). Each dot indicates the normalized probability assigned by the five-way classifier in a 10 kb window to one of the four non-neutral classes (see legend).

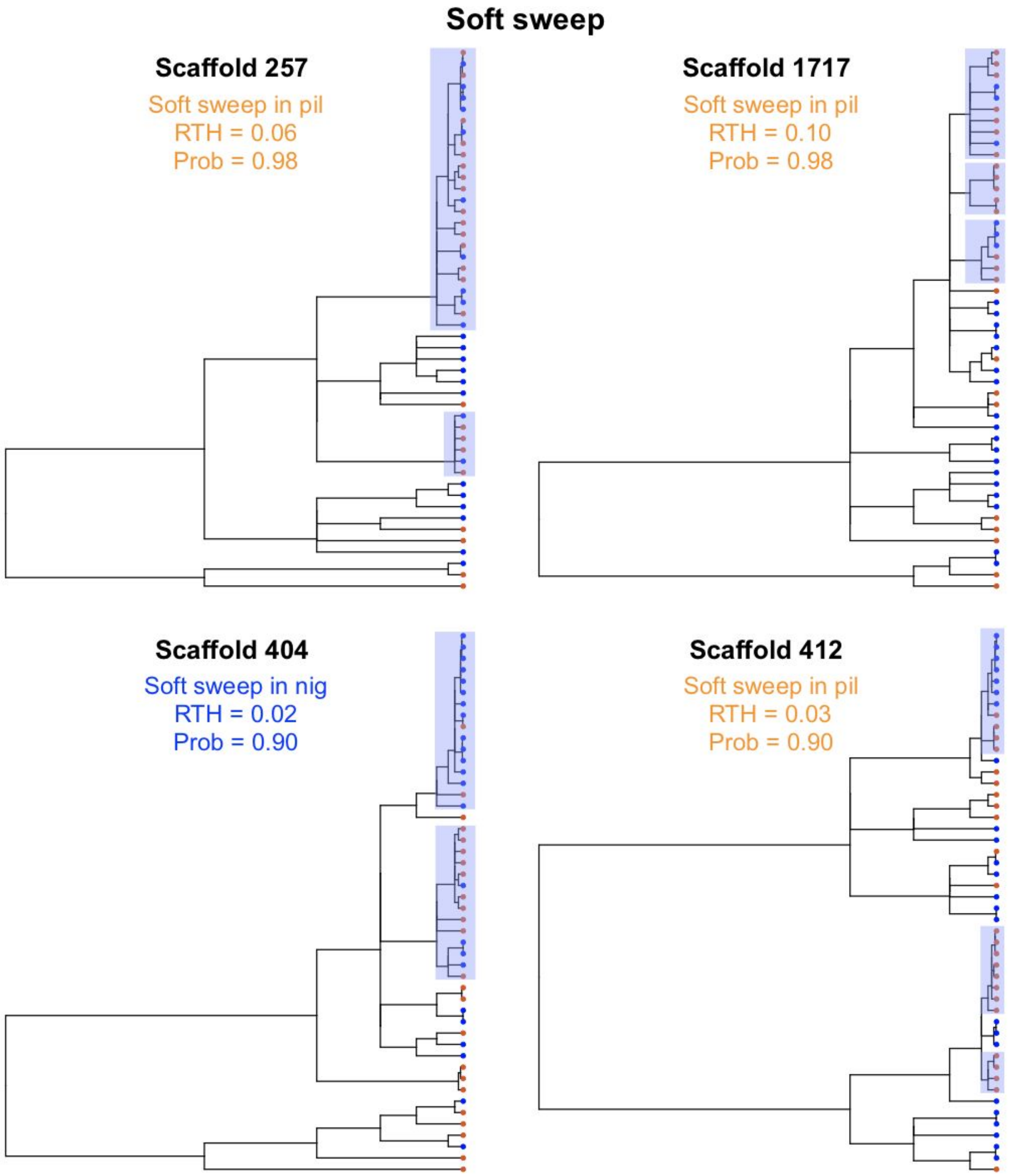


**Supplementary Figure S18:** Example of local trees extracted from 10 kb windows classified as soft sweeps in the species-pair analysis of *S. nigrorufa* (nig) and *S. pileata* (*pil*). In each window, we extracted the local tree inferred by *ARGweaver*, which had the lowest RTH for the species that was predicted to experience the sweep. The tree leaves were pruned to show only the haploid samples from *S. nigrorufa* (blue) and *S. pileata* (orange). For each tree, we report the prediction probability of the assigned class and the RTH of the species for which the soft sweep was inferred (font color indicates the species). Each of these four trees contains multiple young and moderately large clades enriched for the species associated with the sweep, suggesting that multiple haplotypes rose to high frequency in these regions in parallel, as expected in the scenario of soft sweeps.

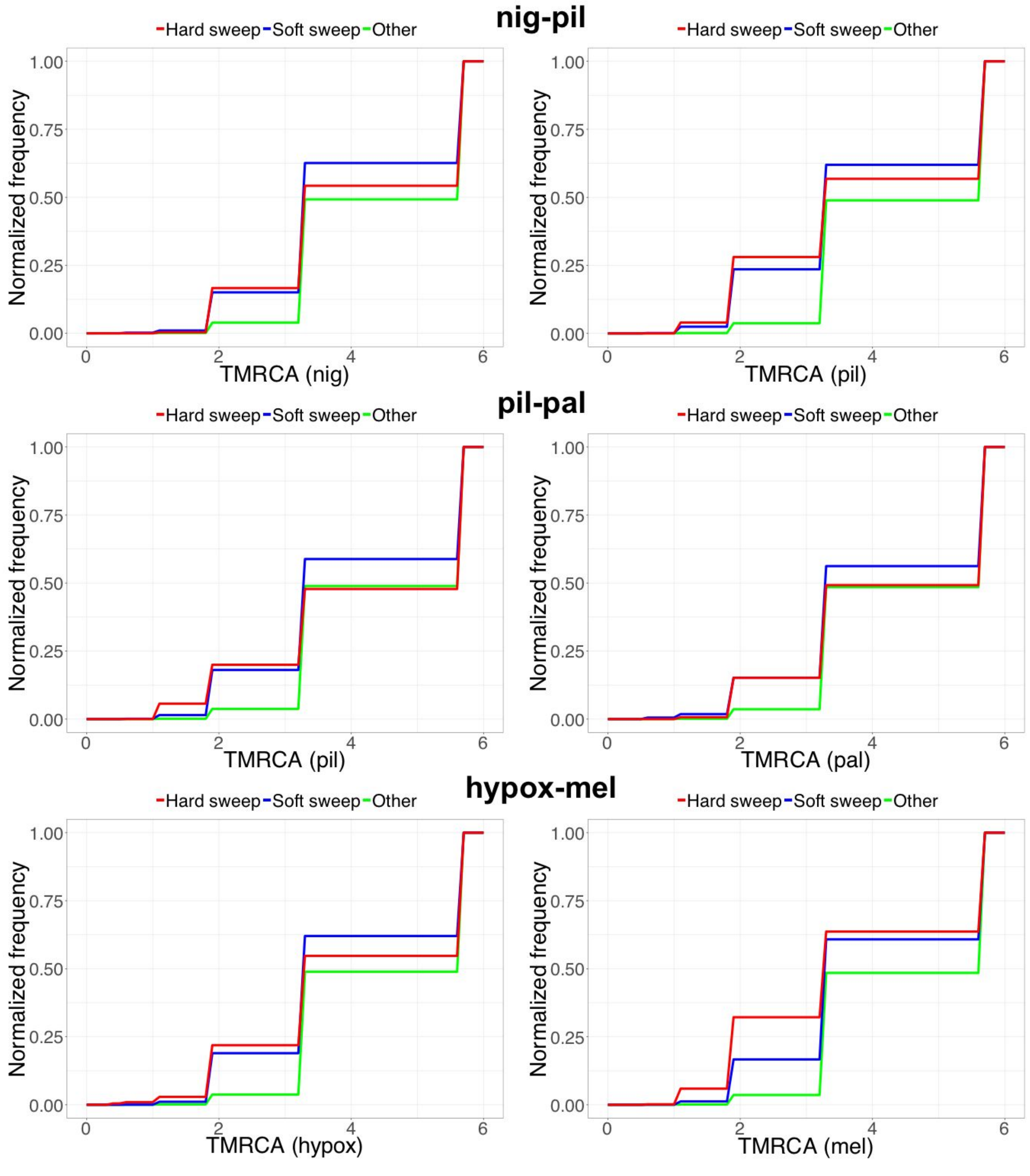


**Supplementary Figure S19:** Cumulative distribution functions (CDFs) for species-specific TMRCAs as measured in local trees inferred by *ARGweaver* in regions partitioned according to their classification in the species-pair analyses. The TMRCA of every species appears to be considerably reduced in regions predicted to be influenced by soft or hard selective sweeps in that species, as would be expected in theory. However, little differences are observed between soft and hard sweeps.

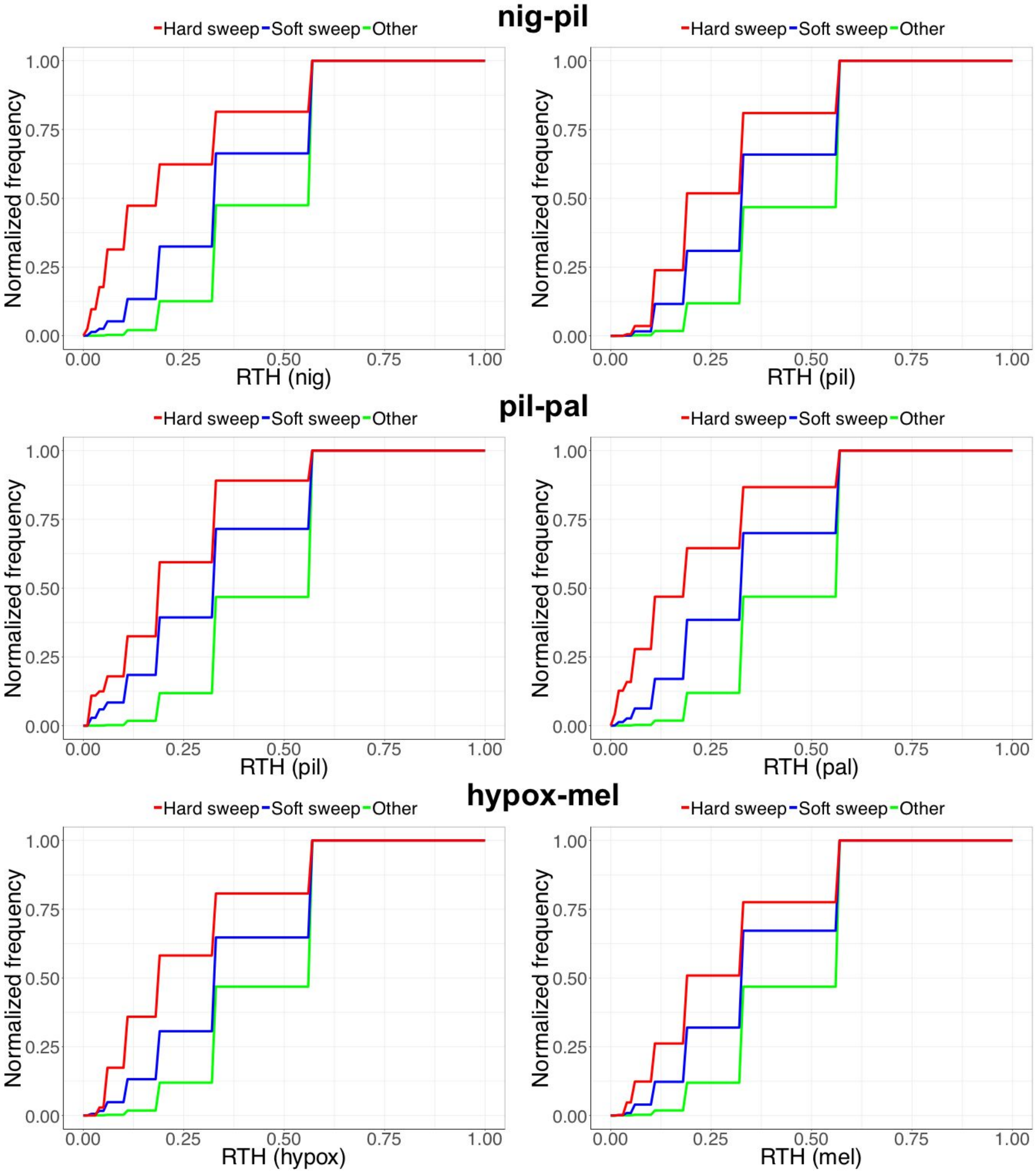


**Supplementary Figure S20:** Cumulative distribution functions (CDFs) for species-specific RTHs as measured in local trees inferred by *ARGweaver* in regions partitioned according to their classification in the species-pair analyses. The RTH is defined as the age of the youngest clade that contains at least half of the haploid samples from a given species, divided by the TMRCA of that species. The RTH of all species appears to be considerably reduced in regions predicted to be influenced by soft sweeps or hard selective sweeps in that species, as would be expected in theory. A greater reduction is observed in regions predicted as hard sweeps, likely reflecting the fact that in hard sweeps the mutation starts sweeping when it first appears in a single individual.

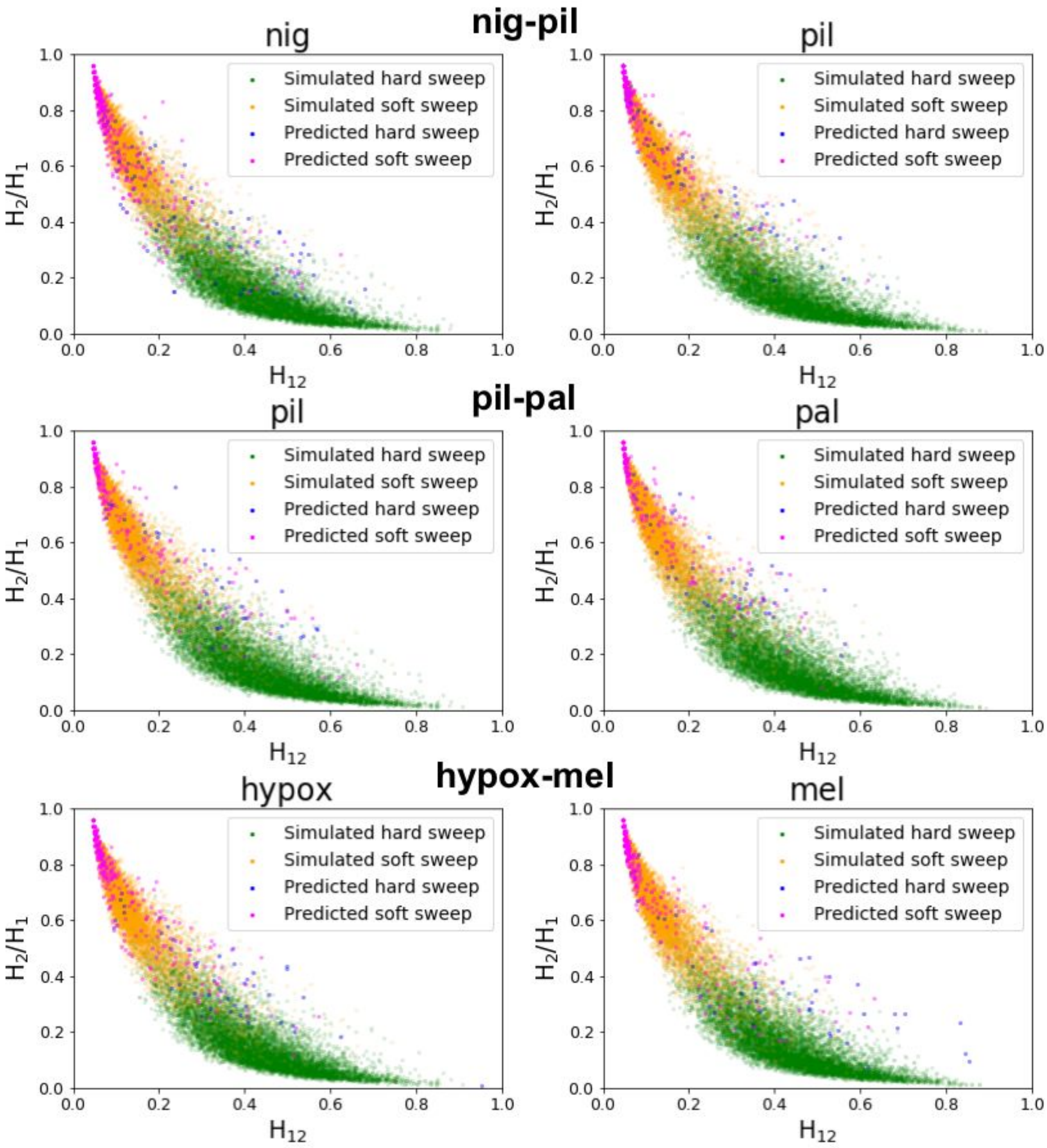


**Supplementary Figure S21:** Haplotype homozygosity statistics used to differentiate soft sweeps from hard sweeps in the species-pair analyses. Each dot depicts the $H_{12}$ and $\frac{H_2}{H_1}$ statistics for a 50 kb region, either simulated or genomic (see **Methods**). Regions simulated with hard sweeps (green) tend to have high $H_{12}$ and low $\frac{H_2}{H_1}$, whereas regions simulated with soft sweeps (orange) tend to have low $H_{12}$ and high $\frac{H_2}{H_1}$. Genomic regions predicted to contain soft sweeps in the species-pair analyses (magenta) also tend to have low $H_{12}$ and high $\frac{H_2}{H_1}$. This indicates that the great majority of them are indeed more likely to contain soft sweeps than hard sweeps. Genomic regions predicted to contain hard sweeps in the species-pair analyses (blue) are much fewer, but most of them appear to have high $H_{12}$ and low $\frac{H_2}{H_1}$, as their simulated counterparts.

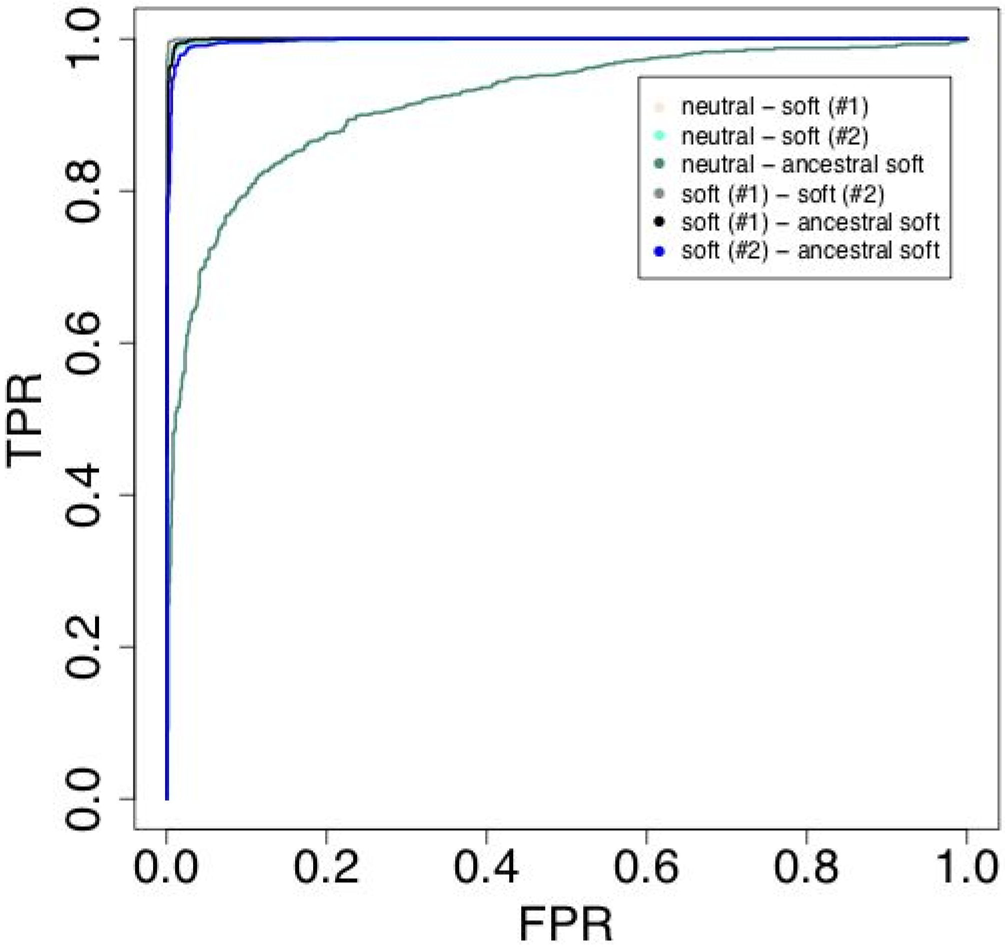


**Supplementary Figure S22:** Receiver Operating Characteristic (ROC) curves for the classification task for species-specific versus ancestral soft sweeps in a two-species analysis. The classification task involves four classes: (1) neutral, (2-3) soft sweep in species #1 or #2, and (4) soft sweep in the population ancestral to species #1 and #2. A four-way linear SVM classifier was trained using a training set of 32,000 simulated regions of length 50 kb (8,000 per class; see **Methods**). The classifier was then tested on a separate set comprised of 4,000 regions (1,000 per class). Here, we consider all pairs of classes (see legend), and the ROC curve for each pair records the true positive rate (TPR) as a function of the false positive rate (FPR). See caption of Supplementary Figure S7 for details on the experimental setup.The Area Under Receiver Operating Characteristics (AUROC) for these curves ranged between 0.91 and 0.99.

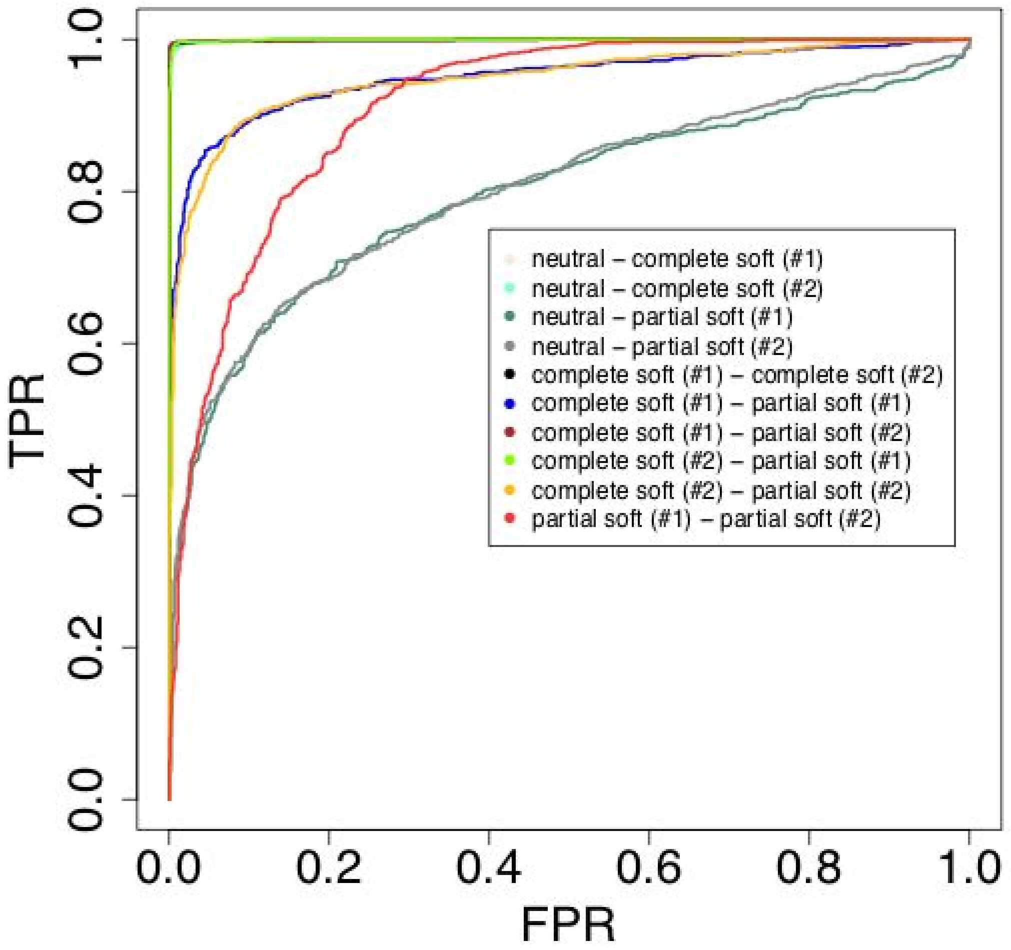


**Supplementary Figure S23:** Receiver Operating Characteristic (ROC) curves for the classification task for complete versus partial soft sweeps in a two-species analysis. The classification task involves five classes: (1) neutral, (2-3) complete soft sweep in species #1 or #2, and (4-5) partial soft sweep in species #1 or #2. A five-way linear SVM classifier was trained using a training set of 40,000 simulated regions of length 50 kb (8,000 per class; see Methods). The classifier was then tested on a separate set comprised of 5,000 regions (1,000 per class). Here, we consider all pairs of classes (see legend), and the ROC curve for each pair records the true positive rate (TPR) as a function of the false positive rate (FPR). See caption of Supplementary Figure S7 for details on the experimental setup.The Area Under Receiver Operating Characteristics (AUROC) for these curves ranged between 0.79 and 0.99.

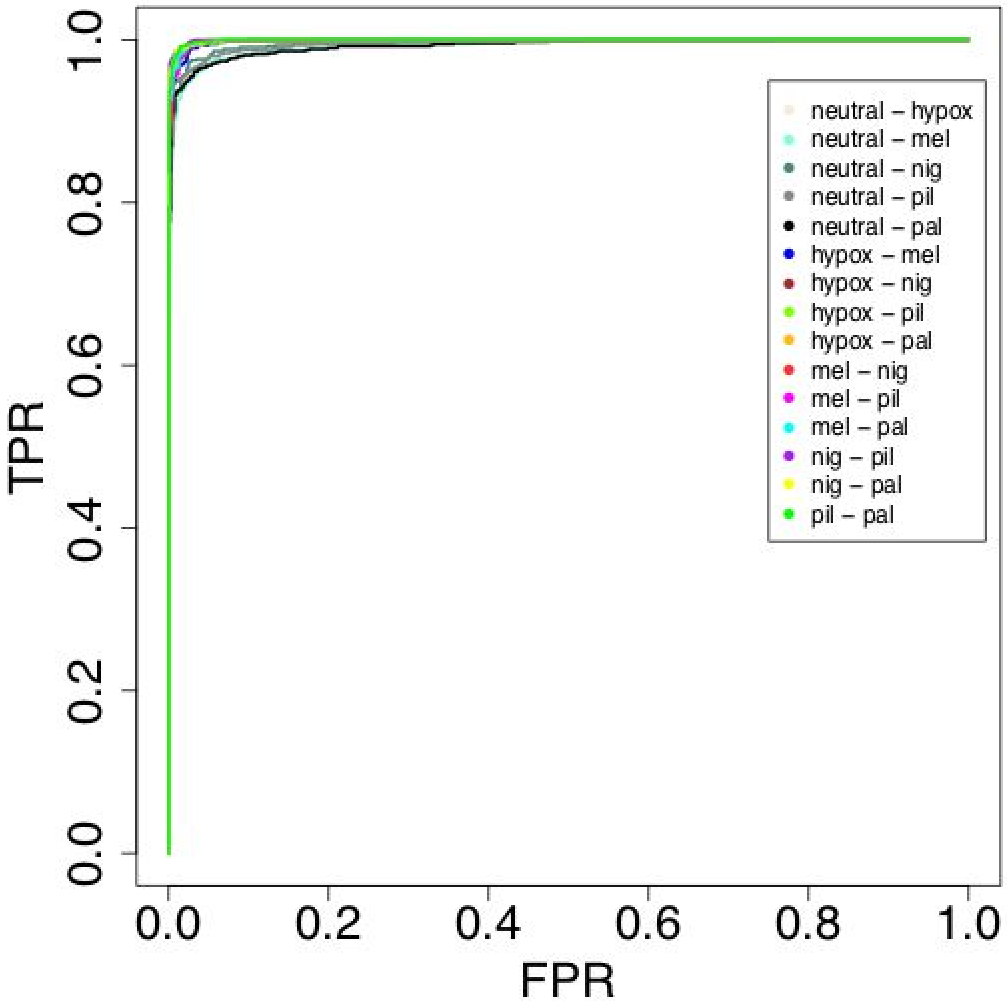


**Supplementary Figure S24:** Receiver Operating Characteristic (ROC) curves for the classification task for neutral regions and regions experiencing soft sweeps in the expanded five-species analysis. The classification task involves six classes: (1) neutral and (2-6) soft sweep in species #1 to #5. A six-way linear SVM classifier was trained using a training set of 48,000 simulated regions of length 50 kb (8,000 per class; see **Methods**). The classifier was then tested on a separate set comprised of 6,000 regions (1,000 per class). Here, we consider all pairs of classes (see legend), and the ROC curve for each pair records the true positive rate (TPR) as a function of the false positive rate (FPR). See caption of Supplementary Figure S7 for details on the experimental setup. The Area Under Receiver Operating Characteristics (AUROC) for these curves ranged between 0.98 and 1.

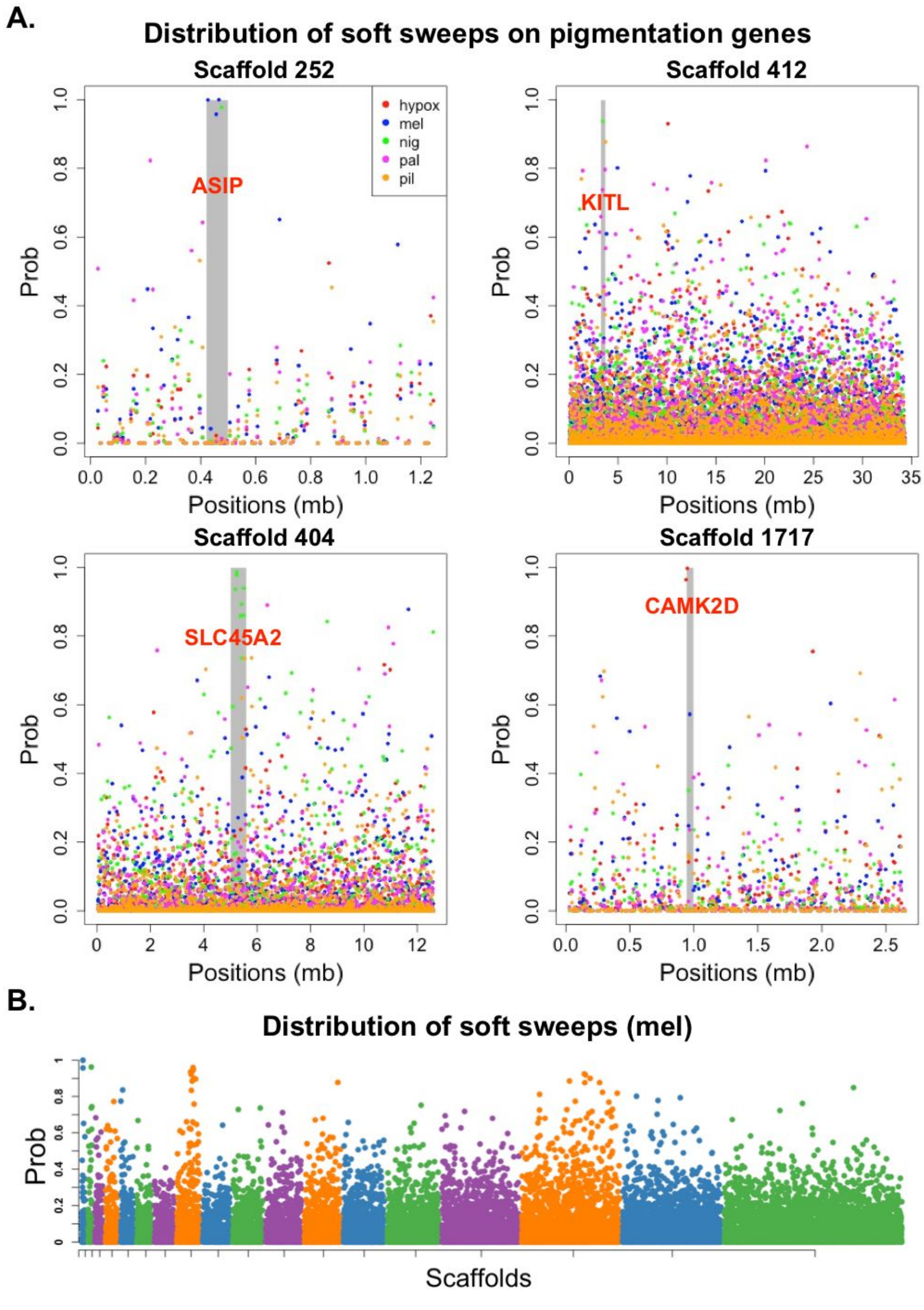


**Supplementary Figure S25:** Manhattan plots showing prediction probabilities for species-specific soft sweeps based on the expanded analysis of all five southern capuchino species. (**A**) Prediction probabilities for all five species are labeled by color (see legend) across four scaffolds harboring top $F_{ST}$ peaks and known pigmentation genes (labeled in *red*). (**B**) Prediction probabilities for species-specific soft sweeps in S. *melanogaster* across all 19 scaffolds harboring $F_{ST}$ peaks. The scaffolds are ordered from smallest to largest.

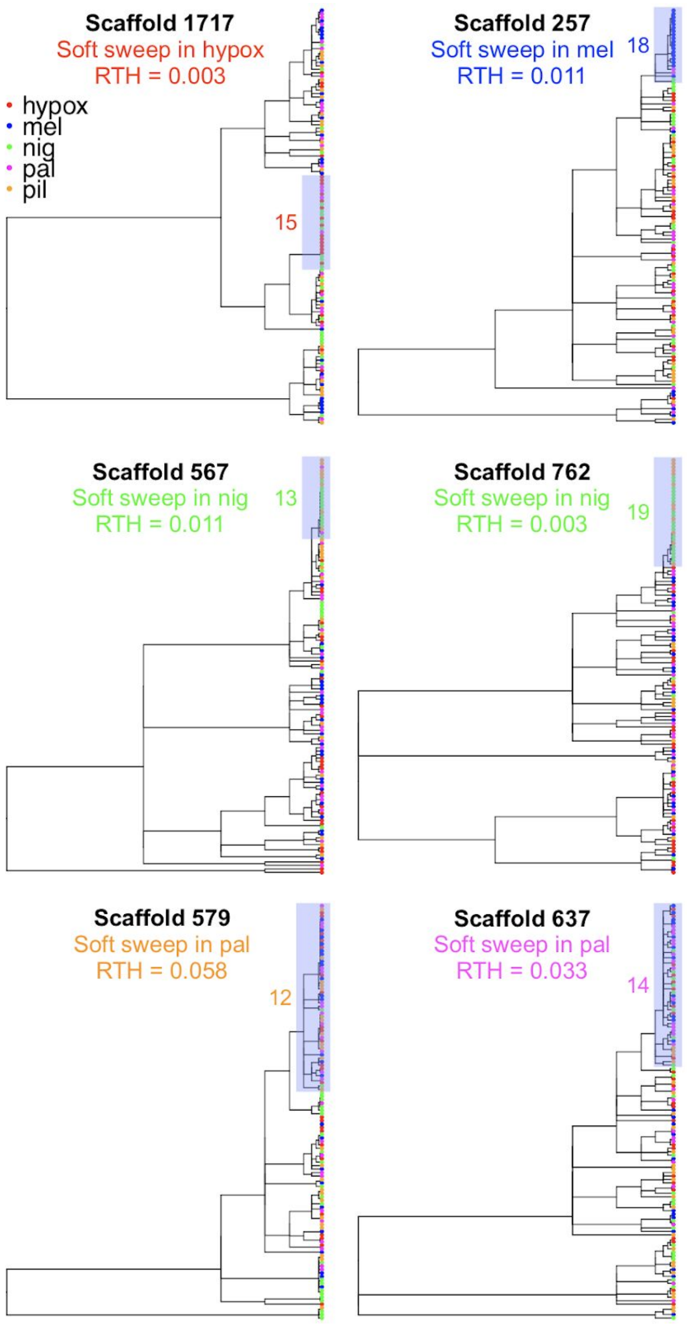


**Supplementary Figure S26:** Example of local trees extracted from 10 kb windows classified as soft sweeps (with prediction probability P>0.9) in the expanded analysis of all five species. In each window, we extracted the local tree inferred by *ARGweaver*, which had the lowest RTH for the target species. The tree leaves are colored based on their species labels (see legend). For each tree, we report the species in which the soft sweep is predicted and the RTH of that species. We also highlight the smallest clade containing at least half the number of samples for the target species which is likely associated with the species-specific sweep, and indicate the number of haploid samples in the clade that belong to the target species.

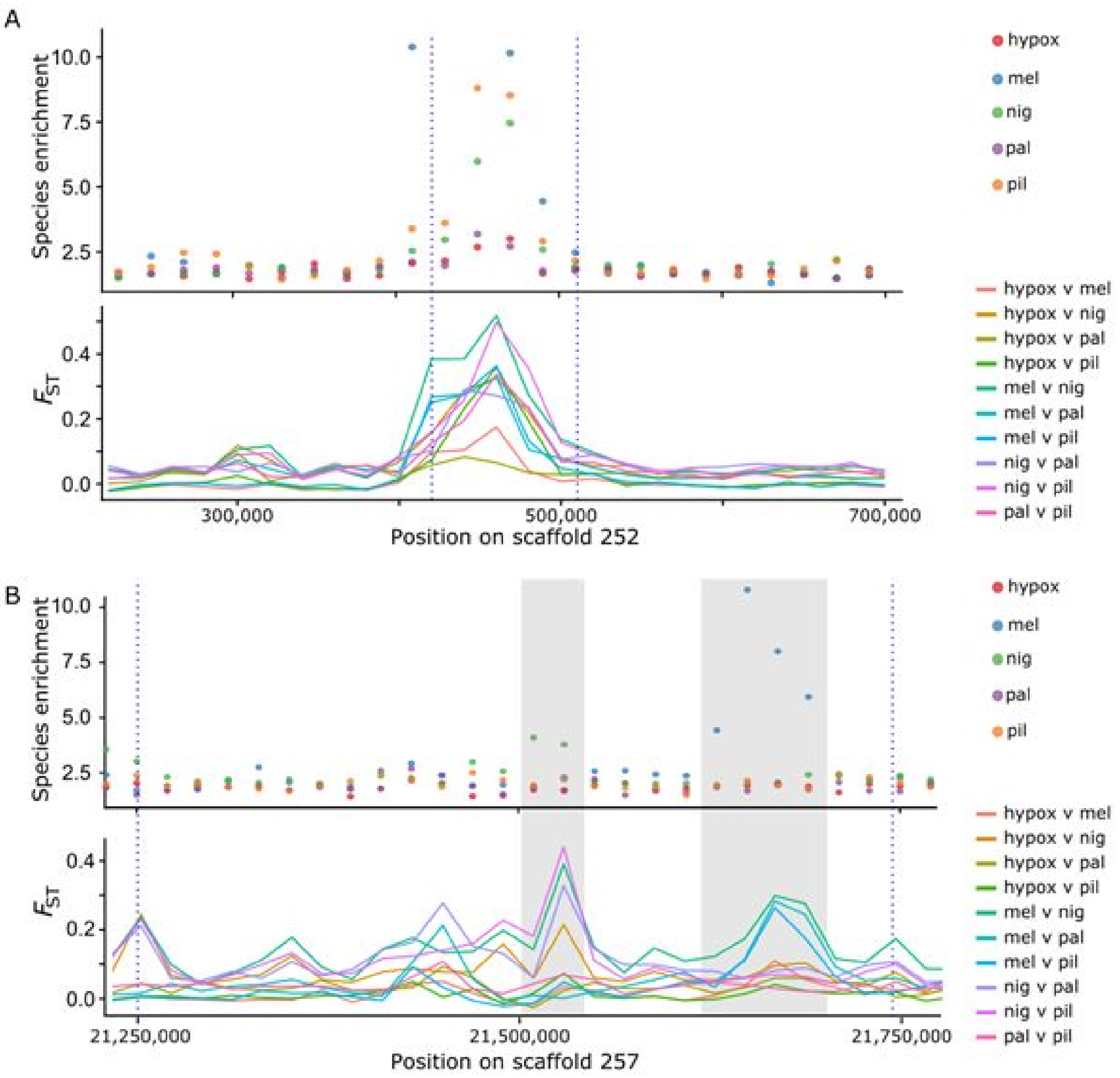


**Supplementary Figure S27:** ARG-based species enrichment scores and sequence-based $F_{ST}$ as measures for species differentiation. Enrichment scores are derived for each species using local trees extracted from the inferred ARG (**Methods**), and $F_{ST}$ values are computed for each species pair using the R package PopGenome V2.7.5. Both scores are averaged across non-overlapping 20 kb windows in the genomic regions containing two $F_{ST}$ peaks (vertical dashed lines) on scaffolds 252 (A) and 257 (B). (A) The $F_{ST}$ peak on scaffold 252 has significantly high enrichment scores for all species other than *S. hypoxantha* (test 1 in **Table 1**), and as a result all $F_{ST}$ values other than $F_{ST}$(hypox,mel) and $F_{ST}$(hypox,pal) exceed the threshold value 0.2, which was used to define peaks. (B) The $F_{ST}$ peak on scaffold 257 (peak 257b in **Supplementary Table S1**) has significantly high enrichment scores for *S. melanogaster* in one region (right vertical gray bar) and elevated enrichment scores for *S. nigrorufa* that do not reach a significance level in another region (left vertical gray bar). In each region we observe elevated values of $F_{ST}$ for pairwise comparisons that involve the enriched species. Note that $F_{ST}$(mel,nig) exceeds 0.2 in both regions. These examples demonstrate that species enrichment scores and $F_{ST}$ provide consistent indications for species differentiation, with species enrichment being easier to interpret due to the fact that it provides a single measure per species.

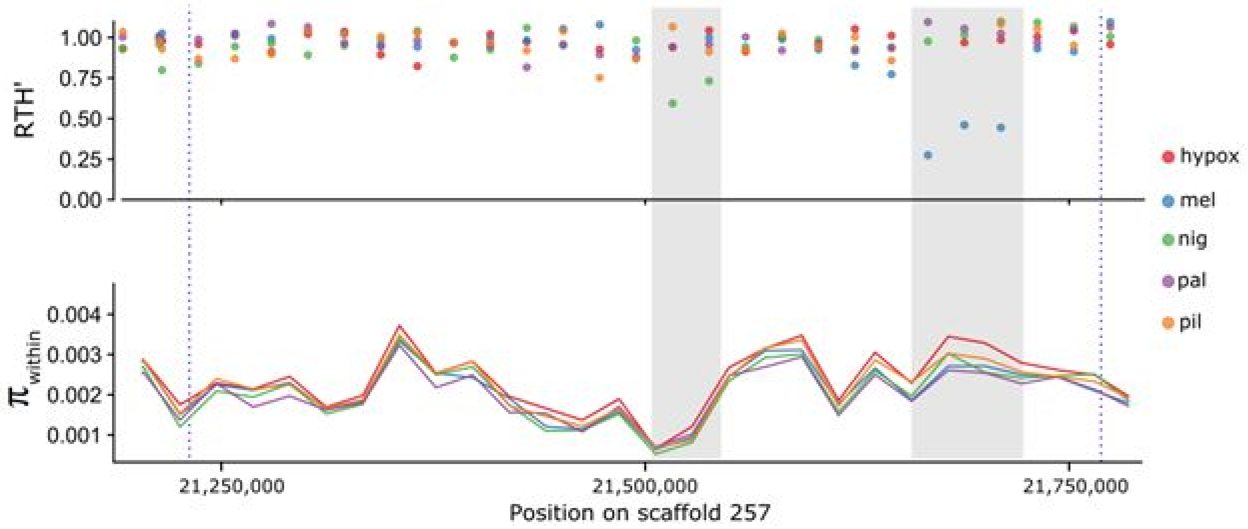


**Supplementary Figure S28:** ARG-based RTH' and sequence-based $\pi_{within}$ (within-species sequence diversity) as potential indicators for species-specific sweeps. RTH' is derived for each species using local trees extracted from the inferred ARG (**Methods**), and $\pi_{within}$ is computed for each species using the R package PopGenome V2.7.5. Both scores are averaged across non-overlapping 20 kb windows in the genomic region containing the $F_{ST}$ peak 257b on scaffolds 257 (vertical dashed lines). RTH' is reduced for *S. melanogaster* and *S. nigrorufa* in two regions that exhibit elevated enrichment scores for these species (vertical gray bars; **Supplementary Figure S27**). In contrast, $\pi_{within}$ is reduced only in one of the two regions (left), and in that region it is similarly reduced for all species. Thus, while we expect $\pi_{within}$ to be reduced in species specific sweeps, it is too noisy to provide a reliable and sensitive method for detection for such sweeps. RTH' is more suitable for this task for three main reasons: (1) The inferred ARG reduces some of the noise caused by missing or faulty genotypes; (2) RTH' is based on the TMRCA of half the species samples, making it sensitive also to partial sweeps; and (3) our normalization scheme for RTH', which divides the TMRCAH of a given species by the general TMRCAH (**Methods**), reduces the genome-wide variance, as seen outside of the two highlighted regions.

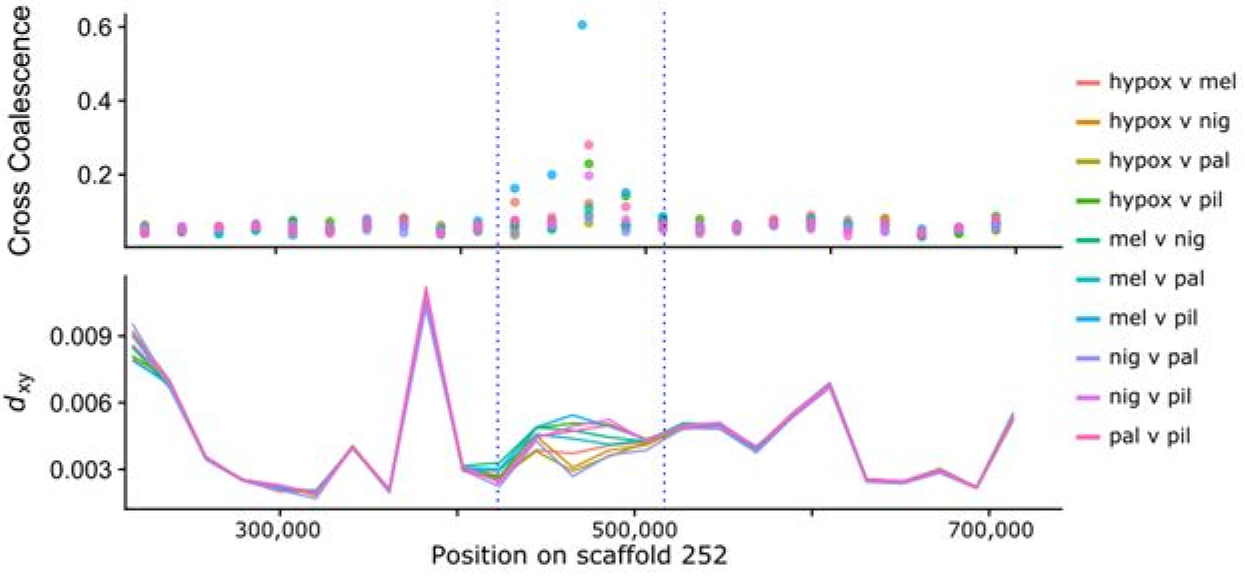


**Supplementary Figure S29:** ARG-based normalized cross coalescence (CC) times and sequence-based $d_{xy}$ as potential indicators for selection against gene flow. Normalized CC times are derived using local trees extracted from the inferred ARG (**Supplementary Text**) and $d_{xy}$ values are computed using the R package PopGenome V2.7.5. Both scores are averaged across non-overlapping 20 kb windows in the genomic region containing the $F_{ST}$ peak on scaffolds 252 (vertical dashed lines). Elevated CC times are observed for *S. pileata* with the other four species, where CC(nig,pil) is the only pair not exceeding the significance threshold (**Supplementary Table S5**). In contrast, $d_{xy}$ shows only a moderate increase in this region for some of these pairwise comparisons, while the regions flanking the peak exhibit higher values. Thus, the signal captured by the sequence-based $d_{xy}$ is much noisier because of high genomic variation. ARG-based CC times are more suitable for this task for three main reasons: (1) The inferred ARG reduces some of the noise caused by missing or faulty genotypes; (2) our measure for CC times considers the most recent cross coalescence events ($n$=10), making it sensitive to subtle changes in the distribution of CC times caused by selection against gene flow; and (3) our normalization scheme, which divides the CC times by the general TMRCAH (**Methods**), reduces the genome-wide variance, as seen outside of the $F_{ST}$ peak.

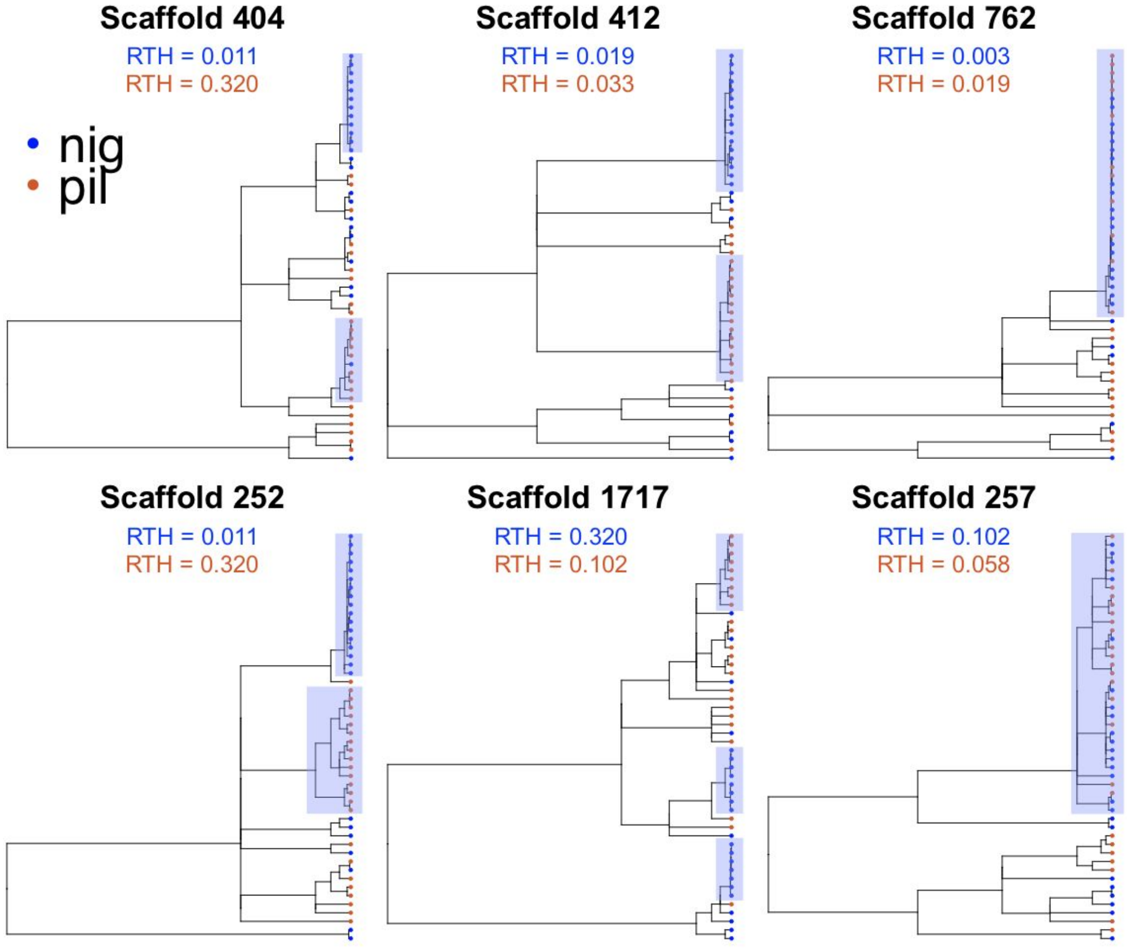


**Supplementary Figure S30:** Example of local trees extracted from 10 kb windows classified with probability P>0.95 as parallel soft sweeps in *S. nigrorufa* and *S. pileata* (see Supplementary Table S16 for more details on the classifier used for this task). The local trees were inferred by *ARGweaver* and pruned to show only the haploid samples from *S. nigrorufa* (blue) and *S. pileata* (orange). The RTH statistic is reported for each species in each local tree. The four trees extracted from scaffolds 404, 252, 412, and 1717 show multiple young and moderately large clades, each enriched for one of the two species, as would be expected in the scenario of a parallel sweep. On the other hand, the trees extracted from scaffolds 762 and 257 have a young and very large clade that contains individuals from both species. Such a “shared sweep” could be ancestral or a result of adaptive introgression. For example, the shared sweep on scaffold 257 appears to have a more ancient origin, suggesting that the sweep may have started before the two species diverged. However, the sweep on scaffold 762 appears to have a very recent origin, suggesting that it originated in one of the species after the species split, and then introduced into the other species as a result of adaptive introgression (see also Supplementary Figure S31).

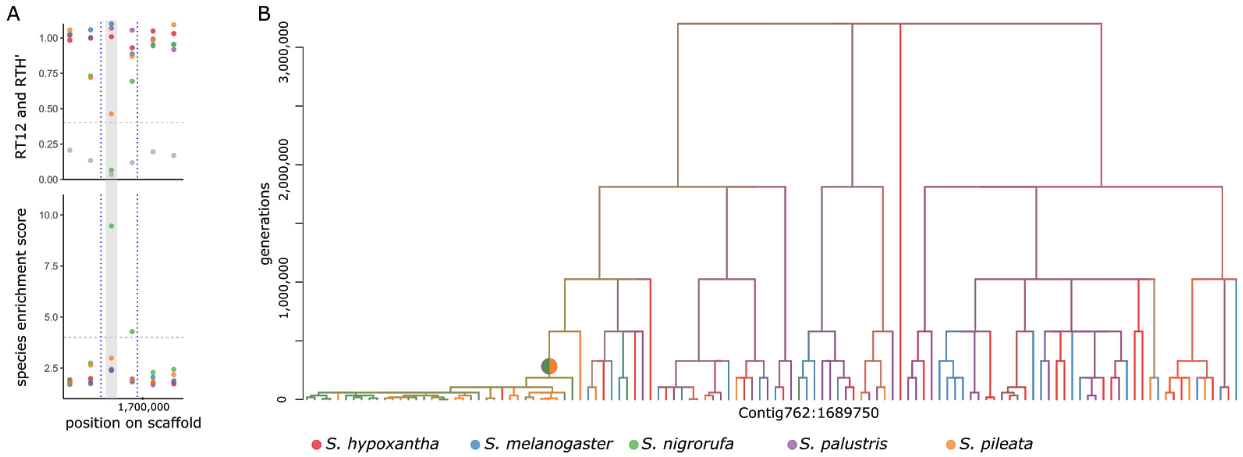


**Supplementary Figure S31:** Genealogical statistics and a representative tree for the $F_{ST}$ peak on scaffold 762. **A)** RTH', RT12, and species enrichment scores averaged across non-overlapping 20 kb segments in the genomic region containing the divergence peak. **B)** A representative tree extracted from the inferred ARG inside the 20 kb segment with the highest enrichment scores within the peak (gray vertical bar in panel A). See caption of **Supplementary Figure S5** for complete specification of color code and vertical and horizontal dashed lines in panel A. In this peak we observe a significantly high enrichment score for *S. nigrorufa*, and significantly low RTH' for *S. nigrorufa* and *S. pileata* (**Table 1**, and **Supplementary Tables S2** and **S3**). A young clade enriched for both species is indicated by a colored circle at its root. This clade is roughly 250,000 generations old and contains 35 samples, out of which 21 are from *S. nigrorufa* and 13 samples from *S. pileata*, suggesting a recent selective sweep shared by both species, possibly due to adaptive introgression (see also **Supplementary Figure S30**). In this peak we also observe elevated inter-species cross coalescence times between either *S. nigrorufa* or *S. pileata* and the other three species (**Supplementary Table S5**). However, since there is no evidence for a deep enriched clade in this peak, we attribute this elevation to the fact that the recent sweep experienced by *S. nigrorufa* and *S. pileata* reduced the number of lineages from these two species that were free to coalesce with lineages from other species before the time that the sweep initiated (**Supplementary Text**).

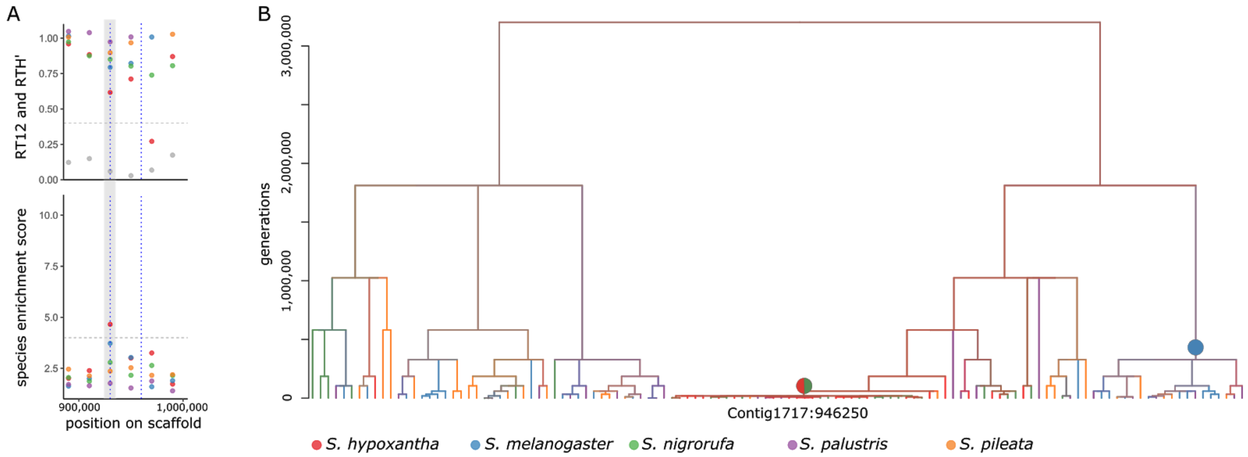


**Supplementary Figure S32:** Genealogical statistics and a representative tree for the $F_{ST}$ peak on scaffold 1717. **A)** RTH', RT12, and species enrichment scores averaged across non-overlapping 20 kb segments in the genomic region containing the divergence peak. **B)** A representative tree extracted from the inferred ARG inside the 20 kb segment with the highest enrichment scores within the peak (gray vertical bar in panel A). See caption of **Supplementary Figure S5** for complete specification of color code and vertical and horizontal dashed lines in panel A. In this peak we observe significantly high enrichment scores for *S. melanogaster* and *S. hypoxantha* (**Table 1**, and **Supplementary Table S2**), and a significantly low RTH' for *S. hypoxantha* in the 20 kb segment immediately adjacent to the peak boundaries. Two clades, one enriched for *S. melanogaster* and one enriched for *S. hypoxantha* and *S. nigrorufa*, are indicated by colored circles at their roots. The second clade is young (roughly 50,000 generations), and out of the 33 samples it contains, 17 are from *S. hypoxantha* and 12 are from *S. nigrorufa*. This suggests a recent selective sweep shared by both species in this region, possibly due to adaptive introgression. The other clade is considerably deeper (roughly 500,000 generations), but only marginally enriched for *S. melanogaster*, with nine out of its 20 samples belonging to that species. In this peak we also observe elevated inter-species cross coalescence times between *S. melanogaster* and *S. nigrorufa* or *S. hypoxantha*, which could be a result of this relatively deep and enriched clade (**Supplementary Table S5**). However, this elevation could also be a result of the selective sweep experienced by *S. hypoxantha* and *S. nigrorufa,* which reduced the number of lineages from these two species that were free to coalesce with lineages from *S. melanogaster* before the time that the sweep initiated. This, together with the fact that enrichment for *S. melanogaster* is marginally significant, reduces the likelihood of a significant contribution of selection against gene flow on divergence in this region.

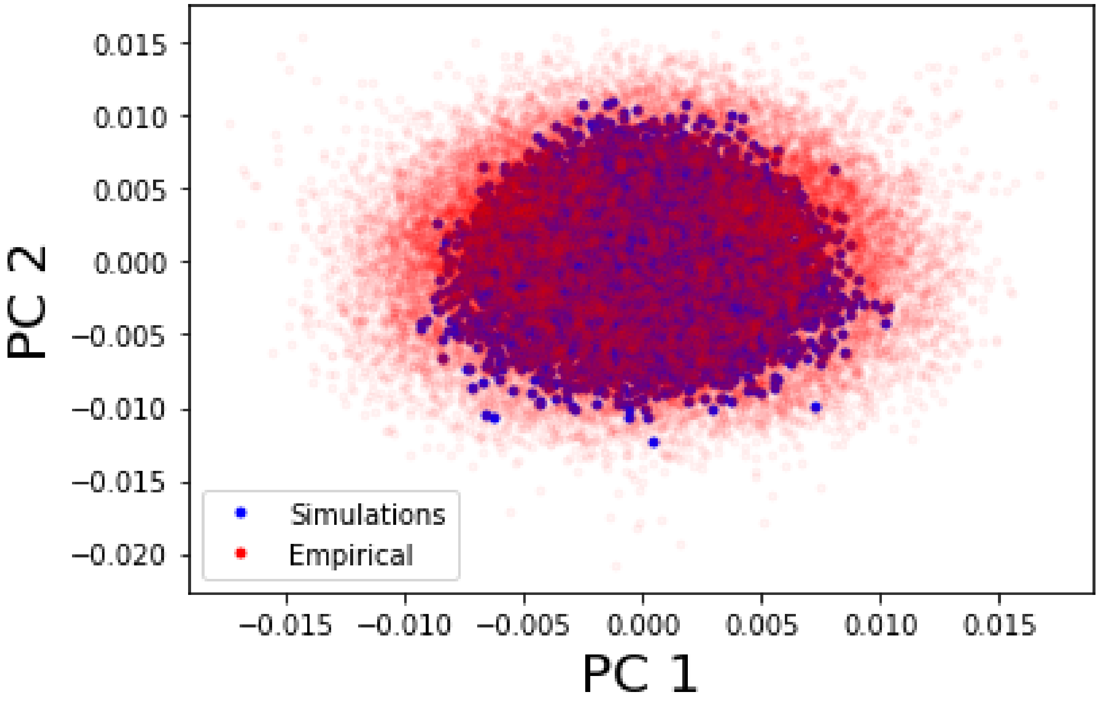


**Supplementary Figure S33:** Principal Component Analysis (PCA) applied on simulations based on the inferred Capuchino demographic model from RAD-seq data. Results suggest that the simulations fit the genomic data. PCA provides a reasonable check that our simulations generalize well to represent the empirical data. The x-axis represents the top principal component (i.e. PC with the largest eigenvalue) while the y-axis represents the second top principal component (i.e. PC with the second largest eigenvalue). We applied PCA to the summary statistics extracted from both the empirical data and the simulations based on the demographic model inferred from RAD-seq data. We found that the top two principal components for the two datasets largely overlap, suggesting that the inferred demographic model fits the genomic data reasonably well.

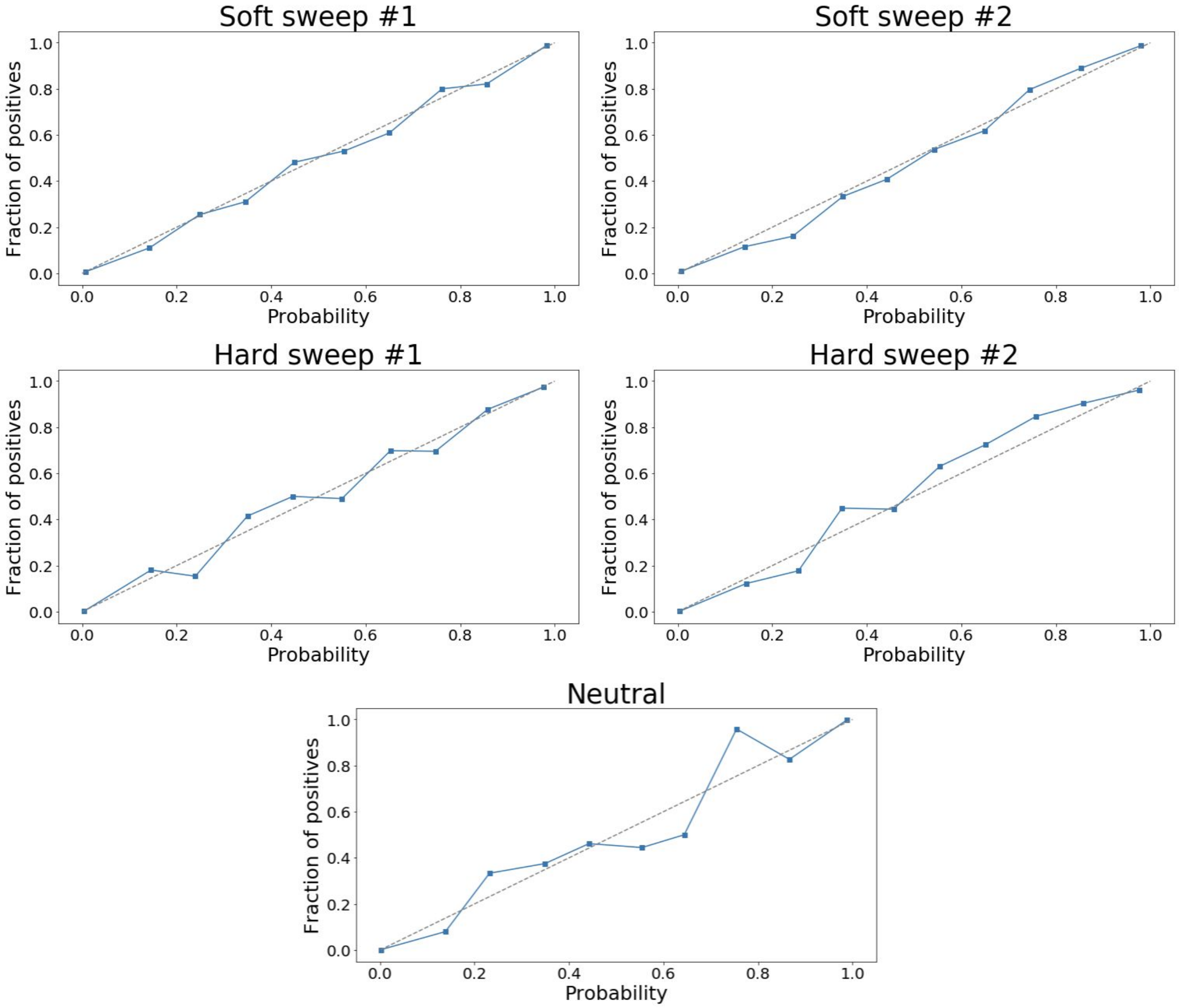


**Supplementary Figure S34:** Calibration curves for the classification task for soft versus hard sweeps in a two-species analysis**.** We generated these curves to assess how well calibrated are the probabilities generated by the classifier. The classification task involves five classes: (1) neutral, (2-3) soft sweep in species #1 or #2, and (4-5) hard sweep in species #1 or #2. A five-way linear SVM classifier was trained using a training set of 40,000 simulated regions of length 50 kb (8,000 per class; see **Methods**). The classifier was then tested on a separate set comprised of 5,000 regions (1,000 per class). For each subplot, we took a one-versus-rest approach (i.e. soft sweep in species #1 versus all). The *x*-axis reports the mean predicted probability in each bin, while the *y*-axis reports the fraction of samples in the positive class in each bin. We discretize the probabilities in the [0, 1] interval into 10 bins. The points on these curves appear to fall on the main diagonal, suggesting that these probabilities are well calibrated.
